# ALT Neuroblastoma Chemoresistance due to ATM Activation by Telomere Dysfunction is Reversible with the ATM Inhibitor AZD0156

**DOI:** 10.1101/2021.04.06.438692

**Authors:** Balakrishna Koneru, Ahsan Farooqi, Thinhh H. Nguyen, Wan Hsi Chen, Ashly Hindle, Cody Eslinger, Monish Ram Makena, Trevor A. Burrow, Joanne Wilson, Aaron Smith, Venkatesh Pilla Reddy, Elaine Cadogan, Stephen T. Durant, C. Patrick Reynolds

**Author notes:** Correspondence: C. Patrick Reynolds, MD PhD, Address: Cancer Center, Texas Tech University Health Sciences Center School of Medicine, 3601 4th Street, Mail Stop 9445 Lubbock, Texas 79430-6450.

## Abstract

Cancers overcome replicative immortality by activating either telomerase or an alternative lengthening of telomeres (ALT) mechanism. ALT occurs in ∼ 25% of high-risk neuroblastomas and relapse or progression in ALT neuroblastoma patients during or after front-line therapy is frequent and almost uniformly fatal. Temozolomide + irinotecan is commonly used as salvage therapy for neuroblastoma. Patient-derived cell-lines and xenografts established from relapsed ALT neuroblastoma patients demonstrated *de novo* resistance to temozolomide + irinotecan (as SN-38 *in vitro, P*<0.05) and *in vivo* (mouse event-free survival (EFS) *P*<0.0001) relative to telomerase-positive neuroblastomas. We observed that ALT neuroblastoma cells manifest constitutive ATM kinase activation due to spontaneous telomere dysfunction while telomerase-positive tumors lacked constitutive ATM activation or spontaneous telomere DNA damage. We demonstrated that induction of telomere dysfunction resulted in ATM activation that in turn conferred resistance to temozolomide + SN-38 (4.2 fold-change in IC50, *P*<0.001). ATM kinase shRNA knock-down or inhibition using a clinical-stage small molecule inhibitor (AZD0156) reversed resistance to temozolomide + irinotecan in ALT neuroblastoma cell-lines *in vitro* (*P*<0.001) and in 4 ALT xenografts *in vivo* (EFS *P*<0.0001). AZD0156 showed modest to no enhancement of temozolomide + irinotecan activity in telomerase-positive neuroblastoma cell lines and xenografts. ATR inhibition using AZD6738 did not enhance temozolomide + SN-38 activity in ALT neuroblastoma cell lines. Thus, resistance to chemotherapy in ALT neuroblastoma occurs via ATM kinase activation and was reversed with the ATM inhibitor AZD0156. Combining AZD0156 with temozolomide + irinotecan warrants clinical testing in neuroblastoma.

**One Statement Summary:** ATM activation at telomeres confers resistance to DNA damaging chemotherapy in ALT neuroblastoma that was reversed with ATM knockdown or inhibition.

## Introduction

Unlimited proliferation of cancer cells requires that they maintain telomeres (*1*). Most cancers maintain their telomeres by activating telomerase (*2*). Alternatively, some cancers use the recombination-mediated alternative lengthening of telomeres (ALT) mechanism (*3*), which is more prevalent in tumors of mesenchymal and neuro-epithelial origin, including osteosarcoma, pancreatic neuroendocrine tumors, gliomas, and neuroblastoma (*4*). ALT is characterized by a high frequency of telomere-sister chromatid exchanges (T-SCEs) (*5, 6*), heterogeneous telomere length (*7*), presence of ALT-associated promyelocytic leukemia (PML) nuclear bodies (APBs) (*8*), and presence of extrachromosomal telomeric DNA repeats in the form of partially double-stranded circles, termed C-circles (*9*). Although the molecular mechanism of how ALT telomere maintenance occurs is not yet clear, previous studies suggest that ALT cells extend telomeres via a complex break-induced replication pathway initiated by either DNA damage at telomeres or telomere replication stress (*10–12*). A previous study showed that ALT cells are hypersensitive to ATR inhibitors due to telomere replication stress (*13*). However, several subsequent studies refuted these findings, suggesting that ALT cancer cells do not show general hypersensitivity to ATR inhibitors, including neuroblastoma (*14, 15*).

With the recent advances in understanding ALT-mediated telomere maintenance, several other therapeutic strategies have been proposed to exploit the ALT associated vulnerabilities of ALT tumors (*16–19*), but none of these ALT specific therapeutic strategies have yet translated into the clinic. Also, long-term growth of tumor cells with no apparent telomere maintenance has been reported, suggesting that direct targeting of telomere maintenance mechanisms as a therapeutic strategy may encounter resistance mechanisms (*20, 21*).

Neuroblastoma is a pediatric solid tumor of the peripheral sympathetic nervous system that is responsible for 11% of pediatric cancer deaths (*22, 23*). Neuroblastoma patients stratified as high- risk continue to have poor outcomes, with an overall survival rate of ∼ 50% with current comprehensive therapy (*24–26*). Using next generation sequencing and molecular approaches, we and several other groups observed that most common genomic alterations in high-risk neuroblastoma (*MYCN* amplification, *TERT* rearrangements, and *ATRX* genomic alterations) converge on activating telomere maintenance mechanisms (*20, 27–29*). We have recently shown that telomere maintenance mechanisms (TMM) define clinical outcome in high-risk neuroblastoma, irrespective of currently employed risk factors, suggesting that TMM are pivotal for neuroblastoma pathogenesis and may potentially provide therapeutic targets (*29*).

Previous studies, using less-than-specific means of identifying ALT, suggest that ALT neuroblastomas have distinct clinical behavior such as older age at diagnosis, protracted disease progression, and poor response to chemotherapy (*30–32*). Employing more specific ALT markers, especially the robust telomeric DNA C-circle assay, we showed that ALT activation occurs in ∼25- 30% of high-risk neuroblastomas with or without *ATRX* genomic alterations (*29*). Patients with ALT neuroblastoma have very poor long-term survival and are often non-salvageable following progression or relapse (*29*). Similar to ALT neuroblastoma, ALT pancreatic neuroendocrine tumors, gliomas, and soft tissue sarcomas are known to have distinct clinical behavior and outcome relative to non-ALT tumors (*33–37*), suggesting that ALT mediated telomere maintenance may impact cancer cell growth and the response to treatment.

Due to a lack of relevant tractable *in vitro* and *in vivo* ALT neuroblastoma models, development of therapeutic approaches against ALT neuroblastoma has remained challenging since the first report of ALT in neuroblastoma (*38*). In the present study, using patient-derived ALT neuroblastoma cell lines and patient derived xenografts (PDX) established from relapsed neuroblastoma patients (*29, 39*), we demonstrated that ALT neuroblastoma is associated with chemo-resistance to DNA damaging agents due to constitutive activation of ataxia-telangiectasia mutated (ATM) kinase signaling as a result of constitutive telomere dysfunction. ATM is known to play a critical role in detection and repair of double-strand breaks (DSBs) resulting from endogenous genomic stress or due to exogenous irradiation or DNA damaging chemotherapy (*40*). We showed that the chemo-resistant phenotype can be reversed in ALT neuroblastoma by inhibiting ATM kinase using AZD0156, a clinical-stage small molecule ATM inhibitor, *in vitro* and *in vivo*. Together, these data provide a potential therapeutic strategy for treating refractory ALT neuroblastoma patients.

## Results

### ALT neuroblastoma models are associated with resistance to DNA-damaging drugs *in vitro* and *in vivo*

To investigate differences in sensitivity to chemotherapy among neuroblastoma cell lines based on ALT status, we compared cytotoxicity profiles of ALT (n=4) vs non-ALT (n=100) patient-derived neuroblastoma cell lines (Supplementary Table. S1) treated with drugs commonly used to treat neuroblastoma patients: topoisomerase inhibitors 7-Ethyl-10-hydroxycamptothecin (SN-38; an active metabolite of irinotecan), topotecan (TOPO), and etoposide, alkylating agent’s cyclophosphamide (as 4-hydroperoxy cyclophosphamide (4-HC) *in vitro*) and melphalan, and carboplatin. ALT neuroblastoma cell lines showed significantly higher resistance to these DNA damaging agents when compared to non-ALT neuroblastoma cell lines (Figure. 1A, Wilcoxon, *P*<0.05). Additionally, ALT cell lines (n=4) were also more resistant than non-ALT cell lines (n=36) to drug combinations (4-HC + TOPO, or temozolomide (TMZ) + SN-38) that are commonly used as an induction or salvage therapy for relapsed neuroblastoma patients (Figure. 1B; Wilcoxon, *P*<0.05). All ALT cell lines were DNA C-circle positive, had ALT associated PML bodies and are *TERT* mRNA (enzymatic component of telomerase) non-expressing while most non-ALT lines were *TERT*-expressing (Table S1).

**Figure 1.**
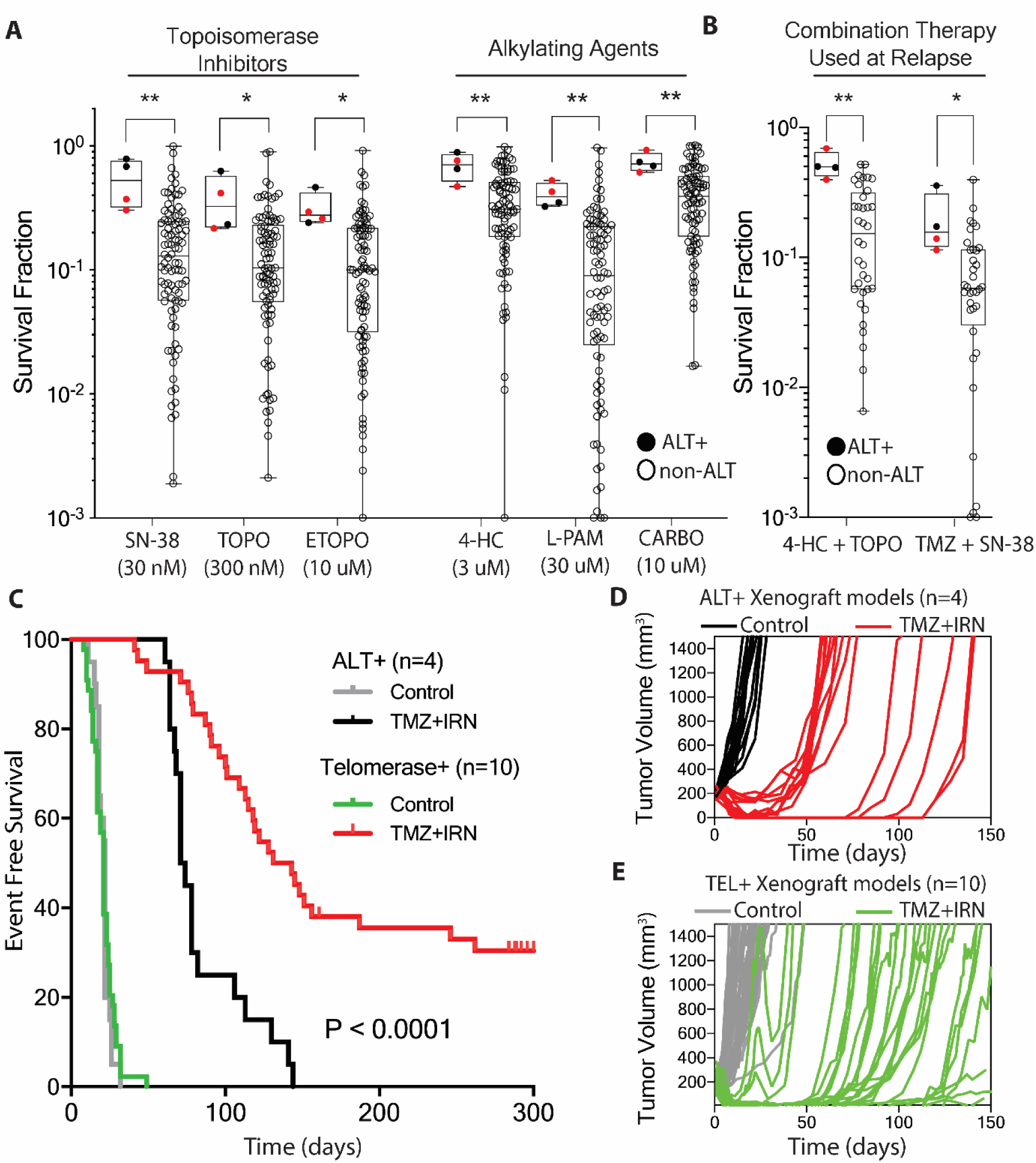
ALT neuroblastoma models manifested chemo-resistance to DNA damaging agents *in vitro* and *in vivo*. (A) Box plot compares survival fraction of ALT (n=4) versus non-ALT (n=100) neuroblastoma cell lines treated with clinically achievable doses of 7-ethyl-10- hydroxycamptothecin (SN-38; an active metabolite of irinotecan), topotecan (TOPO), etoposide (ETOPO), 4-hydroperoxy cyclophosphamide (4-HC; an active metabolite of cyclophosphamide), melphalan (L-PAM), and carboplatin (CARBO. (B) Box plot compares survival fraction of ALT (n=4) versus telomerase-positive (n=36) cell lines in response to combination treatment with either 4-HC (1 μM) + TOPO (100 nM) or temozolomide (TMZ; at 30 μM) + SN-38 (3 nM). Survival fraction was determined using the DIMSCAN cytotoxicity assay. ALT positive cell lines COG-N- 512 and COG-N-515 that were established from same patient are highlighted as red circles in A and B. (C) Kaplan–Meier event-free survival (EFS) curves for ALT (red lines; ALT CDXs: COG- N-515m, CHLA-90m, SK-N-FIm; ALT PDX: COG-N-625x) versus telomerase-positive (green lines; Telomerase-positive CDX: CHLA-79m, CHLA-119m, FELIXm, telomerase-positive PDXs: COG-N-452x, COG-N-519x, COG-N-632x, COG-N-421x, COG-N-470x, COG-N-561x, COG- N-564x) xenograft models in response to treatment with 2 cycles of TMZ (25 mg/kg) + irinotecan (IRN; 7.5 mg/kg) on Days 1-5 in a 21 day cycle. Control mice were administered sterile carrier solutions. EFS was calculated as the time to reach the tumor volume of 1500 mm^3^ from initiation of treatment with vehicle or TMZ+IRN or death from any cause. Each xenograft model had 3-5 mice in both control and treatment groups. (D) Tumor volume of ALT, (E) and telomerase-positive xenograft models treated with vehicle or TMZ+IRN, in same mice as shown in C. Survival fraction between ALT versus non-ALT cell lines was compared using Wilcoxon-rank sum test. Kaplan- Meier survival curves were compared using the log-rank test. *: *P*<0.05, **: *P*<0.01.

Loss of p53 function has been shown to be associated with chemo-resistance in neuroblastoma (*41*). Our group previously reported *TP53* mutations in 2 neuroblastoma cell lines (CHLA-90 and SK-N-FI) (*39*). We established ALT neuroblastoma cell lines COG-N-512 and COG-N-515 from different biopsies obtained from the same patient; both cell lines match by short tandem repeat assay and are wild-type for *TP53* using whole exome and Sanger sequencing and wild-type for p53-pathway genes (Table S2). However, COG-N-515 failed to induce p53 and p21 protein expression upon irradiation (Supplementary Figure. S1A & S1B), indicating that this cell line is non-functional for p53. As p53 loss-of-function can induce chemo-resistance in neuroblastoma (*41*), we determined if the ALT cell lines have cytotoxicity profiles similar to that of p53 non-functional telomerase-positive cell lines. ALT neuroblastoma cell lines showed hyper-resistance to topoisomerase inhibitors (SN-38, etoposide, and topotecan; *P*<0.05), alkylating agents (4-HC, melphalan), and carboplatin; *P*<0.05, and to DNA-damaging drug combinations (TMZ+SN-38 and 4-HC+TOPO; *P*<0.05) when compared to p53 non-functional telomerase-positive cell lines (Supplementary Figure. S1C).

To determine if chemo-resistance observed *in vitro* can be recapitulated *in vivo,* we evaluated the *in vivo* response to TMZ + irinotecan (IRN) in an ALT patient-derived xenograft (PDX) (COG-N-625x) and cell line-derived xenografts (CDX) (CHLA-90m, COG-N-515m and SK-N-FIm) compared to 10 randomly chosen telomerase-positive PDXs (COG-N-519x, COG-N- 421x, COG-N-470x, COG-N-561x, COG-N-564x COG-N-452x, COG-N-623x) and CDX (CHLA-119m, CHLA-79m and Felix-m) models (Table S1), all of which were established from neuroblastoma patients after therapy at time of disease progression or at time of death from progressive disease. Mice with ALT xenografts treated with TMZ+IRN had a substantially lower event-free survival (EFS) when compared to telomerase-positive xenograft models (log-rank test: *P*<0.0001; Figure. 1C). The majority of mice in ALT xenograft models (3 of 4 ALT models; CHLA-90m, SK-N-FIm, COG-N-625x) treated with TMZ+IRN either progressed rapidly after responding or had minimal to no response (Figure. 1D) compared to complete responses in 8/10 telomerase-positive xenograft models (Figure 1E) following treatment with TMZ+IRN (Fisher exact test: mice in complete response for ALT vs telomerase-positive models: *P*<0.0001). Telomerase-positive PDXs COG-N-564x and COG-N-519x were the only telomerase-positive models that did not achieve complete responses when treated with TMZ+SN38. Although complete responses were observed in most telomerase-positive xenograft models, tumors progressively grew after completion of treatment cycles.

### ALT neuroblastoma was associated with constitutive DNA damage signaling and ATM activation

To elucidate mechanisms of the high-level chemo-resistance observed in ALT neuroblastoma, we compared gene expression profiles of ALT (n=3) versus randomly selected telomerase-positive non-*MYCN*-amplified (n=8) neuroblastoma cell lines (all established from patients with progressive disease) using RNA sequencing (Table S1). *MYCN*-amplified cell lines were excluded from global gene expression analysis as ALT occurs mutually exclusive of *MYCN* amplification in neuroblastoma and *MYCN* amplified tumors have chromosomal alterations distinct from non-*MYCN* amplified tumors (*29, 39*). Gene set enrichment analysis (GSEA) of Biocarta gene sets indicated that ATM and ATR-BRCA signaling pathways were enriched in ALT neuroblastoma cell lines (Figure. 2A and 2B; Table S3). Both ATM and ATR-BRCA signaling pathways are known to be key players in DNA damage repair following genomic stress, such as double strand breaks (DSBs). As DNA damage signaling appears to be constitutively active in ALT cell lines, we assessed basal levels of DNA damage by immunofluorescent staining for 53BP1 (a DSBs marker) in ALT (n=3) versus telomerase-positive (n=8) neuroblastoma cell lines; telomerase-positive cell lines used for 53BP1 analysis included 4 *MYCN* non-amplified cell lines used for GSEA and 4 randomly selected *MYCN* amplified cell lines (Table S1). ALT cell lines had substantially higher levels of 53BP1 foci compared to telomerase-positive cell lines irrespective of *MYCN* amplification (Figure. 2C; *P*<0.001). In line with the notion that ALT is associated with chronic endogenous DNA damage, higher tail moment by comet assay (Supplementary Figure. S2A & S2B; *P* < 0.001) and higher levels of γ-H2AX (marker for DNA-damage; Supplementary Figure. S2C; *P*<0.01) were observed in ALT cell lines compared to 2 telomerase-positive p53 non-functional cell lines: (SK-N-BE(2) a *MYCN* amplified line and CHLA-171 a non-*MYCN* amplified) line.

**Figure 2.**
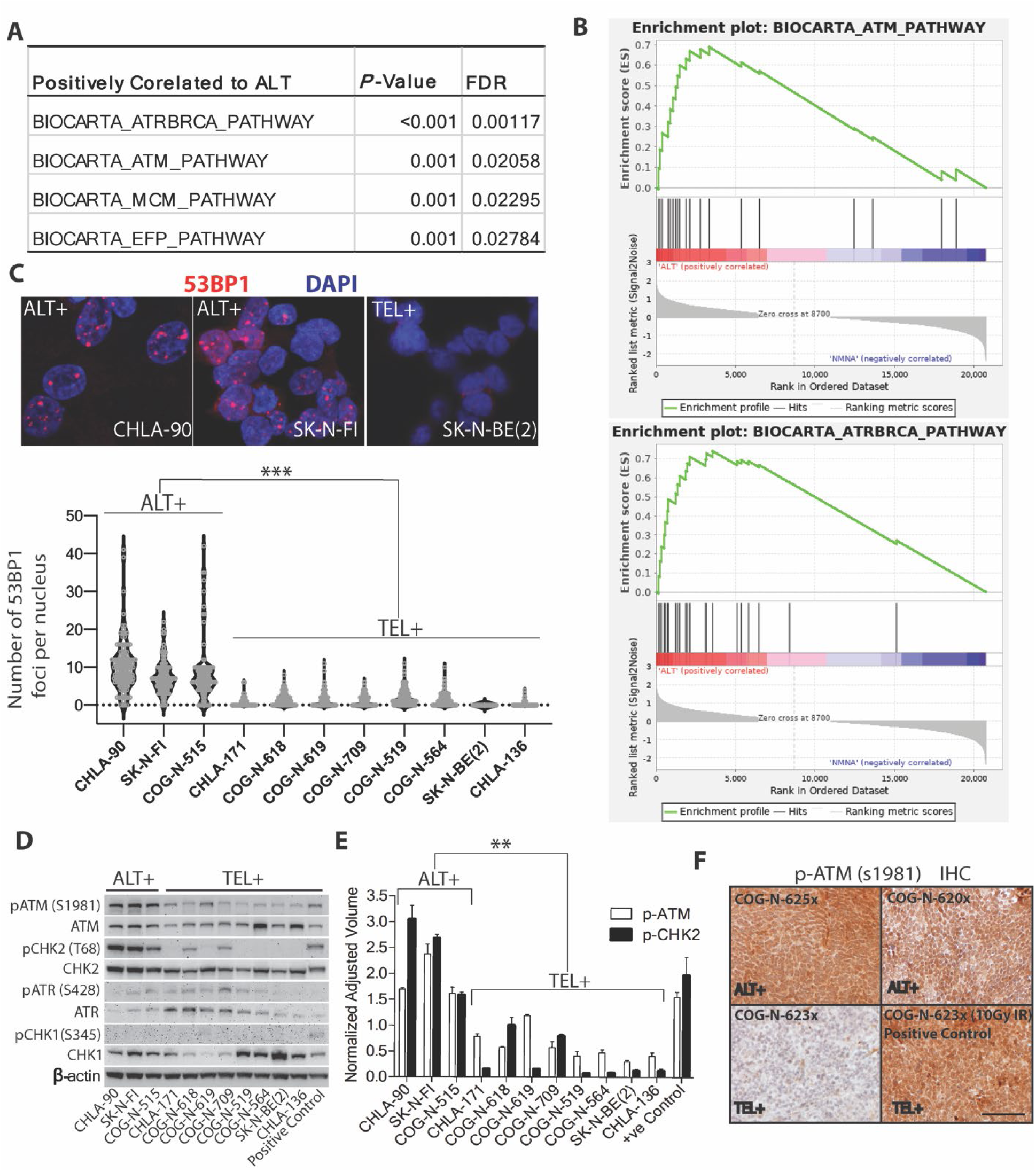
ALT neuroblastoma is associated with constitutive DNA-damage signaling and ATM activation. (A) Table lists significantly upregulated Biocarta gene sets comparing ALT (n=3) versus telomerase-positive non-*MYCN* amplified (n=8) neuroblastoma patient-derived cell lines using Gene Set Enrichment Analysis GSEA. Biocarta gene sets with *P*-value < 0.05 and False Discovery Rate (FDR) < 5% are listed. (B) Enrichment plot showing positive enrichment for ATM and ATR-BRCA pathway are shown. Enrichment plots were derived from GSEA results in gene pattern environment. (C) Immunofluorescence (IF) staining for 53BP1 (a double strand break marker) in ALT versus telomerase-positive neuroblastoma cell lines. Top panel displays representative images of IF staining for 53BP1 (red) in ALT cell lines CHLA-90 and SK-N-FI when compared to telomerase-positive cell line SK-N-BE(2). Nuclei were stained with DAPI (blue). Bottom panel displays a violin plot for 53BP1 foci quantification in ALT (CHLA-90, SK-N-FI, and COG-N-515) versus telomerase-positive (n=8; Non-*MYCN* amplified: CHLA-171, COG-N-618, COG-N-619 and COG-N-709; *MYCN* amplified COG-N-519, COG-N-564, SK-N-BE(2) and CHLA-136) cell lines. A minimum of 100 nuclei per cell line were analyzed for quantification of 53BP1 foci. (D) Immunoblotting for pATM (S1981), ATM, pCHK2 (T68), CHK2, pATR (S345), ATR, pCHK1 (S345), CHK1 and β-actin in same cell lines as shown in C. CHLA-136 twelve hours post-irradiation with x-rays (10 Grays) served as a positive control. (E) Quantification of pATM (S1981) and pCHK2 (T68) expression for the immunoblot in D and its replicates. The bars represent means with SDs from three experimental replicates. (F) Representative images of immunohistochemistry for pATM (s1981) in ALT PDXs (COG-N-625x and COG-N-620x) versus telomerase-positive PDX (COG-N-623x). COG-N-623x mice irradiated with 10 grays of x-ray served as a positive control for ATM activation. Scale bar is a reference for 50 µm in distance. Statistical analysis for C and E was performed by Wilcoxon-rank sum test. ***: *P*<0.001, **: *P*<0.01.

Immunoblotting for ATM and ATR activation demonstrated that ATM kinase and its downstream target CHK2 were constitutively phosphorylated in ALT cell lines but not in telomerase-positive cell lines (Figure. 2D & 2E; *P*<0.01). Consistent with immunoblotting, ALT neuroblastoma cell lines have numerous p-ATM (s1981) foci (by immunofluorescence) in the nucleus relative to two telomerase-positive cell lines (Supplementary Figure. S3). Consistent with the ATM activation observed *in vitro*, ALT patient-derived xenografts (PDXs), but not a telomerase-positive PDX, were also positive for p-ATM (s1981) by immunohistochemistry (IHC) (Figure. 2F). We did not observe selective ATR/CHK1 activation in ALT relative to telomerase-positive neuroblastoma cell lines (Figure 2D and Supplementary Figure. S4; *P* = ns). As we have observed that ALT neuroblastoma cell lines have a basal level of DNA damage, we determined if ALT cells are defective in repairing DNA by assessing for γ-H2AX at different time points following irradiation. We observed that both ALT cell lines and a telomerase-positive cell line robustly induced γ-H2AX at 1 hour following irradiation that returned to basal levels of γ-H2AX by 24 hours following irradiation (Supplementary Figure. S2D), suggesting that ALT and telomerase-positive cell lines have a similar ability to repair DNA damage following genotoxic stress.

### ALT neuroblastoma cell lines manifested constitutive telomere damage signaling via ATM kinase

As we did not observe a difference between ALT and telomerase-positive cell lines in DNA repair following exogenous genotoxic stress, we determined if constitutive DNA damage signaling occurred in ALT cells at telomeres. We assessed telomere dysfunction induced foci (TIFs) (*42, 43*), detected by the co-localization of the DNA damage repair marker 53BP1 (immunofluorescence) to telomeres (identified by fluorescence *in situ* hybridization (FISH)). We observed that ∼40% of 53BP1 foci localized to telomeres in ALT neuroblastoma cell lines (Figure 3A and 3B). However, it is important to note that ALT cell lines often have heterogeneous telomere length with some chromosomal ends having undetectable telomere signals, making it difficult to determine if all of the DNA damaging signals emanate from telomeres or from other genomic loci. Further, we observed that ALT-positive cell lines have a substantially higher number of TIFs compared to 3 *TP53* mutant telomerase-positive cell lines (Figure. 3C), which is consistent with previous findings in fibroblast transformed ALT cell lines (*44*).

**Figure 3.**
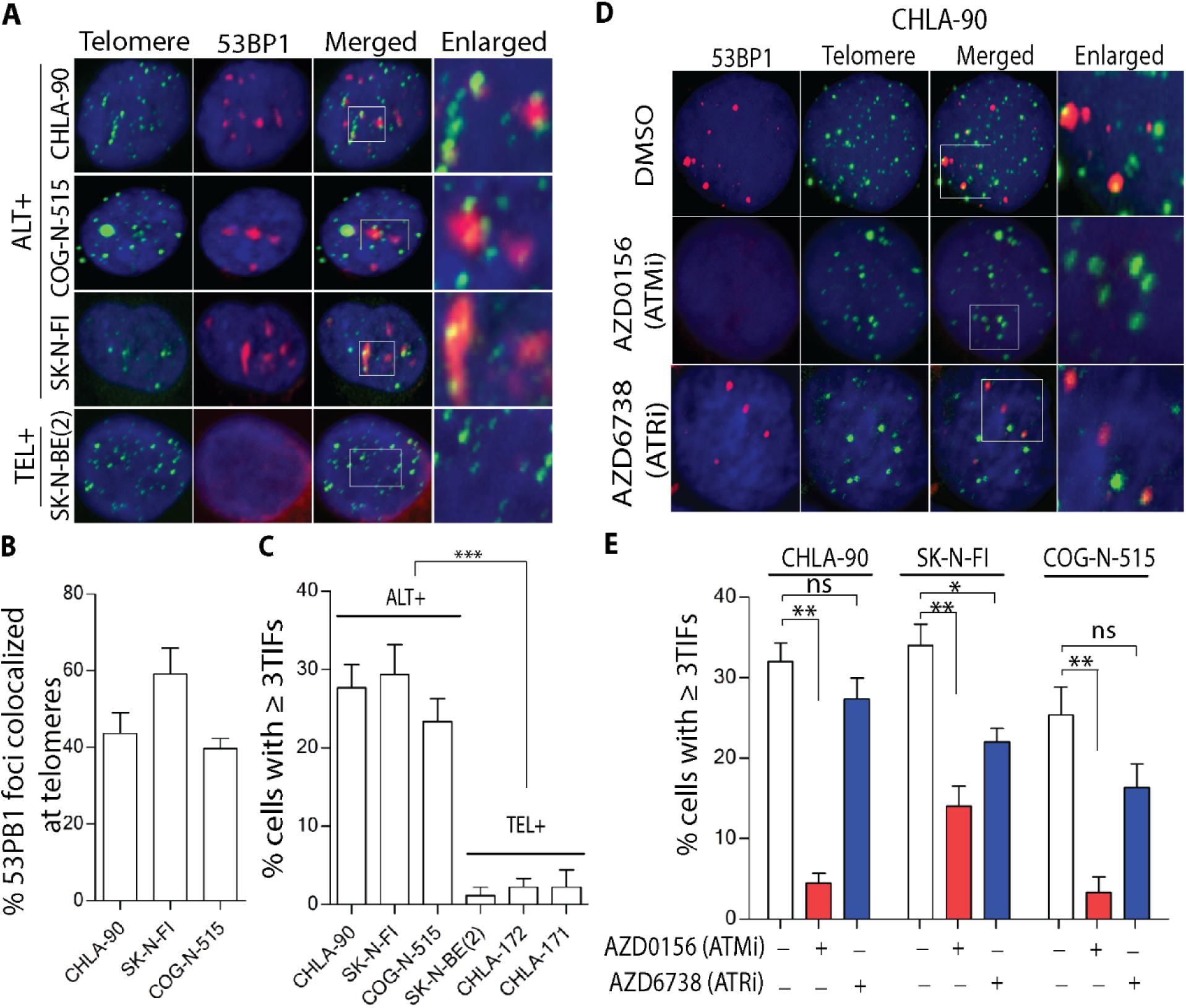
ALT neuroblastoma cell lines manifested constitutive ATM dependent telomere damage signaling. (A) Representative images of immunofluorescence in combination with florescent *in situ* hybridization (IF-FISH) to detect telomere dysfunction-induced foci (TIFs) in ALT (n=3) versus telomerase-positive cell line. 53BP1 was detected by IF (red) and telomeres by FISH with a [TTAGGG] 3 probe (green). (B) Percentage of 53BP1 foci that are co-localized to telomeres in 3 ALT cell lines. (C) Percentage of cells with ≥ 3 TIFs in ALT versus telomerase- positive (TEL+) cell lines. (D) Representative IF-FISH staining to detect TIFs in the ALT neuroblastoma cell line CHLA-90 treated with the ATM inhibitor AZD0156 (100 nM) or the ATR inhibitor AZD6738 (300 nM). Cells treated with DMSO were used as controls. 53BP1 was detected by IF (red) and telomeres by FISH with a [TTAGGG]3 probe (green). (E) Bar graph shows percentage of cells with ≥ 3 TIFs in ALT cell lines treated with ATM or ATR inhibitor relative to vehicle. A minimum of 50 cells for each experimental replicate were assessed using IF- FISH for B, C and E. The bars represent mean with SD from three experimental replicates. Statistical significance was calculated using Wilcoxon-rank sum test for C and two-tailed t-test for E. ***: *P*<0.001, **: *P*<0.01, *: *P*<0.05, ns: not significant.

As 53BP1 foci could be induced due to activation of either ATM or ATR kinase signaling (*43*), we assessed the effect of ATM or ATR inhibition on TIFs in ALT neuroblastoma cell lines (Figure. 3D). We observed that the frequency of TIFs were consistently diminished with the inhibition of ATM kinase (Figure. 3D and 3E; *P*<0.01) in ALT neuroblastoma cell lines, pointing towards an ATM-dependent DNA damage response at telomeres. However, ATR inhibition had modest to no effect on TIFs in ALT neuroblastoma cell lines; differences were not statistically significant in 2 of 3 ALT cell lines (Figure. 3E).

### Induction of ATM-dependent TIFs in a p53 non-functional telomerase-positive cell line was associated with high resistance to TMZ+SN-38

It is well known that in a *p53*-deficient setting, ATM protects cells from DNA-damaging chemotherapy by inducing DNA repair (*45*). However, it is not known if constitutive activation of ATM has any effect on response to DNA-damaging chemotherapy. To mimic the ATM-dependent TIF response observed in ALT cells in a telomerase-positive p53 non-functional cell line, we over expressed a dominant-negative TRF2 (TRF2ΔBΔM; Figure. 4A) using a retroviral vector in a telomerase-positive *TP53* mutant (SK-N-BE (2)) neuroblastoma cell line. The dominant-negative TRF2 is known to induce an ATM kinase- dependent TIF response at telomeres (*44*). Expression of TRF2ΔBΔM in SK-N-BE (2) (Figure. 4B) induced ATM activation (phospho-ATM) in both the TRF2ΔBΔM clones relative to the empty vector control (Figure. 4B and 4C). As expected, we were able to induce ATM-dependent TIFs in cells transduced with pLPC-nMyc-TRF2ΔBΔM and the TIFs were abrogated by treating with the ATM inhibitor AZD0156 (Figure 4D).

**Figure 4.**
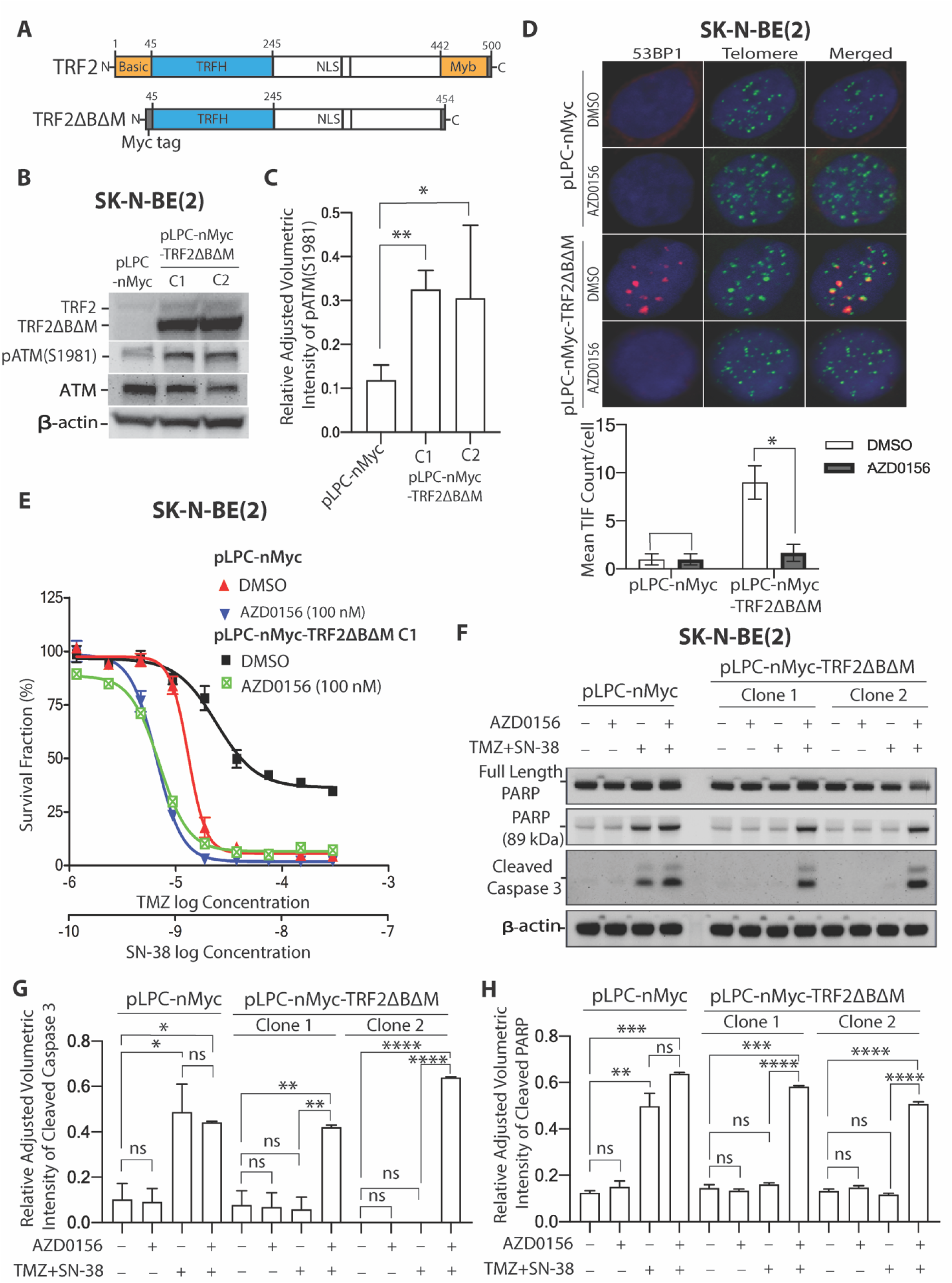
Induction of ATM dependent TIFs caused hyper-resistance to TMZ+SN-38 that can be reversed with the ATM inhibitor AZD0156. (A) Schematic of TRF2 and dominant- negative form of TRF2 (TRF2 ΔBΔM). (B) Immunoblotting for TRF2, pATM (S1981), ATM and β-actin in cells transduced with pLPC-nMyc or pLPC-nMyc-TRF2ΔBΔM. (C) Quantitative analysis of immunoblotting for pATM (S1981) in B and its replicates. (D) (Top panel) Representative images of telomere dysfunctional foci (TIF) analysis using IF-FISH in same cells as in B (Clone 1 only) +/-ATM inhibitor (AZD0156). 53BP1 was detected by IF (red) and telomeres by FISH with a [TTAGGG]3 probe (green). (Bottom panel) Bar graph showing meant TIF count in same cells as in B +/-ATM inhibitor (AZD0156). A minimum of 50 cells for each experimental replicate were assessed using IF-FISH. (E) DIMSCAN cytotoxicity assay curves in response to TMZ+SN-38 +/- AZD0156 in same cells as shown in C. (F) Immunoblotting for PARP, cleaved-PARP, cleaved caspase-3, and β-actin in same cells as shown in B treated with TMZ+SN-38 +/- AZD0156, cells with no treatment were included for comparison. (G) Quantification of cleaved caspase 3 and (H) cleaved PARP for immunoblot in F and its replicates. The bars represent means with SDs from three experimental replicates. Statistical significance was calculated using two-tailed t-test for G and H.****: P<0.0001; ***: *P*<0.001, **: *P*<0.01, *: *P*<0.05, ns: not significant.

To evaluate the effect of constitutive ATM activation on DNA-damaging chemotherapy in neuroblastoma cell lines, we assessed the cytotoxic response to TMZ+SN-38 (a drug combination used to salvage relapsed neuroblastoma patients) in SK-N-BE(2) transduced with pLPC-nMyc- TRF2ΔBΔM or pLPC-nMyc (empty vector) +/- the ATM inhibitor AZD0156. Cells transduced with pLPC-nMyc-TRF2ΔBΔM showed increased resistance to TMZ+SN-38 when compared to the empty vector control (Figure.4E & Supplementary Figure. S5A; IC50: *P*<0.05; two-way ANOVA: *P*<0.001). Transduction of pLPC-nMyc-TRF2ΔBΔM into another p53 non-functional telomerase-positive cell line (CHLA-171) also increased resistance to TMZ+ SN-38 (Supplementary Figure. 5B; IC50: *P*<0.05; two-way ANOVA: *P*<0.001). Consistent with the cytotoxicity assay, cells transduced with pLPC-nMyc-TRF2ΔBΔM showed lower induction by TMZ+ SN-38 of markers for apoptosis, caspase-3, and PARP cleavage. (Figure. 4F-H). Addition of the ATM inhibitor AZD0156 to TMZ+SN-38 reversed the chemo-resistance in cells transduced with pLPC-nMyc-TRF2ΔBΔM (Figure 4E-H, Supplementary Figure. S5A and S5B; IC50: *P* = ns; two-way ANOVA: *P* = ns).

**Figure 5.**
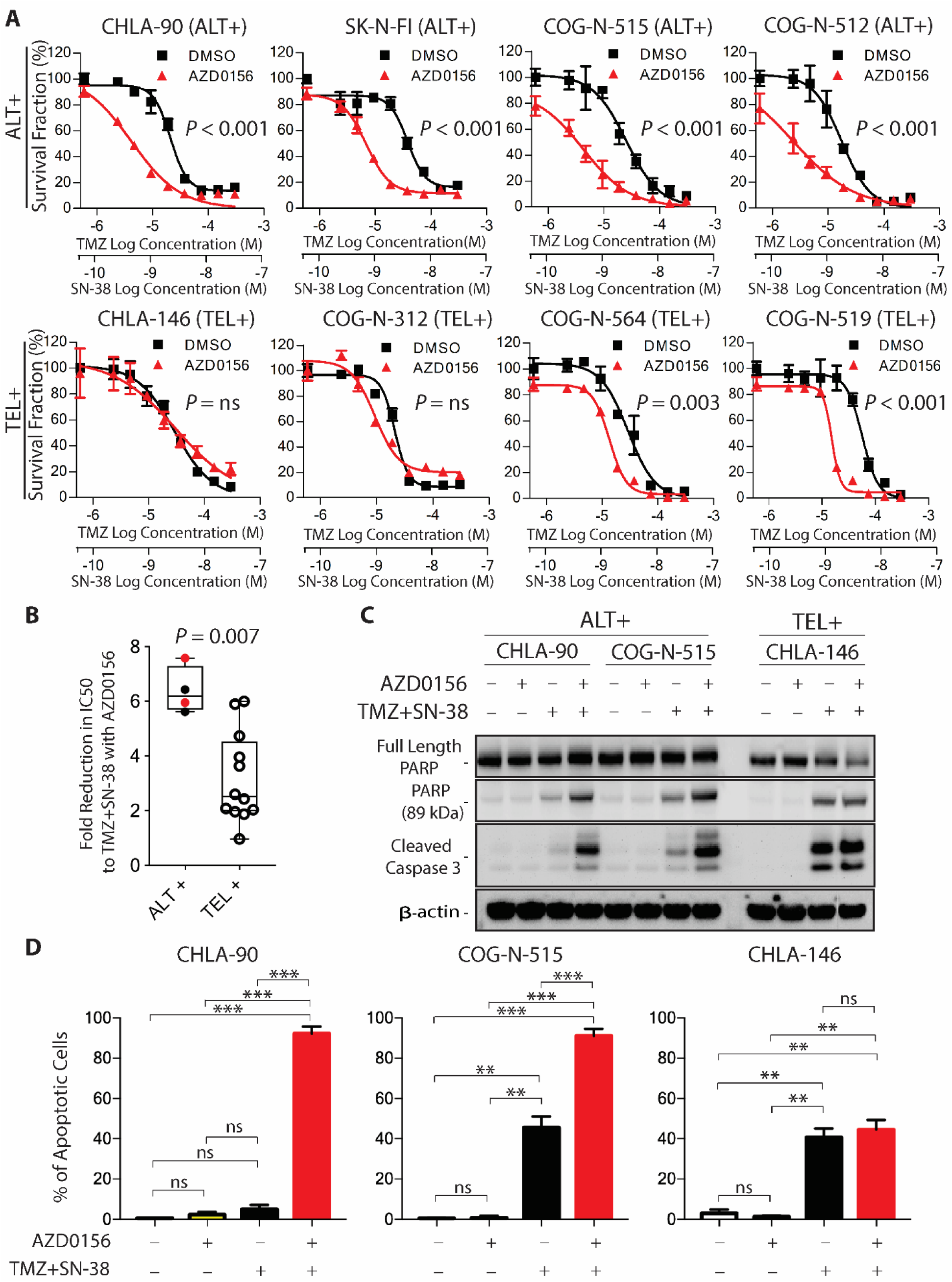
The ATM inhibitor AZD0156 sensitized ALT neuroblastoma cell lines to TMZ+SN-38. (A) DIMSCAN cytotoxicity assay curves in ALT (n=4) and chemo-resistant telomerase- positive (n=4) cell lines treated with TMZ+SN-38 +/- AZD0156 (100 nM). Statistical significance was calculated using two-way ANOVA. (B) Fold-change in IC50 with the addition of AZD0156 to TMZ+SN-38 in ALT (n=4) versus telomerase-positive (n=12) cell lines *in vitro*. ALT cell lines COG-N-512 and COG-N-515 that were established from same patient are highlighted in red. Statistical significance was calculated using Wilcoxon-rank sum test. (C) Immunoblotting for PARP, cleaved-PARP, cleaved caspase-3, and β-actin in ALT cell lines (CHLA-90 and COG-N-515) and a telomerase-positive cell line (CHLA-146) treated with TMZ+SN-38 +/- AZD0156 (100 nM). (D) Tunnel assay to detect apoptosis using flow cytometry in same cells as shown in C. The bars represent means with SDs from three experimental replicates. Statistical significance was calculated using two-tailed t-test. ***: *P*<0.001, **: *P*<0.01, *: *P*<0.05, ns: not significant.

As overexpression of TRF2ΔBΔM can increase the frequency of non-homologous end joining (*46*), we employed an alternative approach to induce ATM dependent TIFs in the SK-N- BE (2) cell line by overexpressing the nuclease domain of FOKI fused to N-terminus domain of shelterin protein TRF1 (FokI^WT^TRF1) using a retroviral vector. A nuclease dead mutant (FokI^D450A^TRF1) fused to TRF1 was used as a control (Supplementary Figure. S6A). As expected, fusion proteins expressed in these cells co-localized to telomeres in both FokI^WT^TRF1 and FokI^D450A^TRF1 (Supplementary Figure. S6B). Cells expressing FokI^WT^TRF1 showed higher phosphorylation of ATM compared to nuclease dead mutant and the empty vector control (Supplementary Figure. S6C and S6D). Consistent with previous work, (*47*) we were able to induce ATM-dependent TIFs in cells transduced with pLPC-FokI^WT^TRF1 and the TIFs were abrogated when treated with the ATM inhibitor AZD0156 (Supplementary Figure. S6E).

We assessed the cytotoxic response to TMZ+SN-38 in SK-N-BE (2) transduced with pLPC- FokI^WT^TRF1 or pLPC- FokI^D450A^TRF1 (control vector) +/- the ATM inhibitor AZD0156. Cells transduced with pLPC-FokI^WT^TRF1 were relatively more resistant to TMZ+SN-38 when compared to cells transduced with the nuclease dead (pLPC- FokI^D450A^TRF1) transduced cells (Supplementary Figure. S6F; IC50: *P*<0.05; two-way ANOVA: *P*<0.001). Addition of the ATM inhibitor AZD0156 to TMZ+SN-38 reversed the chemo-resistance in cells transduced with pLPC- FokI^WT^TRF1 reaching cytotoxicity levels equal to control vector transduced cells treated with AZD0156+TMZ+SN-38 (Supplementary Figure. S6F; IC50: *P* = ns; two-way ANOVA: *P* = ns).

### The ATM inhibitor AZD0156 reversed resistance of ALT neuroblastoma cell lines to TMZ+SN-38 *in vitro*

To evaluate the therapeutic potential for inhibition of ATM activation in ALT cells, we knocked-down ATM using lentiviral shRNA in two ALT neuroblastoma cell lines (CHLA-90 and COG-N-515; Supplementary Figure S7A). Knockdown or pharmacological inhibition of ATM (Supplementary Figure. S7A, S7C-D) reduced DNA C-circle content (Supplementary Figure. S7B and S7E) in ALT neuroblastoma cell lines, but cell viability was not altered following ATM knockdown or inhibition, indicating that inhibition of ATM activation by itself does not have an immediate effect on ALT cell survival.

As a small molecule ATM inhibitor (AZD0156) showed reversal of drug resistance in cells with ATM dependent TIFs (Figure 4E, 4F and S6F), we determined if pharmacological inhibition of ATM was effective in sensitizing ALT cell lines to TMZ+SN-38, by comparing the cytotoxic response to TMZ+SN-38 +/-AZD0156 in ALT (n=4) versus telomerase-positive (n=12) cell lines *in vitro;* the telomerase-positive *in vitro* models selected here were those least responsive to TMZ+SN38 in Figure 1B. AZD0156 consistently cause a marked increase in sensitivity of ALT neuroblastoma cell lines to TMZ+SN-38 (Figure 5A, fold-change in IC 50: *P* < 0.005; two-way ANOVA: *P*<0.001), while telomerase-positive cell lines showed a heterogeneous response to the combination (7/12 telomerase-positive cell lines showed enhanced cytotoxicity to TMZ+SN38 with AZD0156: fold-change in IC 50: *P* < 0.05; two-way ANOVA: *P*<0.01; Figure. 5A and Supplementary Figure. S8). The ATM kinase inhibitor KU60019 also sensitized ALT neuroblastoma cell lines to TMZ+SN-38 (Supplementary Figure S9; fold-change in IC 50: *P* < 0.001; two-way ANOVA: *P*<0.001).

Across all cell lines, the fold decrease in IC50 for TMZ+SN-38 with the addition of AZD0156 was significantly greater in ALT relative to telomerase-positive cell lines (Figure 5B; *P* = 0.007). AZD0156 at concentrations as low as 6.25 nM significantly sensitized ALT cells to TMZ+SN-38 (Supplementary Figure. S10; P<0.05). We assessed the combination index (CI) in ALT neuroblastoma cell lines using fixed-ratio dosing and demonstrated that AZD0156 synergized with TMZ+SN-38 in all 4 ALT cell lines *in vitro* (Supplementary Figure. S11; Supplementary Table. S4; CI < 1). Additionally, we observed a dramatic increase in PARP cleavage, caspase-3 cleavage, and TUNEL staining in ALT cells treated with TMZ+SN- 38+AZD0156, relative to TMZ+SN-38 alone (Figure. 5C, 5D and supplementary Figure. S12), indicating that synergy is driven by apoptotic cell death. A telomerase-positive cell line (CHLA-146) with no enhancement in cytotoxicity to TMZ+SN38 with AZD0156 was used for comparison. Consistent with the results obtained using pharmacological ATM inhibition, ATM knockdown sensitized ALT cell lines to TMZ+SN-38 (Supplementary Figure. S13; fold-change in IC 50 *P*<0.001; two-way ANOVA: *P*<0.001).

ALT cells have been reported to be to be hypersensitive to ATR inhibitors (*13*), but we did not see a marked difference in sensitivity to an ATR inhibitor (AZD6738) between ALT and telomerase-positive cell lines (Supplementary Figure. S14A; *P* = ns). Additionally, we did not see consistently enhanced sensitization to TMZ+SN-38 with the ATR inhibitor AZD6738 in ALT neuroblastoma cell lines *in vitro* (Supplementary Figure. S15B).

Thus, the ATM inhibitor AZD0156 significantly enhanced TMZ+SN-38 activity in all ALT neuroblastoma cell lines, but to lesser degree in a subset of comparably chemo-resistant telomerase-positive neuroblastoma cell lines *in vitro*. Reversal of drug resistance in ALT cell lines was not observed with an ATR inhibitor.

### The ATM inhibitor AZD0156 enhanced the activity of TMZ+IRN in ALT patient-derived (PDX) and cell line-derived (CDX) xenografts

We assessed the activity of the ATM inhibitor AZD0156 in combination with TMZ+IRN in 4 ALT neuroblastoma xenograft models (3 CDXs and 1 PDX) and 2 telomerase-positive neuroblastoma PDXs. We determined that AZD0156 at 20 mg/kg was well tolerated when sequenced following treatment with TMZ+IRN in nu/nu mice.

Pharmacokinetic assessment of AZD0156 at 20 mg/kg showed that maximum unbound plasma concentration was 204 nM at 2 hours after dosing and dissipating over a 24-hour period (Supplementary Figure. S15; Supplementary Table S5). AZD0156 alone had low activity against 3 of 4 ALT xenografts and intermediate activity against 1 of 4 ALT xenografts, compared to only low activity in 1 of 2 telomerase-positive xenografts (Figure. 6A and 6B; Table 1). AZD0156 alone failed to induce objective responses in any of the xenografts (Table 1). All mice in control and AZD0156 treated groups showed progressive disease (PD) (Supplementary Figure. S16; Table 1). Treatment of ALT xenografts with TMZ+IRN increased event-free survival (EFS) relative to control and AZD0156 groups (Figure. 6A; Table 1). TMZ+IRN achieved partial responses in most of the mice in 2 of 4 ALT xenograft models (CHLA-90m and SK-N-FIm), progressive disease in an ALT PDX (COG-N-625x), and complete responses in ALT COG-N-515m (Supplementary Figure. S16; Table 1). Although objective responses were observed in 3 of 4 ALT xenografts, responses were maintained for < 60 days in most indicating severe (COG-N-625x) or partial (CHLA-90m and SK-N-FIm) chemo-resistance to TMZ+IRN *in vivo* (Supplementary Figure. S16A; Table 1).

**Figure 6.**
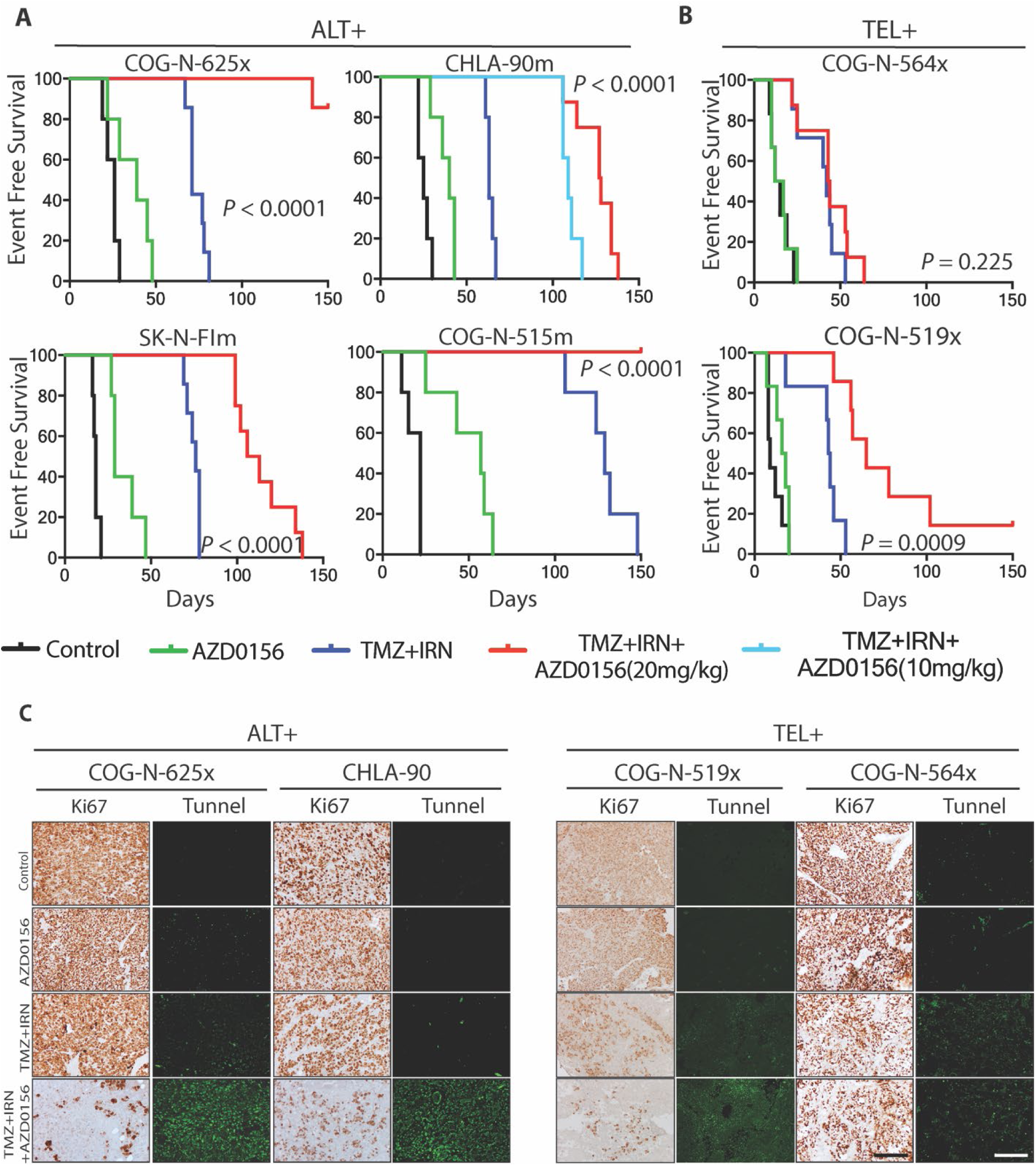
ATM inhibitor AZD0156 enhanced temozolomide + irinotecan activity in ALT PDX and CDXs *in vivo*. (A) Kaplan–Meier event-free survival (EFS) curves according to each treatment group for tumor-bearing either ALT PDX (COG-N-625x) or ALT CDXs (CHLA-90m, SK-N-FIm or COG-N-515m). EFS was defined as the time taken for tumors to reach 1500 mm^3^ in tumor volume, from initiation of treatment, or to death from any cause. Mice treated with temozolomide (TMZ) + irinotecan (IRN) were given 2 cycles of TMZ (25 mg/kg) + IRN (7.5 mg/kg) on Days 1-5 in a 21-day cycle; For the TMZ+IRN+AZD0156 group, AZD0156 (20 mg/kg) was administered following TMZ+IRN treatment on Days 6-19. An additional TMZ+IRN+AZD0156 group with a lower dose of AZD0156 (10 mg/kg) for the ALT xenograft CHLA-90 is included. COG-N-515m mice treated with 2 cycles of TMZ (25 mg/kg + IRN (7.5 mg/kg) on Days 1-5 in a 21 day cycle were repurposed in Figure 1C and 1D. (B) Kaplan–Meier EFS curves according to each treatment group for tumor-bearing telomerase-positive PDXs (COG- N-564x and COG-N-519x). (C) Representative images of immunohistochemistry for Ki67 and immunofluorescence staining for apoptosis using tunnel assay (in green) on FFPE sections in each treatment group. KI-67 and tunnel staining are shown for 1 ALT PDX (COG-N-625x), 1 ALT CDX (CHLA-90m) and 2 telomerase-positive PDX (COG-N-519x and COG-N-564x) models. FFPE sections were obtained from resected tumor on day 9 from initiation of treatment. Scale bar represeents 50 µm. EFS was assessed for statistical significance using the log-rank test.

**Table 1.**
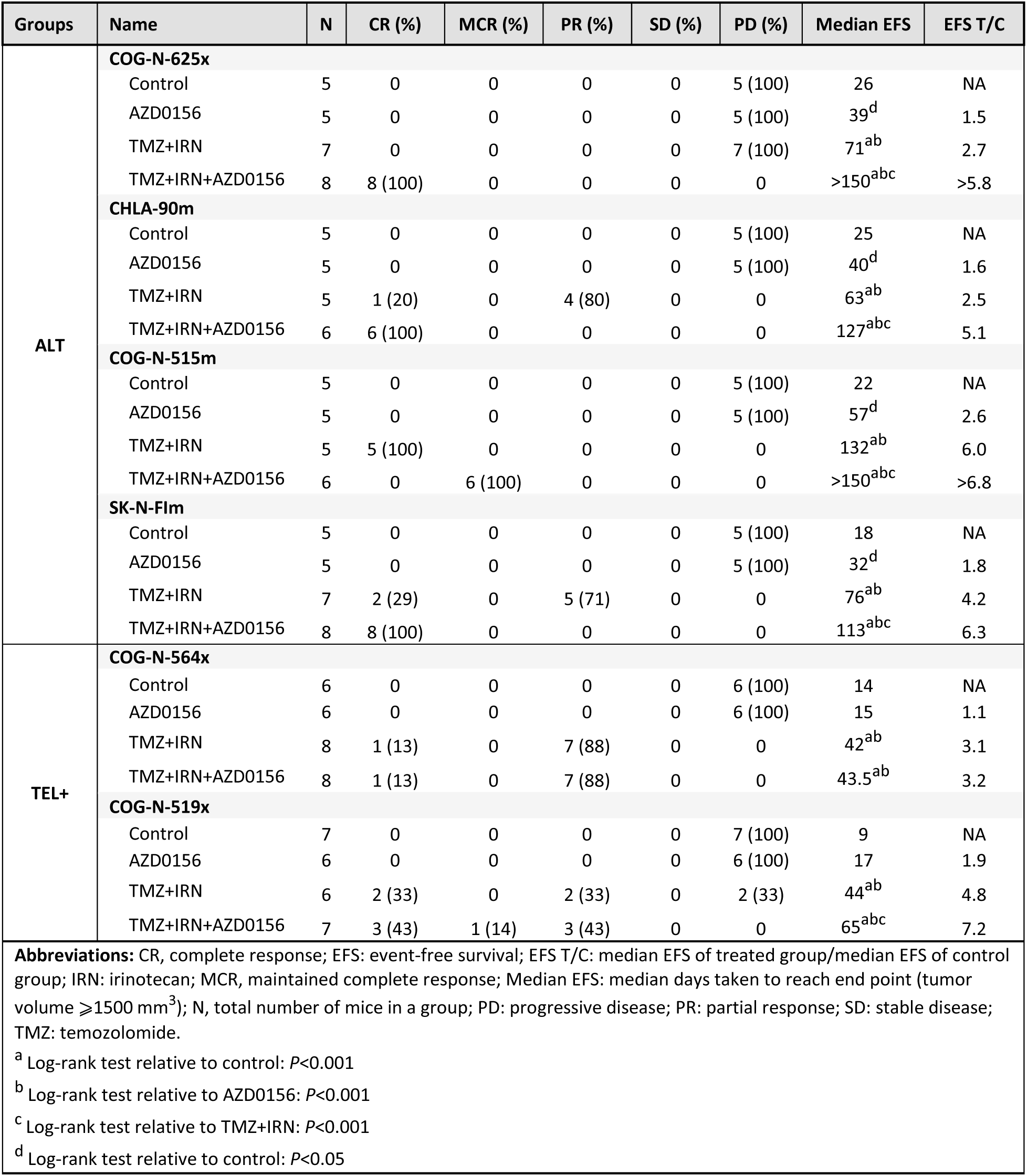
Summary of *in vivo* response to TMZ+IRN+AZD0156, TMZ+IRN, AZD0156, and no treatment control in 4 ALT and 2 telomerase-positive xenograft models.

To assess the effect of ATM inhibition on TMZ+IRN activity, we treated mice first with TMZ+IRN (days 1 to 5) and then followed with AZD0156 given on days 6 to 19, to mimic clinical use and prior *in vivo* preclinical studies of AZD0156 in combination with IRN (*48*). All mice bearing ALT xenografts treated with TMZ+IRN+AZD0156 achieved complete responses (Supplementary Figure. S16A; Table 1) and EFS was significantly longer for mice treated with TMZ+IRN+AZD0156 relative to control, AZD0156 alone, or TMZ+IRN groups (Figure 6A; Table 1; log-rank test: *P*<0.0001). In the CHLA-90m xenograft model, mice treated with TMZ+IRN+AZD0156 (AZD0156 at 20 mg/kg) showed greater activity than mice treated with TMZ+IRN+AZD0156 (AZD0156 at 10 mg/kg) (Figure 6A. log-rank test: *P*=0.005). The combination of AZD0156+TMZ+IRN was well tolerated as measured by mouse weights over time (Supplementary Figure. S17). Histopathological examination at day 9 of therapy for xenografts treated with TMZ+IRN+AZD0156 showed a reduction in proliferative activity by Ki-67 immunohistochemistry and increased apoptosis by TUNEL staining relative to mice in the control, AZD0156, and TMZ+IRN treatment groups (Figure. 6C).

To assess the activity of AZD0156+TMZ+IRN in telomerase-positive neuroblastoma, we chose two telomerase-positive PDXs (COG-N-564x and COG-N-519x) that show resistance to TMZ+IRN, comparable to that seen in the ALT PDX (COG-N-625x). Mice treated with TMZ+IRN had increased EFS compared to mice in the control and AZD0156 groups in both the telomerase-positive PDXs (Figure 6B and Supplementary Figure. S16B; Table 1). Mice treated with TMZ+IRN+AZD0156 showed an increase of EFS in COG-N-519x but no improvement of EFS in COG-N-564x relative to TMZ+IRN (Figure 6B and Supplementary Figure. S16B; Table 1). Histopathological examination of COG-N-519x and COG-N-564x mice treated with TMZ+IRN+AZD0156 had substantial reduction in proliferative activity and showed increased apoptosis when compared to the TMZ+IRN group in COG-N-519x but not in COG-N-564x (Figure. 6C). The median EFS (Table 1) was > 100 days for all four ALT xenografts treated with TMZ+IRN+AZD0156 (> 150 days for 2 of 4), while median EFS for telomerase-positive PDXs was 43.5 and 65 days.

## Discussion

About 25-30% of high-risk neuroblastomas (age > 18 months) activate the ALT mechanism to maintain telomeres (*28, 29*), with a long-term overall survival under 25%, despite intensive multimodal treatment regimens (*29*). Unfortunately, relapse or progression in ALT neuroblastoma patients following front-line therapy is almost uniformly fatal (*29*). Clinical observations have categorized presumptive ALT neuroblastoma tumors as chemo-resistant (*32*). Although potentially therapeutically actionable genomic alterations such as *ATRX* are being investigated in ALT neuroblastoma (*49*), ∼50% of ALT neuroblastoma tumors lack any known genomic alterations (*27, 29*). Furthermore, due to a prior lack of robust *in vitro* and *in vivo* ALT neuroblastoma models, development of targeted therapy against ALT neuroblastoma has been challenging. We recently established and characterized a small panel of ALT neuroblastoma patient-derived cell lines (PCLs) and a patient-derived xenograft (PDX) by screening a large number of patient-derived neuroblastoma cell lines and PDXs (*29*). As expected from their indolent clinical behavior, ALT neuroblastomas are more difficult to capture as PCLs and PDXs. ALT neuroblastomas take longer to establish and generally grow slower than PCLs or PDXs from telomerase-positive neuroblastomas (with and without *MYCN* amplification), hence contributing to the more limited numbers of available ALT PCLs and PDXs.

Using PCLs and PDXs from neuroblastoma patients with progressive disease that were established and/or characterized in our lab (*29, 39*), we demonstrated both *in vitro* and *in vivo* that, relative to telomerase-positive neuroblastoma, ALT neuroblastoma was associated with hyper-resistance to DNA damaging agents used to treat neuroblastoma patients. ALT neuroblastoma PCLs and PDXs are p53 non-functional, in line with previous reports on ALT cancers (*39, 50, 51*). Although, p53 non-functionality induces chemotherapy resistance in neuroblastoma (*41*), ALT neuroblastoma PCLs showed greater *de novo* resistance to DNA damaging agents than did p53 non-functional telomerase-positive lines. Thus, based on our data, the ALT phenotype is associated with hyper-resistance to DNA damaging agents in neuroblastoma.

To elucidate chemo-resistant mechanisms operative in ALT neuroblastoma, we performed gene set enrichment analysis in comparing ALT to telomerase-positive non-*MYC*N-amplified PCLs and we observed an enrichment of ATM and ATR-BRCA pathways in ALT cell lines. ATM, but not ATR, was specifically activated in ALT but not in telomerase-positive PCLs *in vitro* and in PDXs *in vivo*. Additionally, ALT cells manifested evidence of constitutive DNA damage signaling. However, ALT PCLs repaired DNA upon irradiation, indicating that endogenous DNA damage is not due to major defects in DNA repair. Previous studies on the ALT phenotype suggested that ALT cells have high levels of spontaneous telomere DNA damage (*44*). We demonstrated in ALT neuroblastoma cell lines large amounts of DNA damage foci that were localized to telomeres, indicating that telomere dysfunction is partly responsible for constitutive DNA damage signaling in ALT neuroblastoma. We observed that the frequency of telomere dysfunctional foci (TIFs) markedly diminished with the inhibition of ATM kinase in ALT cell lines, pointing towards an ATM-dependent DNA damage response at telomeres. Our data indicate that constitutive ATM activation is at least partially due to telomere dysfunction in ALT neuroblastoma.

ATM kinase is known to act as a binary switch to control the contribution of p53 signaling to the DNA damage response (*45*). In the presence of functional p53, ATM kinase activation largely contributes to induction of apoptosis via p53. In cells with loss of p53 function, apoptosis is reduced and ATM signaling is redirected to promote homologous recombination mediated DSB repair. The net result of this switch in ATM signaling leads to increased cellular survival in response to genotoxic stress (*45*), but how endogenous ATM activation (such as we observed in ALT cells) affects the response to genotoxic stress in a p53 deficient setting is unclear. To investigate the role of constitutive ATM activation in chemo-resistance, we artificially induced telomere dysfunction by forced expression of a dominant-negative TRF2 or TRF1-FOKI in a p53 non-functional telomerase-positive cell line, as described previously (*44*). Forced generation of telomere dysfunction in p53-non-functional telomerase-positive cells induced hyper resistance to temozolomide (TMZ) + irinotecan (IRN, as the active metabolite SN-38 i*n vitro*), DNA damaging agents used commonly to salvage relapsed neuroblastoma patients. ATM inhibition reversed resistance to TMZ + SN-38, indicating constitutive ATM activation (in response to telomere dysfunction) in ALT cells is at least partly responsible for the chemo-resistant phenotype.

We demonstrated that AZD0156, a clinical-stage ATM kinase inhibitor, sensitized ALT neuroblastoma cell lines and xenografts to TMZ + IRN (as SN-38 i*n vitro*). By contrast, modest or no enhancement of TMZ + IRN activity was observed in telomerase-positive models *in vitro* and *in vivo.* Although AZD0156 consistently (and to a greater degree) enhanced the activity of TMZ + IRN in ALT cell lines and xenografts, our data suggest that a subset of telomerase-positive neuroblastomas showed an enhanced response to TMZ+IRN when combined with AZD0156, indicating that addition of ATM inhibitor to TMZ+IRN might be an useful approach in treating a subset of telomerase positive neuroblastomas. However, it is currently unclear how to identify telomerase-positive tumors for which AZD0156 may enhance the activity of TMZ + IRN.

In summary, ALT neuroblastomas are a distinct subgroup of patients with a robust biomarker (DNA C-circles) that are in great need of novel therapeutic options. We have demonstrated that telomere dysfunction induced ATM kinase in ALT neuroblastomas, and that constitutive activation of ATM kinase promoted resistance to DNA damaging therapy observed in ALT neuroblastoma. We also showed that the clinical-stage ATM inhibitor AZD0156 can reverse resistance to temozolomide + irinotecan in ALT neuroblastoma cell lines and xenografts. These data support undertaking early-phase clinical trials of AZD0156 with temozolomide + irinotecan in children with neuroblastoma.

## Materials and Methods

### Study Design

The objectives of this study were to identify mechanism(s) of chemo-resistance in relapse/refractory ALT neuroblastoma and to identify a means of reversing such resistance. To enable *in vitro* and *in vivo* experimentation, we used ALT and telomerase-positive neuroblastoma patient-derived cell lines (PCLs), patient-derived xenografts (PDXs), and cell line-derived xenografts (CDXs) established from neuroblastoma patients at progressive disease or postmortem from progressive disease. Using genetic and pharmacological experimentation, we found that telomere dysfunction drives chronic ATM activation, which in turn induced chemo-resistance to DNA damaging agents in ALT neuroblastoma. To determine if we can reverse the chemo-resistant phenotype in ALT PDXs and CDXs *in vivo*, we employed the clinical-stage orally available ATM kinase inhibitor AZD0156 given following treatment with TMZ + IRN. We used 4 ALT xenografts (3 CDX and 1 PDX) and 2 telomerase-positive PDXs to assess the activity of AZD0156 alone or in combination with TMZ+IRN. Sample size was determined by previous experimental experience and based on the growth profile of individual models (n= 5 to 8), with 8 animals providing an 80% power (at 0.05 significance level) for a Wilcoxon rank-sum test to detect a significant difference between the treated and control mice, assuming that the standard deviation is 13.4 days, the increase in median survival is 40 days, and no censoring in the xenograft experiments. This was not a blinded study at the time of investigation. The endpoint was tumor volume ≥ 1500 mm^3^, if mouse weight was reduced by ≥ 20% of initial weight, if the mice exhibited signs of impaired health, or death from any cause. Mice were monitored until they reached endpoint of the study or until 150 days from the start of treatment.

### RNA Sequencing

RNA sequencing libraries were prepared from polyA-selected RNA and sequenced on an Illumina HiSeq 2500 as described in (31). The FASTQ formatted sequence reads were aligned to GRCh37.69 using the STAR RNA Seq aligner (v2.3.1) (32) and transcripts were assembled and quantified using Cufflinks v2.1.1 (33). GSEA was performed using the GSEA v4.0.2 (http://software.broadinstitute.org/gsea) using normalized read counts.

### DIMSCAN Cytotoxicity Assay

DIMSCAN was used to assess cytotoxic drug response as previously described (*52*). Cytotoxic response to commonly utilized DNA damaging agents was determined by assessing survival fraction following treatment with a range of drug concentrations that included clinically achievable levels of drug concentration for 96 hours. For combination studies with temozolomide (TMZ) + SN-38 (the active metabolite of irinotecan), cells were treated with fixed ratio serial dilutions of drugs (maximum dose of 300 μM for TMZ and 30 nM for SN-38); cytotoxic response was assessed after 7 days from initiation of drug treatment. To test activity of the ATM inhibitor AZD0156 in combination with TMZ+SN-38, cells were treated with a constant dose of AZD0156 (100 nM) 1h prior to TMZ+SN-38. IC50 values (concentration that is cytostatic or cytotoxic to 50% of the control value) were calculated as described previously (*52*).

### Plasmids and Cloning

pLPC TRF2 deltaB deltaM (#19008), pLPC-MYC (#12540), and pLPC-MYC-hTRF1 (#64164) were obtained from Addgene. Cambridge, MA. FOK1^WT^and FOKI^D450A^ nuclease domain were custom synthesized using Integrated DNA Technologies (IDT) custom synthesis tool, tailored to insert at BamHI restriction site in pLPC-MYC-hTRF1 plasmid.

### Immunofluorescence and Telomere Fluorescence In Situ Hybridization (IF-FISH)

IF and IF- FISH were performed as described previously (*29*). Further details are available in supplementary methods.

### Immunoblotting

Immunoblotting was performed as described previously (*53*). Further details are available in supplementary methods.

### Tunnel assay

Tunnel assay was performed using BD APO-DIRECT kit on BD LSR2, as described previously (*54*).

### *In vivo* Drug Testing

The TTUHSC Institutional Animal Care and Use Committee (IACUC) approved all animal protocols. Xenograft drug testing was as previously described (*53*). Additional details are available in supplementary methods.

### Statistical Analysis

Comparison of two sample sets was done using Mann-Whitney U test. Comparison of biological replicates was based on unpaired two-tailed t-test with Welch’s correction. Dose response curves were assessed using two-way-ANOVA test. IC50 concentration was calculated, as described previously (*52*); comparison of IC50 was based on unpaired two-tailed t-test with Welch’s correction. Survival analysis for xenograft studies was done by the Kaplan-Meier method, as assessed using a log-rank test. All statistical analysis was done using GraphPad prism v7.0 and were considered statistically significant if *P* ≤ 0.05.

## Supplementary Methods

### List of Supplementary Materials

**Figure S1.** ALT neuroblastoma models were associated with resistance to DNA damaging agents irrespective of p53 functionality.

**Figure S2:** ALT neuroblastoma was associated with constitutive DNA damage.

**Figure S3:** ALT neuroblastoma cell lines were associated with constitutive ATM activation.

**Figure S4:** ALT neuroblastoma cell lines were not exclusively associated with constitutive ATR and CHK1 activation in neuroblastoma cell lines.

**Figure S5.** ATM inhibitor AZD0156, reversed resistance to TMZ+SN-38 in p53 non-functional telomerase positive cells with induced telomere dysfunction.

**Figure S6.** ATM inhibitor AZD0156, reversed resistance to TMZ+SN-38 in p53 non-functional telomerase- positive cell line (SK-N-BE(2)) with induced ATM dependent TIFs by forced expression of TRF1-FOKI.

**Figure. S7.** ATM knockdown reduced C-circle content in ALT neuroblastoma cell lines.

**Figure S8.** AZD0156 showed heterogeneous response when combined with TMZ+SN-38 in telomerase positive cell lines.

**Figure S9.** The ATM inhibitor KU60019 sensitized ALT neuroblastoma cell lines to TMZ+SN- 38.

**Figure S10.** AZD0156 enhanced TMZ+SN-38 activity at dose as low as 6.25nM in ALT neuroblastoma *in vitro*.

**Figure S11.** AZD0156 synergized with TMZ+SN-38 in ALT neuroblastoma *in vitro*.

**Figure S12.** The ATM inhibitor AZD0156 enhanced induction of cleaved caspase 3 and cleaved PARP in response to TMZ+SN-38 in ALT cell lines.

**Figure S13.** ATM knockdown sensitized ALT neuroblastoma cell lines to TMZ+SN-38.

**Figure S14.** ATR inhibitor AZD6738 did not enhance activity of TMZ+SN-38 in ALT neuroblastoma cell lines.

**Figure S15.** Pharmacokinetic evaluation of unbound AZD0156 concentration *in vivo*.

**Figure S16.** Activity of TMZ+IRN +/- the ATM inhibitor AZD0156 combined with in ALT xenografts.

**Figure S17.** AZD0156 sequenced following treatment with TMZ+IRN was well tolerated *in vivo*.

**Table S1:** List of patient-derived neuroblastoma cell lines and patient-derived xenografts used in this study annotated by phase of therapy, sample type, disease stage, MYCN, TH mRNA, p53 status, ALT and telomerase status, and doubling time.

**Table S2:** List of somatic mutations identified for COG-N-515 and COG-N-512 using whole exome sequencing.

**Table S3:** Lists GSEA results for BIOCARTA gene sets comparing ALT versus telomerase positive non-*MYCN* amplified cell lines.

**Table S4:** Combination index values for 4 ALT neuroblastoma cell lines treated with TMZ+SN- 38+AZD0156.

**Table S5:** Pharmacokinetics of AZD0156 in swiss athymic nu/nu male mice.

## Supporting information

Koneru et al Supplemental Data

## Acknowledgments

We thank Children’s Oncology Group repository for providing cell lines and PDXs for this study. We thank the patients and their families for donating samples to enable this research. We thank Titia de Lange for providing pLPC-NMYC TRF2 deltaB deltaM, pLPC-NMYC-hTRF1 and pLPC-NMYC vector. We thank Dr. David Wheeler of the Human Genome Sequencing Center at Baylor College of Medicine, Houston Texas for the RNA sequencing and whole exome sequencing. We thank Tito Woodburn, Heather L. Davidson, Kristyn E. McCoy, and Jonas A. Nance for their efforts in establishing neuroblastoma models.

## Author contributions

B.K., C.P.R., A.F. came up with the concept. B.K. and C.P.R. wrote the manuscript. B.K., C.P.R., and S.T.D. designed the study. S.T.D. leads the Bioscience of ATM projects at AstraZeneca B.K., A.F., T.N., W.H.C., A.H., C.E., M.R.M. and T.B. performed experiments. J.W., A.S., V.P.R. and E.C. designed and acquired pharmacokinetic data *in vivo*. C.P.R. and S.T.D. supervised the study.

## Data and materials availability

Data and materials will be provided upon request. Requests for the data and materials should be submitted to C.P.R. as point of contact. Patient-derived cell lines and xenografts are freely available from the COG/ALSF Childhood Cancer Repository (www.CCcells.org).

## References

1. D. Hanahan, R. A. Weinberg, Hallmarks of cancer: the next generation. Cell 144, 646–674 (2011).

2. N. W. Kim, M. A. Piatyszek, K. R. Prowse, C. B. Harley, M. D. West, P. L. Ho, G. M. Coviello, W. E. Wright, S. L. Weinrich, J. W. Shay, Specific association of human telomerase activity with immortal cells and cancer. Science 266, 2011–2015 (1994).

3. C. M. Heaphy, A. P. Subhawong, S. M. Hong, M. G. Goggins, E. A. Montgomery, E. Gabrielson, G. J. Netto, J. I. Epstein, T. L. Lotan, W. H. Westra, M. Shih Ie, C. A. Iacobuzio-Donahue, A. Maitra, Q. K. Li, C. G. Eberhart, J. M. Taube, D. Rakheja, R. J. Kurman, T. C. Wu, R. B. Roden, P. Argani, A. M. De Marzo, L. Terracciano, M. Torbenson, A. K. Meeker, Prevalence of the alternative lengthening of telomeres telomere maintenance mechanism in human cancer subtypes. Am J Pathol 179, 1608–1615 (2011).

4. J. D. Henson, R. R. Reddel, Assaying and investigating Alternative Lengthening of Telomeres activity in human cells and cancers. FEBS Lett 584, 3800–3811 (2010).

5. M. A. Dunham, A. A. Neumann, C. L. Fasching, R. R. Reddel, Telomere maintenance by recombination in human cells. Nat Genet 26, 447–450 (2000).

6. J. A. Londono-Vallejo, H. Der-Sarkissian, L. Cazes, S. Bacchetti, R. R. Reddel, Alternative lengthening of telomeres is characterized by high rates of telomeric exchange. Cancer Res 64, 2324–2327 (2004).

7. T. M. Bryan, A. Englezou, J. Gupta, S. Bacchetti, R. R. Reddel, Telomere elongation in immortal human cells without detectable telomerase activity. EMBO J 14, 4240–4248 (1995).

8. T. R. Yeager, A. A. Neumann, A. Englezou, L. I. Huschtscha, J. R. Noble, R. R. Reddel, Telomerase-negative immortalized human cells contain a novel type of promyelocytic leukemia (PML) body. Cancer Res 59, 4175–4179 (1999).

9. J. D. Henson, Y. Cao, L. I. Huschtscha, A. C. Chang, A. Y. Au, H. A. Pickett, R. R. Reddel, DNA C-circles are specific and quantifiable markers of alternative-lengthening-of- telomeres activity. Nat Biotechnol 27, 1181–1185 (2009).

10. R. L. Dilley, P. Verma, N. W. Cho, H. D. Winters, A. R. Wondisford, R. A. Greenberg, Break-induced telomere synthesis underlies alternative telomere maintenance. Nature 539, 54–58 (2016).

11. F. M. Roumelioti, S. K. Sotiriou, V. Katsini, M. Chiourea, T. D. Halazonetis, S. Gagos, Alternative lengthening of human telomeres is a conservative DNA replication process with features of break-induced replication. EMBO Rep 17, 1731–1737 (2016).

12. J. Min, W. E. Wright, J. W. Shay, Alternative Lengthening of Telomeres Mediated by Mitotic DNA Synthesis Engages Break-Induced Replication Processes. Mol Cell Biol 37, (2017).

13. R. L. Flynn, K. E. Cox, M. Jeitany, H. Wakimoto, A. R. Bryll, N. J. Ganem, F. Bersani, J. R. Pineda, M. L. Suva, C. H. Benes, D. A. Haber, F. D. Boussin, L. Zou, Alternative lengthening of telomeres renders cancer cells hypersensitive to ATR inhibitors. Science 347, 273–277 (2015).

14. K. I. Deeg, I. Chung, C. Bauer, K. Rippe, Cancer Cells with Alternative Lengthening of Telomeres Do Not Display a General Hypersensitivity to ATR Inhibition. Frontiers in Oncology 6, (2016).

15. S. L. George, V. Parmar, F. Lorenzi, L. V. Marshall, Y. Jamin, E. Poon, P. Angelini, L. Chesler, Novel therapeutic strategies targeting telomere maintenance mechanisms in high- risk neuroblastoma. J Exp Clin Cancer Res 39, 78 (2020).

16. R. Lu, J. J. O’Rourke, A. P. Sobinoff, J. A. M. Allen, C. B. Nelson, C. G. Tomlinson, M. Lee, R. R. Reddel, A. J. Deans, H. A. Pickett, The FANCM-BLM-TOP3A-RMI complex suppresses alternative lengthening of telomeres (ALT). Nat Commun 10, 2252 (2019).

17. Y. Wang, J. Yang, A. T. Wild, W. H. Wu, R. Shah, C. Danussi, G. J. Riggins, K. Kannan, E. P. Sulman, T. A. Chan, J. T. Huse, G-quadruplex DNA drives genomic instability and represents a targetable molecular abnormality in ATRX-deficient malignant glioma. Nat Commun 10, 943 (2019).

18. M. Han, C. E. Napier, S. Frolich, E. Teber, T. Wong, J. R. Noble, E. H. Y. Choi, R. D. Everett, A. J. Cesare, R. R. Reddel, Synthetic lethality of cytolytic HSV-1 in cancer cells with ATRX and PML deficiency. J Cell Sci 132, (2019).

19. X. H. Zheng, X. Nie, Y. Fang, Z. Zhang, Y. Xiao, Z. Mao, H. Liu, J. Ren, F. Wang, L. Xia, J. Huang, Y. Zhao, A Cisplatin Derivative Tetra-Pt(bpy) as an Oncotherapeutic Agent for Targeting ALT Cancer. J Natl Cancer Inst 109, (2017).

20. R. A. Dagg, H. A. Pickett, A. A. Neumann, C. E. Napier, J. D. Henson, E. T. Teber, J. W. Arthur, C. P. Reynolds, J. Murray, M. Haber, A. P. Sobinoff, L. M. S. Lau, R. R. Reddel, Extensive Proliferation of Human Cancer Cells with Ever-Shorter Telomeres. Cell Rep 19, 2544–2556 (2017).

21. N. Viceconte, M. S. Dheur, E. Majerova, C. E. Pierreux, J. F. Baurain, N. van Baren, A. Decottignies, Highly Aggressive Metastatic Melanoma Cells Unable to Maintain Telomere Length. Cell Rep 19, 2529–2543 (2017).

22. M. A. Smith, S. F. Altekruse, P. C. Adamson, G. H. Reaman, N. L. Seibel, Declining childhood and adolescent cancer mortality. Cancer 120, 2497–2506 (2014).

23. M. A. Smith, N. L. Seibel, S. F. Altekruse, L. A. Ries, D. L. Melbert, M. O’Leary, F. O. Smith, G. H. Reaman, Outcomes for children and adolescents with cancer: challenges for the twenty-first century. J Clin Oncol 28, 2625–2634 (2010).

24. J. M. Maris, Recent advances in neuroblastoma. N Engl J Med 362, 2202–2211 (2010).

25. J. R. Park, S. G. Kreissman, W. B. London, A. Naranjo, S. L. Cohn, M. D. Hogarty, S. C. Tenney, D. Haas-Kogan, P. J. Shaw, J. M. Kraveka, S. S. Roberts, J. D. Geiger, J. J. Doski, S. D. Voss, J. M. Maris, S. A. Grupp, L. Diller, Effect of Tandem Autologous Stem Cell Transplant vs Single Transplant on Event-Free Survival in Patients With High-Risk Neuroblastoma: A Randomized Clinical Trial. JAMA 322, 746–755 (2019).

26. A. L. Yu, A. L. Gilman, M. F. Ozkaynak, W. B. London, S. G. Kreissman, H. X. Chen, M. Smith, B. Anderson, J. G. Villablanca, K. K. Matthay, H. Shimada, S. A. Grupp, R. Seeger, C. P. Reynolds, A. Buxton, R. A. Reisfeld, S. D. Gillies, S. L. Cohn, J. M. Maris, P. M. Sondel, Anti-GD2 antibody with GM-CSF, interleukin-2, and isotretinoin for neuroblastoma. N Engl J Med 363, 1324–1334 (2010).

27. S. Ackermann, M. Cartolano, B. Hero, A. Welte, Y. Kahlert, A. Roderwieser, C. Bartenhagen, E. Walter, J. Gecht, L. Kerschke, R. Volland, R. Menon, J. M. Heuckmann, M. Gartlgruber, S. Hartlieb, K. O. Henrich, K. Okonechnikov, J. Altmuller, P. Nurnberg, S. Lefever, B. de Wilde, F. Sand, F. Ikram, C. Rosswog, J. Fischer, J. Theissen, F. Hertwig, A. D. Singhi, T. Simon, W. Vogel, S. Perner, B. Krug, M. Schmidt, S. Rahmann, V. Achter, U. Lang, C. Vokuhl, M. Ortmann, R. Buttner, A. Eggert, F. Speleman, R. J. O’Sullivan, R. K. Thomas, F. Berthold, J. Vandesompele, A. Schramm, F. Westermann, J. H. Schulte, M. Peifer, M. Fischer, A mechanistic classification of clinical phenotypes in neuroblastoma. Science 362, 1165–1170 (2018).

28. A. Roderwieser, F. Sand, E. Walter, J. Fischer, J. Gecht, C. Bartenhagen, S. Ackermann, F. Otte, A. Welte, Y. Kahlert, D. Lieberz, F. Hertwig, H. C. Reinhardt, T. Simon, M. Peifer, M. Ortmann, R. Büttner, B. Hero, R. J. O’Sullivan, F. Berthold, M. Fischer, Telomerase Is a Prognostic Marker of Poor Outcome and a Therapeutic Target in Neuroblastoma. JCO Precision Oncology, 1–20 (2019).

29. B. Koneru, G. Lopez, A. Farooqi, K. L. Conkrite, T. H. Nguyen, S. J. Macha, A. Modi, J. L. Rokita, E. Urias, A. Hindle, H. Davidson, K. McCoy, J. Nance, V. Yazdani, M. S. Irwin, S. Yang, D. A. Wheeler, J. M. Maris, S. J. Diskin, C. P. Reynolds, Telomere Maintenance Mechanisms Define Clinical Outcome in High-Risk Neuroblastoma. Cancer Res 80, 2663–2675 (2020).

30. N. K. Cheung, J. Zhang, C. Lu, M. Parker, A. Bahrami, S. K. Tickoo, A. Heguy, A. S. Pappo, S. Federico, J. Dalton, I. Y. Cheung, L. Ding, R. Fulton, J. Wang, X. Chen, J. Becksfort, J. Wu, C. A. Billups, D. Ellison, E. R. Mardis, R. K. Wilson, J. R. Downing, M. Dyer, Association of age at diagnosis and genetic mutations in patients with neuroblastoma. JAMA 307, 1062–1071 (2012).

31. J. J. Molenaar, J. Koster, D. A. Zwijnenburg, P. van Sluis, L. J. Valentijn, I. van der Ploeg, Hamdi, J. van Nes, B. A. Westerman, J. van Arkel, M. E. Ebus, F. Haneveld, A. Lakeman, L. Schild, P. Molenaar, P. Stroeken, M. M. van Noesel, I. Ora, E. E. Santo, H. Caron, E. M. Westerhout, R. Versteeg, Sequencing of neuroblastoma identifies chromothripsis and defects in neuritogenesis genes. Nature 483, 589–593 (2012).

32. S. Kurihara, E. Hiyama, Y. Onitake, E. Yamaoka, K. Hiyama, Clinical features of ATRX or DAXX mutated neuroblastoma. J Pediatr Surg 49, 1835–1838 (2014).

33. V. Hakin-Smith, D. A. Jellinek, D. Levy, T. Carroll, M. Teo, W. R. Timperley, M. J. McKay, R. R. Reddel, J. A. Royds, Alternative lengthening of telomeres and survival in patients with glioblastoma multiforme. Lancet 361, 836–838 (2003).

34. J. D. Henson, J. A. Hannay, S. W. McCarthy, J. A. Royds, T. R. Yeager, R. A. Robinson, S. B. Wharton, D. A. Jellinek, S. M. Arbuckle, J. Yoo, B. G. Robinson, D. L. Learoyd, P. D. Stalley, S. F. Bonar, D. Yu, R. E. Pollock, R. R. Reddel, A robust assay for alternative lengthening of telomeres in tumors shows the significance of alternative lengthening of telomeres in sarcomas and astrocytomas. Clin Cancer Res 11, 217–225 (2005).

35. J. Y. Liau, J. H. Tsai, Y. M. Jeng, J. C. Lee, H. H. Hsu, C. Y. Yang, Leiomyosarcoma with alternative lengthening of telomeres is associated with aggressive histologic features, loss of ATRX expression, and poor clinical outcome. Am J Surg Pathol 39, 236–244 (2015).

36. R. T. Lawlor, N. Veronese, A. Pea, A. Nottegar, L. Smith, C. Pilati, J. Demurtas, M. Fassan, L. Cheng, C. Luchini, Alternative lengthening of telomeres (ALT) influences survival in soft tissue sarcomas: a systematic review with meta-analysis. BMC Cancer 19, 232 (2019).

37. J. Y. Kim, J. A. Brosnan-Cashman, S. An, S. J. Kim, K. B. Song, M. S. Kim, M. J. Kim, D. W. Hwang, A. K. Meeker, E. Yu, S. C. Kim, R. H. Hruban, C. M. Heaphy, S. M. Hong, Alternative Lengthening of Telomeres in Primary Pancreatic Neuroendocrine Tumors Is Associated with Aggressive Clinical Behavior and Poor Survival. Clin Cancer Res 23, 1598–1606 (2017).

38. Y. Onitake, E. Hiyama, N. Kamei, H. Yamaoka, T. Sueda, K. Hiyama, Telomere biology in neuroblastoma: telomere binding proteins and alternative strengthening of telomeres. J Pediatr Surg 44, 2258–2266 (2009).

39. A. S. Farooqi, R. A. Dagg, L. M. Choi, J. W. Shay, C. P. Reynolds, L. M. Lau, Alternative lengthening of telomeres in neuroblastoma cell lines is associated with a lack of MYCN genomic amplification and with p53 pathway aberrations. J Neurooncol 119, 17–26 (2014).

40. S. Matsuoka, B. A. Ballif, A. Smogorzewska, E. R. McDonald, 3rd, K. E. Hurov, J. Luo, C. E. Bakalarski, Z. Zhao, N. Solimini, Y. Lerenthal, Y. Shiloh, S. P. Gygi, S. J. Elledge, ATM and ATR substrate analysis reveals extensive protein networks responsive to DNA damage. Science 316, 1160–1166 (2007).

41. N. Keshelava, J. J. Zuo, P. Chen, S. N. Waidyaratne, M. C. Luna, C. J. Gomer, T. J. Triche, C. P. Reynolds, Loss of p53 function confers high-level multidrug resistance in neuroblastoma cell lines. Cancer Res 61, 6185–6193 (2001).

42. H. Takai, A. Smogorzewska, T. de Lange, DNA damage foci at dysfunctional telomeres. Curr Biol 13, 1549–1556 (2003).

43. E. L. Denchi, T. de Lange, Protection of telomeres through independent control of ATM and ATR by TRF2 and POT1. Nature 448, 1068–1071 (2007).

44. A. J. Cesare, Z. Kaul, S. B. Cohen, C. E. Napier, H. A. Pickett, A. A. Neumann, R. R. Reddel, Spontaneous occurrence of telomeric DNA damage response in the absence of chromosome fusions. Nat Struct Mol Biol 16, 1244–1251 (2009).

45. H. Jiang, H. C. Reinhardt, J. Bartkova, J. Tommiska, C. Blomqvist, H. Nevanlinna, J. Bartek, M. B. Yaffe, M. T. Hemann, The combined status of ATM and p53 link tumor development with therapeutic response. Genes Dev 23, 1895–1909 (2009).

46. B. van Steensel, A. Smogorzewska, T. de Lange, TRF2 protects human telomeres from end-to-end fusions. Cell 92, 401–413 (1998).

47. Y. Doksani, T. de Lange, Telomere-Internal Double-Strand Breaks Are Repaired by Homologous Recombination and PARP1/Lig3-Dependent End-Joining. Cell Rep 17, 1646–1656 (2016).

48. K. G. Pike, B. Barlaam, E. Cadogan, A. Campbell, Y. Chen, N. Colclough, N. L. Davies, C. de-Almeida, S. L. Degorce, M. Didelot, A. Dishington, R. Ducray, S. T. Durant, L. A. Hassall, J. Holmes, G. D. Hughes, P. A. MacFaul, K. R. Mulholland, T. M. McGuire, G. Ouvry, M. Pass, G. Robb, N. Stratton, Z. Wang, J. Wilson, B. Zhai, K. Zhao, N. Al-Huniti, The Identification of Potent, Selective, and Orally Available Inhibitors of Ataxia Telangiectasia Mutated (ATM) Kinase: The Discovery of AZD0156 (8-{6-[3- (Dimethylamino)propoxy]pyridin-3-yl}-3-methyl-1-(tetrahydro-2 H-pyran-4-yl)-1,3- dihydro-2 H-imidazo[4,5- c]quinolin-2-one). J Med Chem 61, 3823–3841 (2018).

49. Z. A. Qadeer, D. Valle-Garcia, D. Hasson, Z. Sun, A. Cook, C. Nguyen, A. Soriano, A. Ma, L. M. Griffiths, M. Zeineldin, D. Filipescu, L. Jubierre, A. Chowdhury, O. Deevy, X. Chen, D. B. Finkelstein, A. Bahrami, E. Stewart, S. Federico, S. Gallego, F. Dekio, M. Fowkes, D. Meni, J. M. Maris, W. A. Weiss, S. S. Roberts, N. V. Cheung, J. Jin, M. F. Segura, M. A. Dyer, E. Bernstein, ATRX In-Frame Fusion Neuroblastoma Is Sensitive to EZH2 Inhibition via Modulation of Neuronal Gene Signatures. Cancer Cell 36, 512–527 e519 (2019).

50. A. R. Gocha, J. Harris, J. Groden, Alternative mechanisms of telomere lengthening: permissive mutations, DNA repair proteins and tumorigenic progression. Mutat Res 743- 744, 142-150 (2013).

51. Y. J. Chen, V. Hakin-Smith, M. Teo, G. E. Xinarianos, D. A. Jellinek, T. Carroll, D. McDowell, M. R. MacFarlane, R. Boet, B. C. Baguley, A. W. Braithwaite, R. R. Reddel, J. A. Royds, Association of mutant TP53 with alternative lengthening of telomeres and favorable prognosis in glioma. Cancer Res 66, 6473–6476 (2006).

52. M. H. Kang, M. A. Smith, C. L. Morton, N. Keshelava, P. J. Houghton, C. P. Reynolds, National Cancer Institute pediatric preclinical testing program: model description for in vitro cytotoxicity testing. Pediatric blood & cancer 56, 239–249 (2011).

53. T. H. Nguyen, B. Koneru, S.-J. Wei, W. H. Chen, M. R. Makena, E. Urias, M. H. Kang, C. P. Reynolds, Fenretinide via NOXA Induction, Enhanced Activity of the BCL-2 Inhibitor Venetoclax in High BCL-2–Expressing Neuroblastoma Preclinical Models. Molecular Cancer Therapeutics 18, 2270–2282 (2019).

54. M. R. Makena, B. Koneru, T. H. Nguyen, M. H. Kang, C. P. Reynolds, Reactive Oxygen Species-Mediated Synergism of Fenretinide and Romidepsin in Preclinical Models of T-cell Lymphoid Malignancies. Mol Cancer Ther 16, 649–661 (2017).

55. B. M. Gyori, G. Venkatachalam, P. S. Thiagarajan, D. Hsu, M. V. Clement, OpenComet: an automated tool for comet assay image analysis. Redox Biol 2, 457–465 (2014).

56. M. M. Tomayko, C. P. Reynolds, Determination of subcutaneous tumor size in athymic (nude) mice. Cancer Chemother Pharmacol 24, 148–154 (1989).

57. J. L. Rokita, K. S. Rathi, M. F. Cardenas, K. A. Upton, J. Jayaseelan, K. L. Cross, J. Pfeil, L. E. Egolf, G. P. Way, A. Farrel, N. M. Kendsersky, K. Patel, K. S. Gaonkar, A. Modi, E. R. Berko, G. Lopez, Z. Vaksman, C. Mayoh, J. Nance, K. McCoy, M. Haber, K. Evans, H. McCalmont, K. Bendak, J. W. Bohm, G. M. Marshall, V. Tyrrell, K. Kalletla, F. K. Braun, L. Qi, Y. Du, H. Zhang, H. B. Lindsay, S. Zhao, J. Shu, P. Baxter, C. Morton, D. Kurmashev, S. Zheng, Y. Chen, J. Bowen, A. C. Bryan, K. M. Leraas, S. E. Coppens, H. Doddapaneni, Z. Momin, W. Zhang, G. I. Sacks, L. S. Hart, K. Krytska, Y. P. Mosse, G. J. Gatto, Y. Sanchez, C. S. Greene, S. J. Diskin, O. M. Vaske, D. Haussler, J. M. Gastier-Foster, E. A. Kolb, R. Gorlick, X. N. Li, C. P. Reynolds, R. T. Kurmasheva, P. J. Houghton, M. A. Smith, R. B. Lock, P. Raman, D. A. Wheeler, J. M. Maris, Genomic Profiling of Childhood Tumor Patient-Derived Xenograft Models to Enable Rational Clinical Trial Design. Cell Rep 29, 1675–1689 e1679 (2019).

58. A. Hindle, B. Koneru, M. R. Makena, L. Lopez-Barcons, W. H. Chen, T. H. Nguyen, C. P. Reynolds, The O6-methyguanine-DNA methyltransferase inhibitor O6-benzylguanine enhanced activity of temozolomide + irinotecan against models of high-risk neuroblastoma. Anticancer Drugs 32, 233–247 (2021).

59. W. Cai, N. V. Maldonado, W. Cui, N. Harutyunyan, L. Ji, R. Sposto, C. P. Reynolds, N. Keshelava, Activity of irinotecan and temozolomide in the presence of O6-methylguanine-DNA methyltransferase inhibition in neuroblastoma pre-clinical models. Br J Cancer 103, 1369–1379 (2010).

60. P. J. Houghton, C. L. Morton, C. Tucker, D. Payne, E. Favours, C. Cole, R. Gorlick, E. A. Kolb, W. Zhang, R. Lock, H. Carol, M. Tajbakhsh, C. P. Reynolds, J. M. Maris, J. Courtright, S. T. Keir, H. S. Friedman, C. Stopford, J. Zeidner, J. Wu, T. Liu, C. A. Billups, J. Khan, S. Ansher, J. Zhang, M. A. Smith, The pediatric preclinical testing program: description of models and early testing results. Pediatr Blood Cancer 49, 928–940 (2007).

