## Supplementary material for "ALT Neuroblastoma Chemoresistance due to ATM Activation by Telomere Dysfunction is Reversible with the ATM Inhibitor AZD0156": Koneru et al Supplemental Data

##### Figure S1

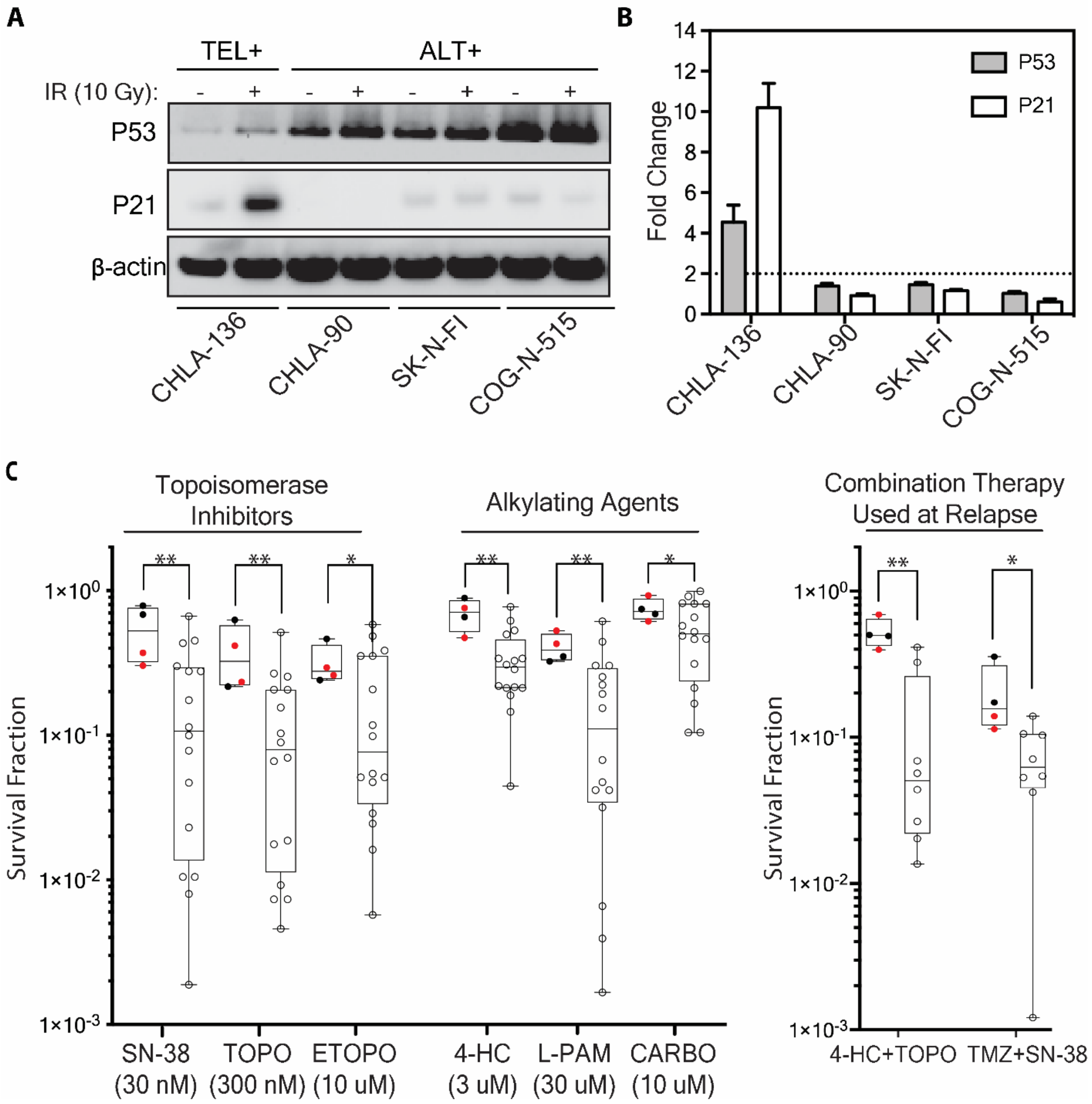

**Figure S1. ALT neuroblastoma models were associated with resistance to DNA damaging agents irrespective of p53 functionality.** (A) Immunoblotting for p53, p21 and beta-actin in a *p53* wild-type telomerase positive (CHLA-136) and 3 ALT+ (CHLA-90, SK-N-FI and COG-N-515) neuroblastoma cell lines treated with or without irradiation (10 Grays). Immunoblotting was performed at 16 hours post irradiation. (B) Bar graph shows quantification of fold-change in p53 and p21 protein levels post irradiation, in the same cells as shown in (A). Horizontal dotted line is used as a cutoff for determining p53 functionality. (C) (left panel) Box plot compares survival fraction of ALT (n=4) versus non-ALT (n=16) p53 non-functional neuroblastoma cell lines treated with clinically achievable doses of 7-ethyl-10-hydroxycamptothecin (SN-38; an active metabolite of irinotecan), topotecan (TOPO), etoposide (ETOPO), 4-hydroperoxy cyclophosphamide (4-HC; an active metabolite of cyclophosphamide), melphalan (L-PAM), or carboplatin (CARBO). (right panel) Box plot compares survival fraction of ALT (n=4) versus non-ALT (n=8) p53 non-functional neuroblastoma cell lines treated with 4-HC (1 μM) + TOPO (100 nM) or temozolomide (TMZ; 30 μM) + SN-38 (3 nM). ALT positive cell lines COG-N-512 and COG-N-515 that were established from same patient are highlighted in red. Survival fraction between ALT versus non-ALT cell lines is compared using Wilcoxon-rank sum test. *: $P<0.05$, **: $P<0.01$, ns: not significant.

**Figure S2**

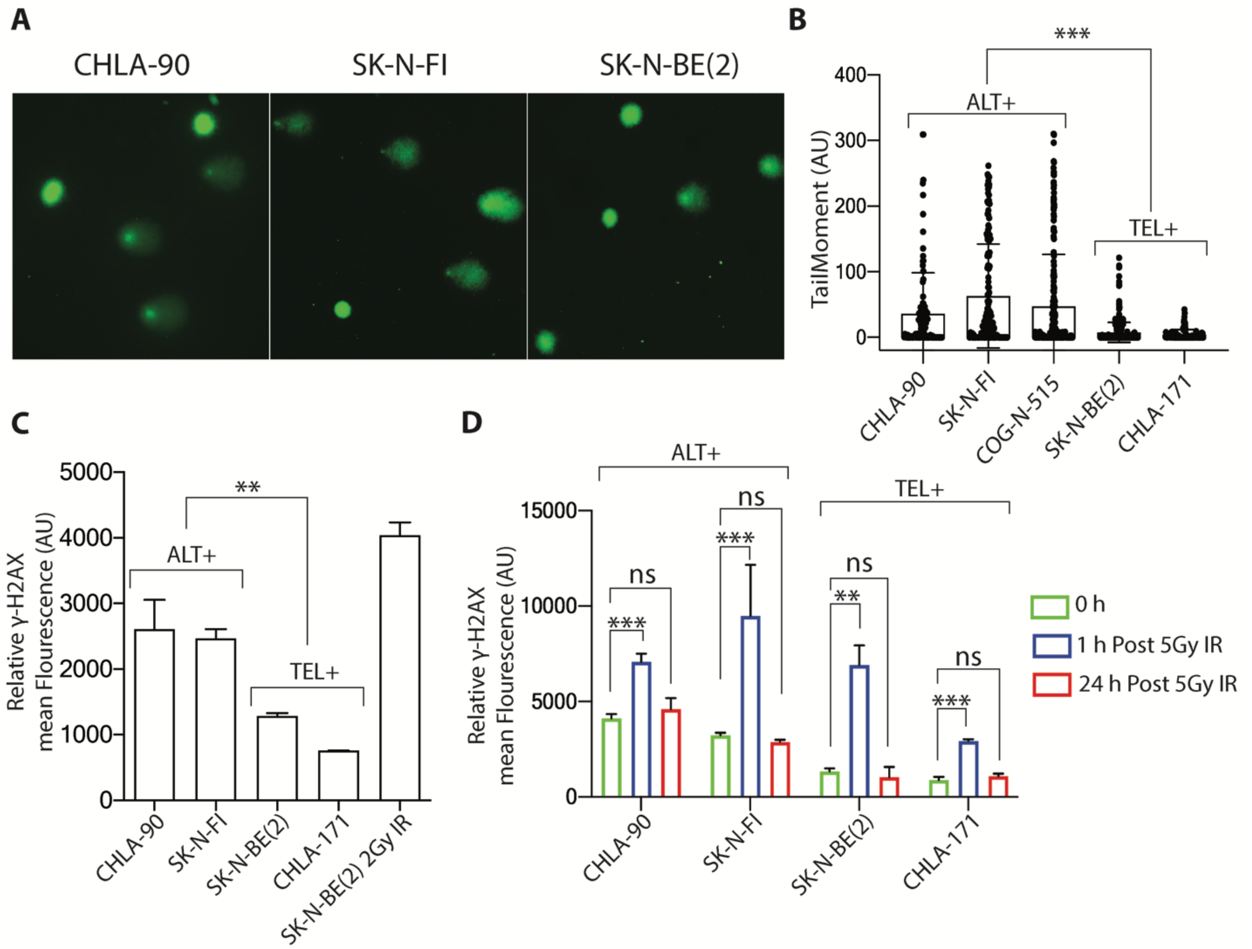

**Figure S2: ALT neuroblastoma was associated with constitutive DNA damage.** (A) Representative images of comet assay in 2 ALT cell lines (CHLA-90 and SK-N-FI) and a p53 non-functional telomerase-positive cell lines (SK-N-BE(2)) (B) Box plot shows tail moment measured on >100 comets in 2 ALT cell lines (CHLA-90 and SK-N-FI) vs 2 p53 non-functional telomerase-positive cell lines (SK-N-BE(2) and CHLA-171). Tail moment was measured using open comet plugin in Fiji software. Statistical significance was measured using Wilcoxon-rank sum test. (C) Graph shows normalized γ-H2AX mean fluorescence intensity measured by flow cytometry in 2 ALT cell lines (CHLA-90 and SK-N-FI) and 2 p53 non-functional telomerase-positive cell lines (SK-N-BE(2) and CHLA-171). SK-N-BE(2) cells at 1 hour post 2 Grays of irradiation are used as a positive control. (B) Bar graph shows normalized γ-H2AX mean fluorescence intensity measured by flow cytometry at 0-, 1- and 24-hours post 5 Grays of irradiation (IR) in same cells as in A. Two-tailed t-test was used to calculate statistical significance in D. ***: $P$<0.001, **: $P$<0.01, *: $P$<0.05, ns: not significant.

**Figure S3**

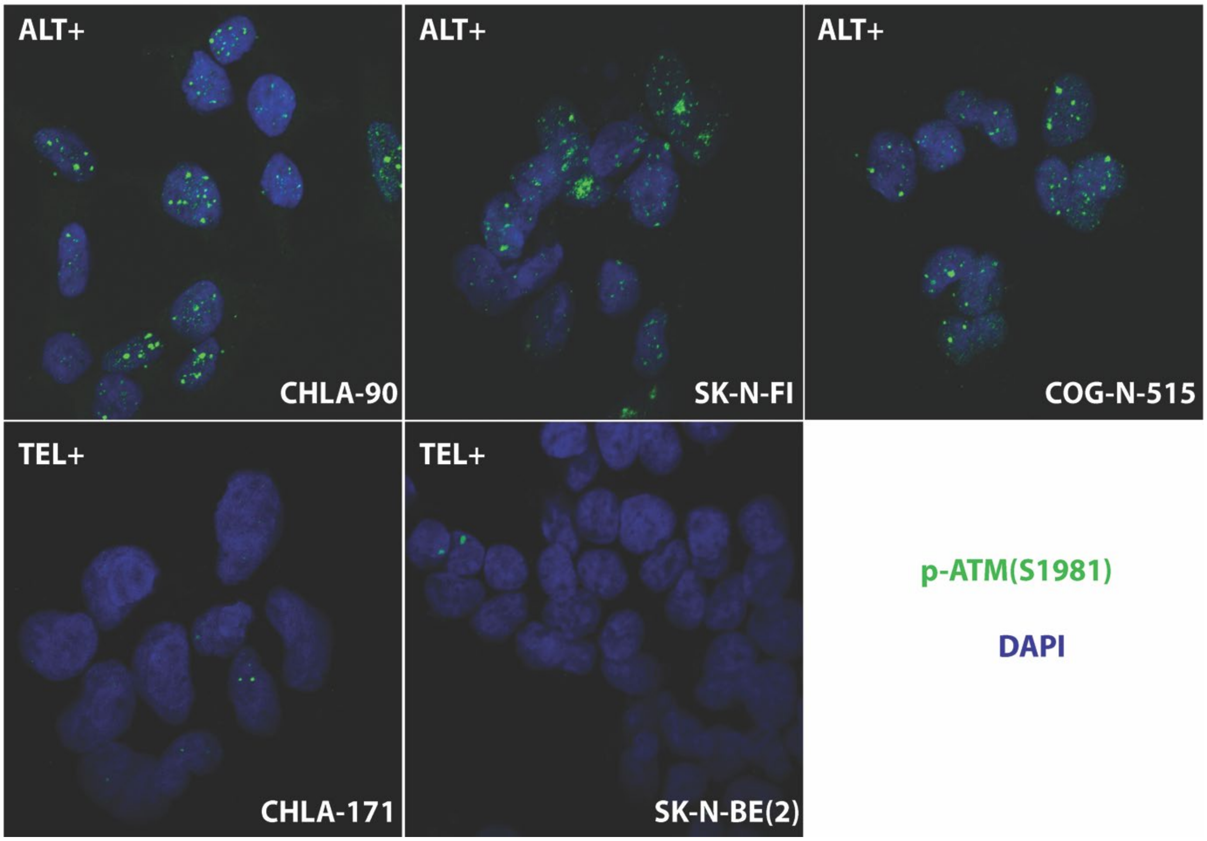

**Figure S3: ALT neuroblastoma cell lines were associated with constitutive ATM activation.** Representative images of Immunofluorescence staining for p-ATM (S1981) (green) in 3 ALT (CHLA-90, SK-N-FI and COG-N-515) and 2 telomerase-positive (CHLA-171 and SK-N-BE(2)) neuroblastoma cell lines. Nuclei were stained with DAPI (blue).

#### Figure S4

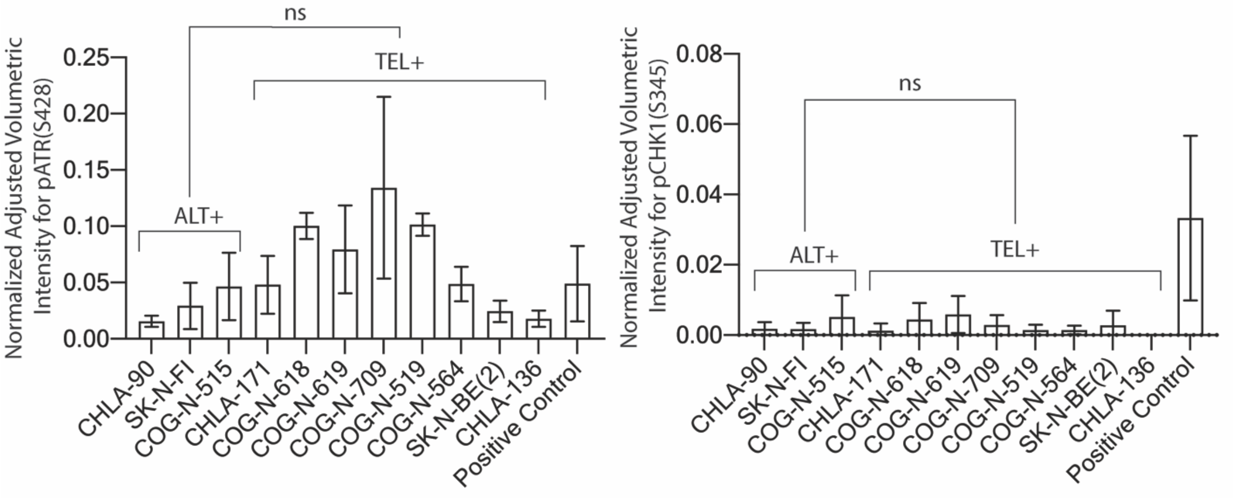

**Figure S4: ALT neuroblastoma cell lines were not exclusively associated with constitutive ATR and CHK1 activation in neuroblastoma cell lines.** (left panel) Bar graph shows quantification of immunoblot for pATR(S248) and (Right panel) pCHK1(S345) in 3 ALT and 8 telomerase-positive cell lines from the immunoblot in Figure 2D and its replicates. CHLA-136 twelve hours post-irradiation with x-rays (10 Grays) served as a positive control. The bars represent means with SDs from three experimental replicates. Protein quantification was compared by Wilcoxon-rank sum test. ns: not significant.

**Figure S5**

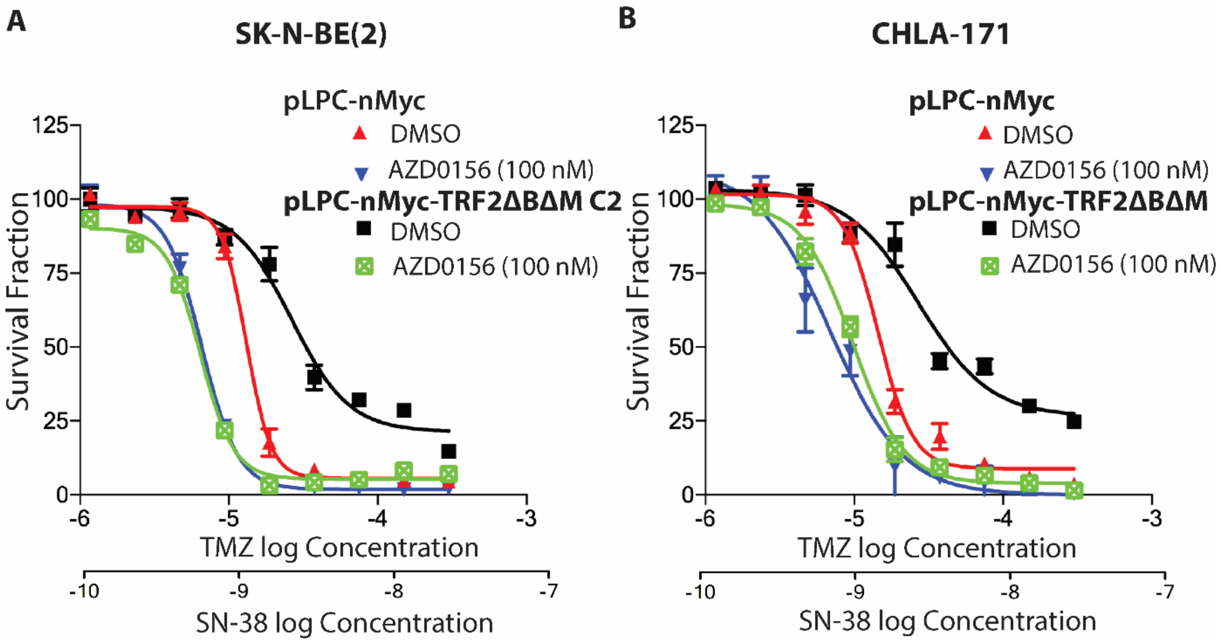

**Figure S5: ATM inhibitor AZD0156, reversed resistance to TMZ+SN-38 in p53 non-functional telomerase-positive cells with induced ATM dependent TIFs by forced expression of TRF2ΔBΔM.** DIMSCAN cytotoxic assay curves in response to TMZ+SN-38 +/- the ATM inhibitor AZD0156 in (A) SK-N-BE(2), and (B) CHLA-171 transduced with pLPC-nMyc or pLPC-nMyc-TRF2ΔBΔM. Clone 2 of SK-N-BE(2) transduced with pLPC-nMyc-TRF2ΔBΔM is shown in this figure, clone 1 is shown in Figure 4.

#### Figure S6

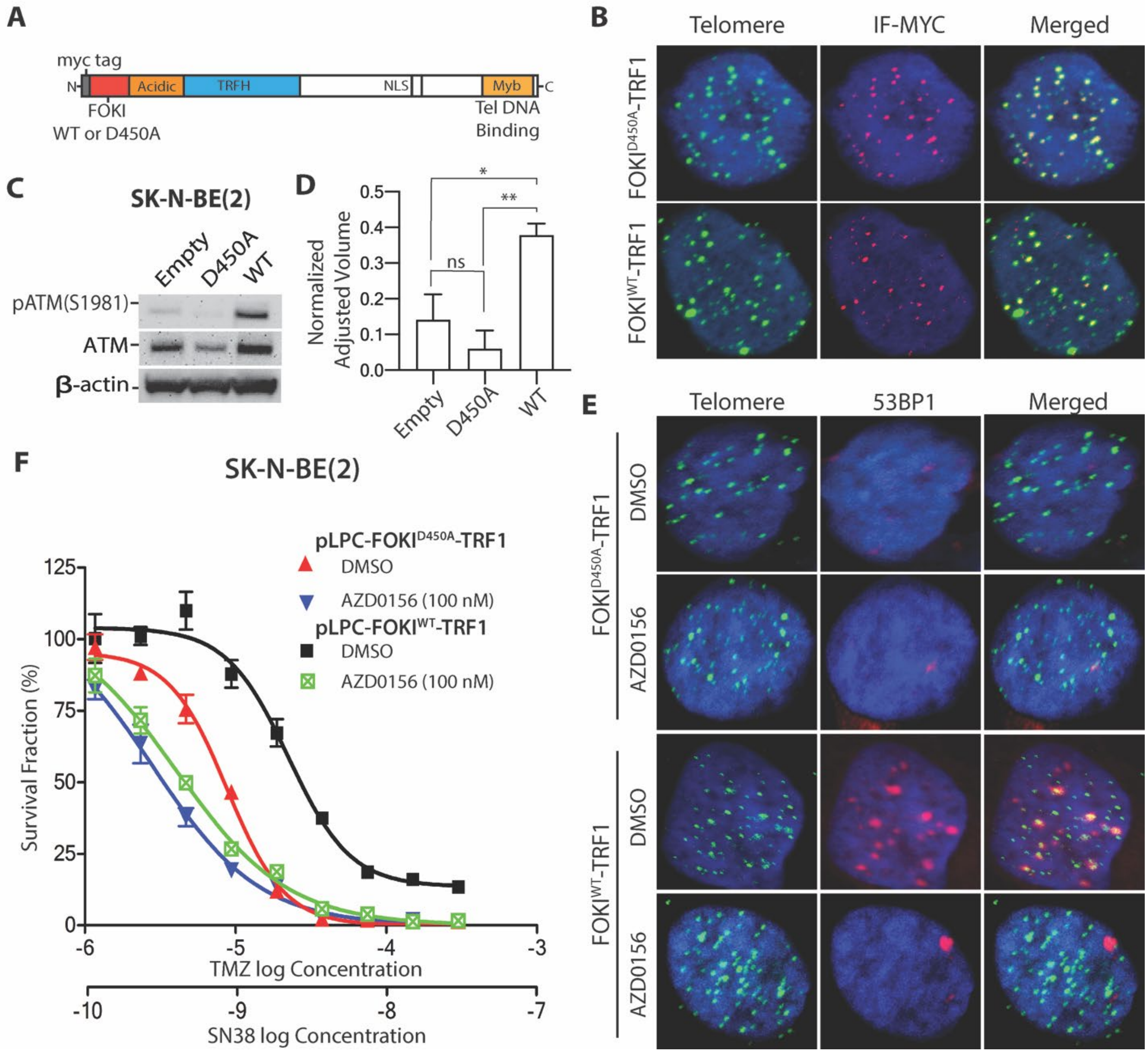

**Figure S6: ATM inhibitor AZD0156, reversed resistance to TMZ+SN-38 in p53 non-functional telomerase-positive cell line (SK-N-BE(2)) with induced ATM dependent TIFs by forced expression of TRF1-FOKI.** (A) Schematics of TRF1 fused with wild-type FOKI or D450A FOKI. (B) Representative images of Immunofluorescence (IF) staining for myc-tag (red) in combination with telomere fluorescent *in situ* hybridization (FISH) with a $[TTAGGG]_3$ probe (green) in SK-N-BE (2) cells transduced with pLPC-Myc-FOKI$^{D450A}$-TRF1 (nuclease dead mutant control) or pLPC-Myc-FOKI$^{WT}$-TRF1. Image shows co-localization of myc tagged protein with telomeres. (C) Immunoblot for pATM(S1981), ATM and β-actin in the telomerase-positive p53 non-functional cell line SK-N-BE(2) transduced with pLPC-Myc empty vector, pLPC-nMyc-FOKI$^{D450A}$-TRF1, or pLPC-nMyc-FOKI$^{WT}$-TRF1. (D) Bar graph shows protein quantification of normalized pATM(S1981) in C and its replicates. The bars represent means with SDs from three experimental replicates. (E) Representative images of TIF analysis using IF-FISH in same cells as in B +/-ATM inhibitor (AZD0156). 53BP1 was detected by IF (red) and telomeres by FISH with a $[TTAGGG]_3$ probe (green). (F) DIMSCAN cytotoxicity assay curves in response to TMZ+SN-38 +/- AZD0156 in same cells as shown in B.

**Figure S7**

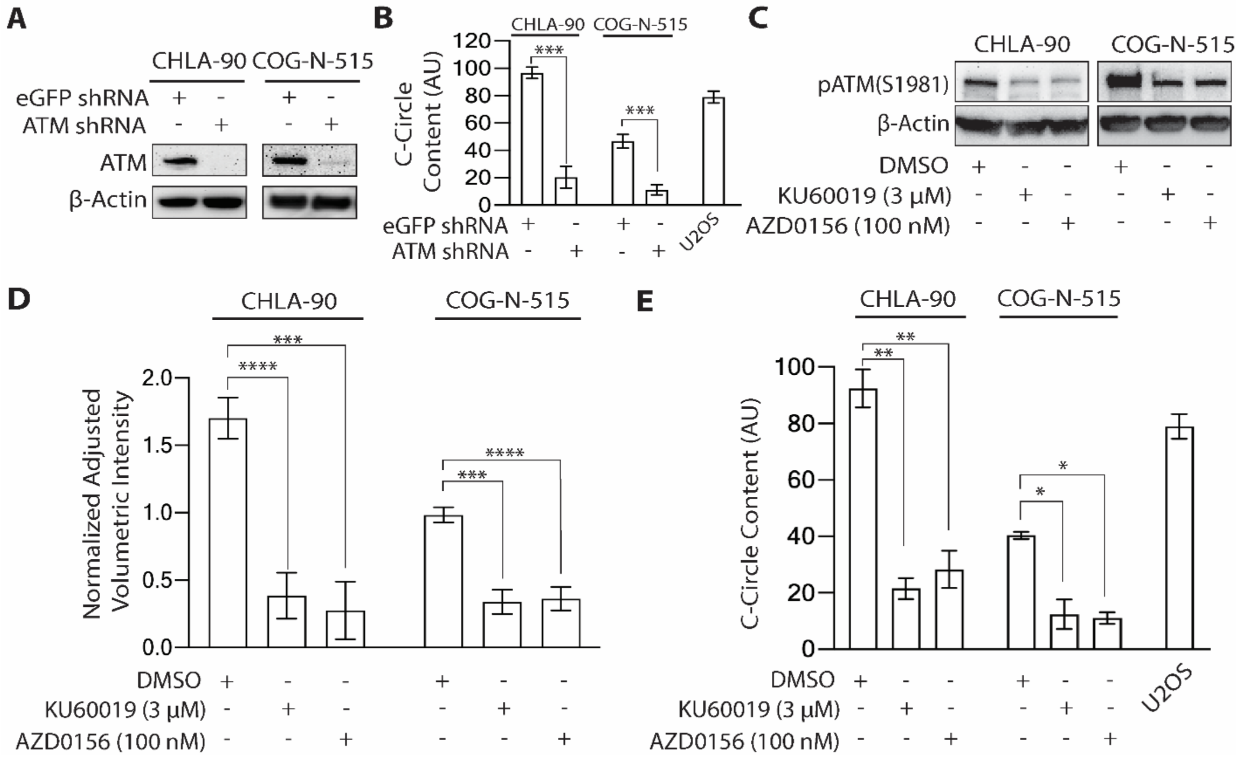

**Figure S7. ATM knockdown reduced C-circle content in ALT neuroblastoma cell lines.** (A) Immunoblotting for ATM and β-Actin in neuroblastoma ALT cell lines (CHLA-90 and COG-N-515) transduced with either eGFP or ATM shRNA. (B) Bar graph shows C-circle content in CHLA-90 and COG-N-515 transduced with either eGFP or ATM shRNA. C-circle content was measured using qPCR. ALT+ U2OS cell line was used as a positive control (C) Immunoblotting for pATM(S1981) and β-Actin in neuroblastoma ALT cell lines (CHLA-90 and COG-N-515) treated with DMSO, ATM inhibitors Ku60019 or ATM inhibitor AZD0156. (D) Normalized quantification of pATM(S1981) in same cells as in C and its replicates. (E) Bar graph shows C-circle content in CHLA-90 and COG-N-515 treated with DMSO, KU60019 or AZD0156. ALT+ U2OS cell line was used as a positive control. The bars represent means with SDs from three experimental replicates. Statistical significance was calculated using two-tailed t-test for D and E. ****: *P*<0.0001, ***: *P*<0.001, **: *P*<0.01, *: *P*<0.05, ns: not significant.

**Figure S8**

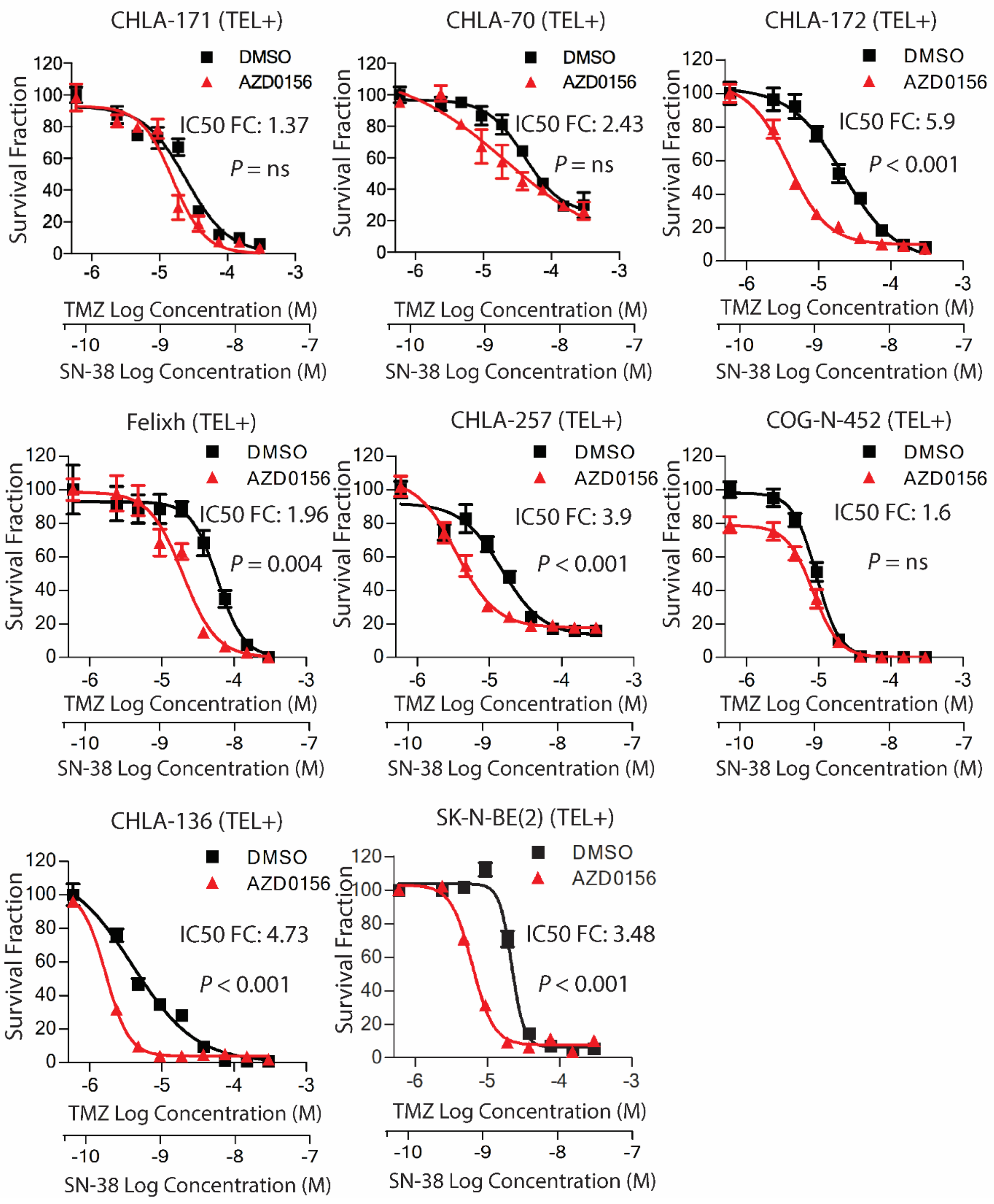

**Figure S8. AZD0156 showed heterogeneous response when combined with TMZ+SN-38 in telomerase positive cell lines.** DIMSCAN cytotoxicity assay curves in additional telomerase-positive neuroblastoma cell lines (n=8) following treatment with TMZ+SN-38 +/- AZD0156 (100 nM). Statistical significance for dose response curves was calculated using two-way ANOVA test. ns: not significant. IC50 FC: The half maximal inhibitory concentration fold change.

**Figure S9**

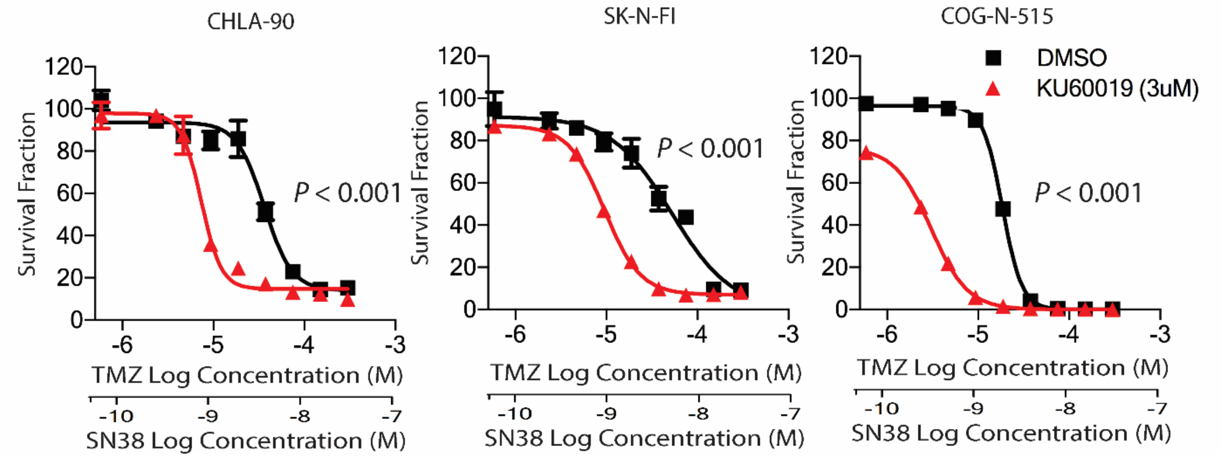

**Figure S9. The ATM inhibitor KU60019 sensitized ALT neuroblastoma cell lines to TMZ+SN-38.** DIMSCAN cytotoxicity assay curves in 3 ALT cell lines following treatment with TMZ+SN-38 +/- KU60019 (3 μM). Statistical significance for dose response curves was calculated using two-way ANOVA test.

**FIgure S10**

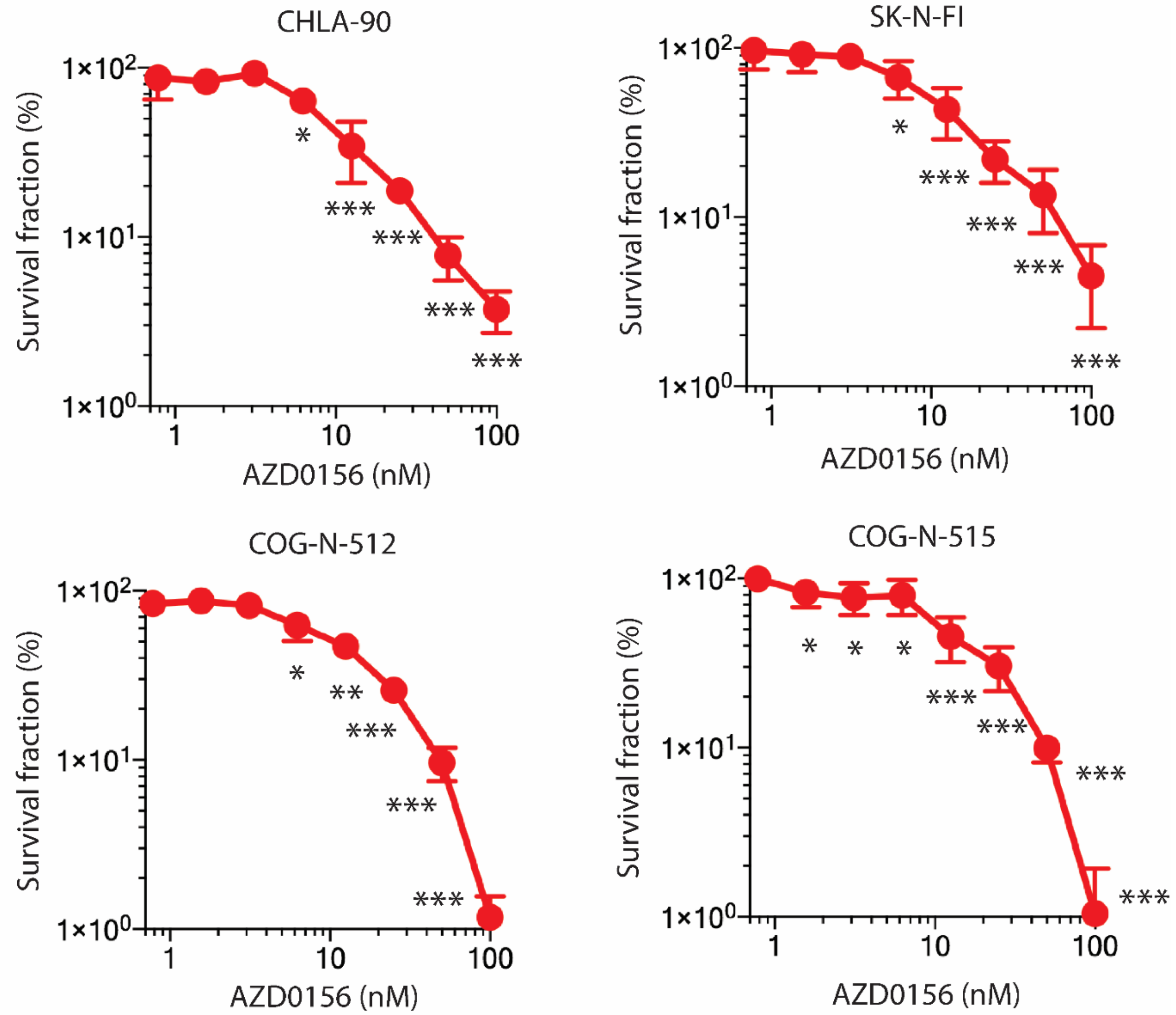

**Figure S10. AZD0156 enhanced TMZ+SN-38 activity at dose as low as 6.25nM in ALT neuroblastoma *in vitro*.** DIMSCAN cytotoxic assay curves in ALT neuroblastoma cell lines (CHLA-90, SK-N-FI, COG-N-512 and COG-N-515) treated with AZD0156 at variable dose in presence of constant TMZ (6.25 μM) + SN-38 (0.625 nM). Survival fraction of cells treated with TMZ (6.25 μM) + SN-38 (0.625 nM) alone is considered as 100%. Statistical significance for each point was calculated using Two-tailed t-test. ***: *P*<0.001, **: *P*<0.01, *: *P*<0.05.

**Figure S11**

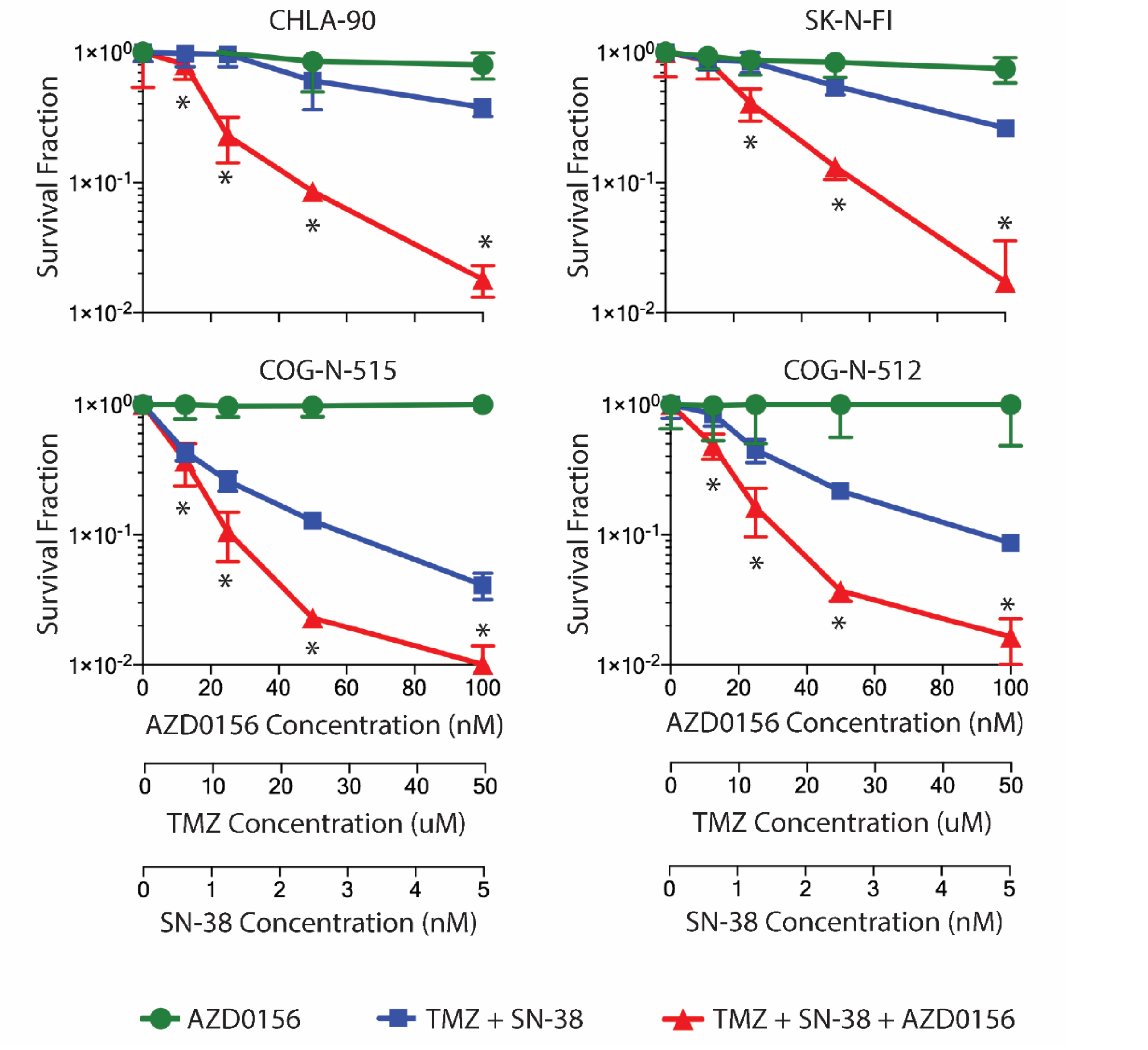

**Figure S11. AZD0156 synergized with TMZ+SN-38 in ALT neuroblastoma *in vitro.*** DIMSCAN cytotoxic assay curves in ALT neuroblastoma cell lines treated with AZD0156 + TMZ + SN-38 at equal molar fixed ratios in 4 ALT neuroblastoma cell lines. Combination Index (CI) for each point was calculated using COMPUSYN software. * = CI <1, indicating synergy.

**Figure S12**

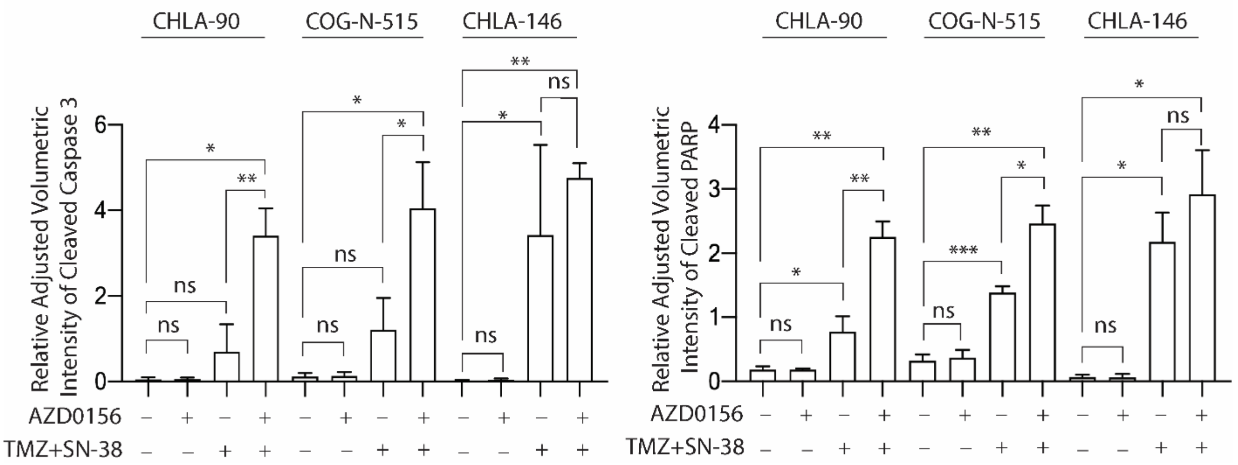

**Figure S12. The ATM inhibitor AZD0156 enhances induction of cleaved caspase 3 and cleaved PARP in ALT cell lines treated with TMZ+SN-38.** Normalized quantification of cleaved caspase 3 and cleaved PARP for the immunoblot in figure 5C and its replicates. The bars represent means with SDs from three experimental replicates. Statistical significance was calculated using two-tailed t-test. ***: *P*<0.001, **: *P*<0.01, *: *P*<0.05, ns: not significant.

**Figure S13**

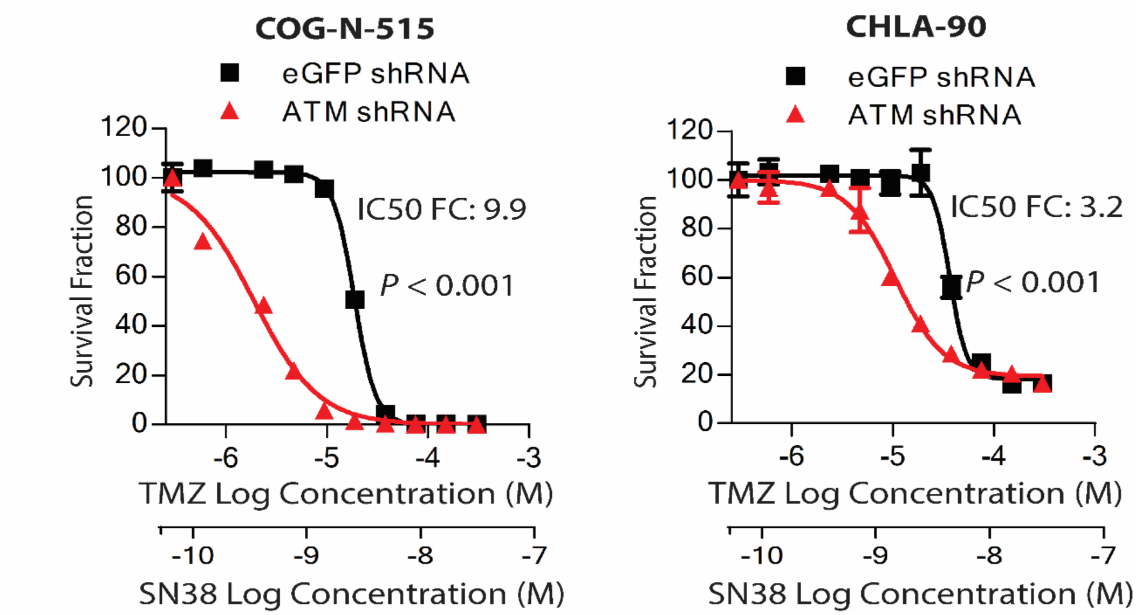

**Figure S13. ATM knockdown sensitized ALT neuroblastoma cell lines to TMZ+SN-38.** DIMSCAN cytotoxicity assay curves in response to TMZ+SN-38 in cells transduced with eGFP shRNA (black) or ATM shRNA (red). Statistical significance for dose response curves was calculated using two-way ANOVA test. IC50 FC: The half maximal inhibitory concentration fold-change.

**Figure S14**

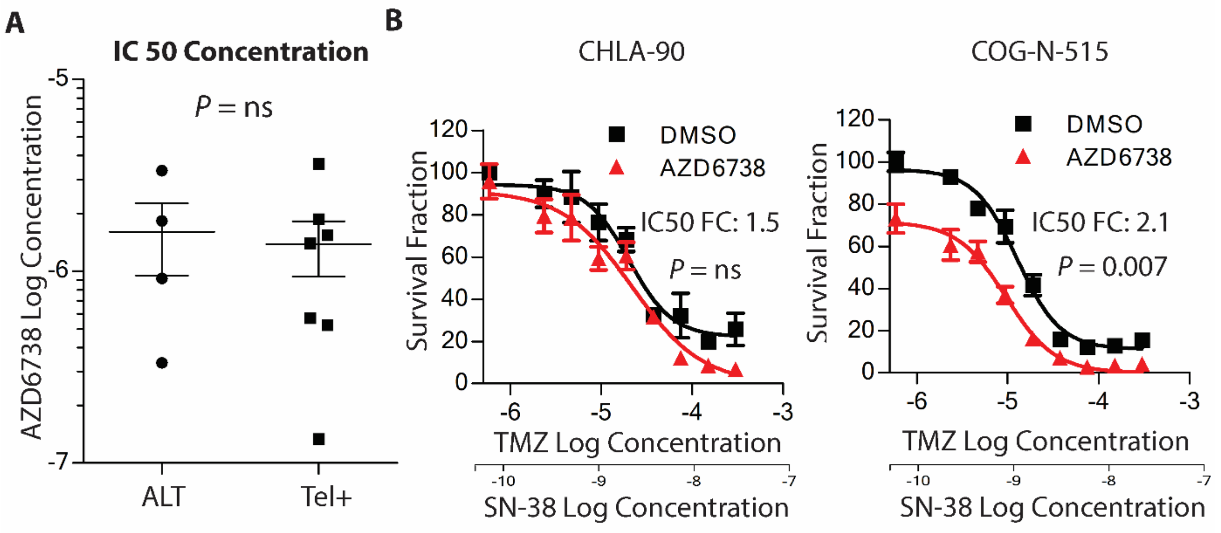

**Figure S14. ATR inhibitor AZD6738 did not enhance activity of TMZ+SN-38 in ALT neuroblastoma cell lines.** (A) AZD6738 IC50 for ALT (n=4) versus telomerase-positive (n=7) neuroblastoma cell lines. (B) DIMSCAN cytotoxicity curves for ALT neuroblastoma cell lines treated with TMZ+SN-38 +/-AZD6738. Statistical significance in A was calculated using Wilcoxon-rank sum test, and in B using two-way ANOVA test. ns: not significant. IC50 FC: The half maximal inhibitory concentration fold change.

**Figure S15**

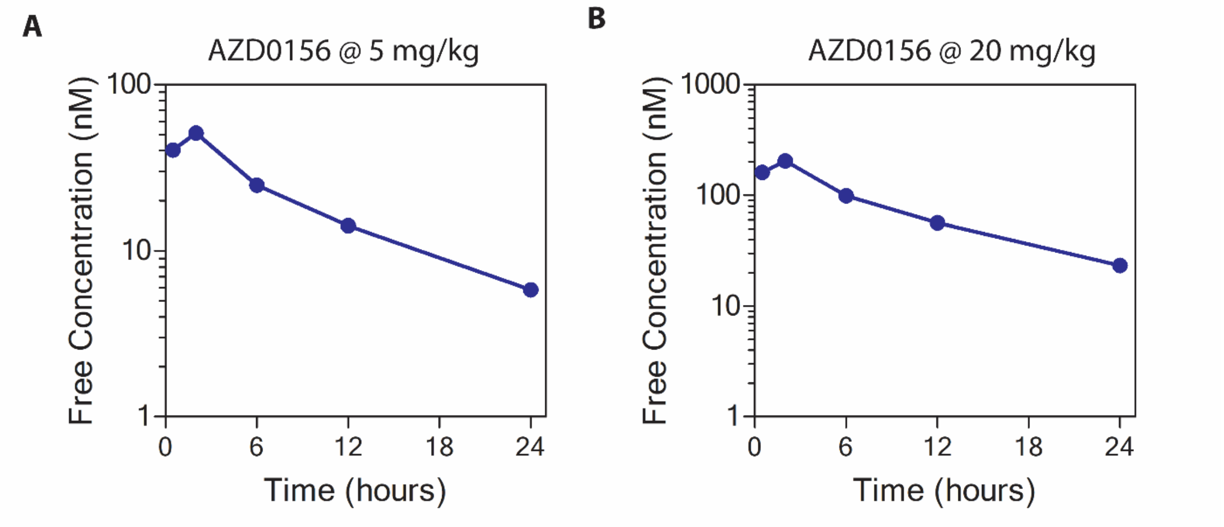

**Figure S15. Pharmacokinetic evaluation of unbound AZD0156 concentration *in vivo*.** Graph shows unbound AZD0156 plasma concentration in mice treated with (A) 5mg/kg or (B) 20mg/kg AZD0156 by oral gavage.

**Figure S16**

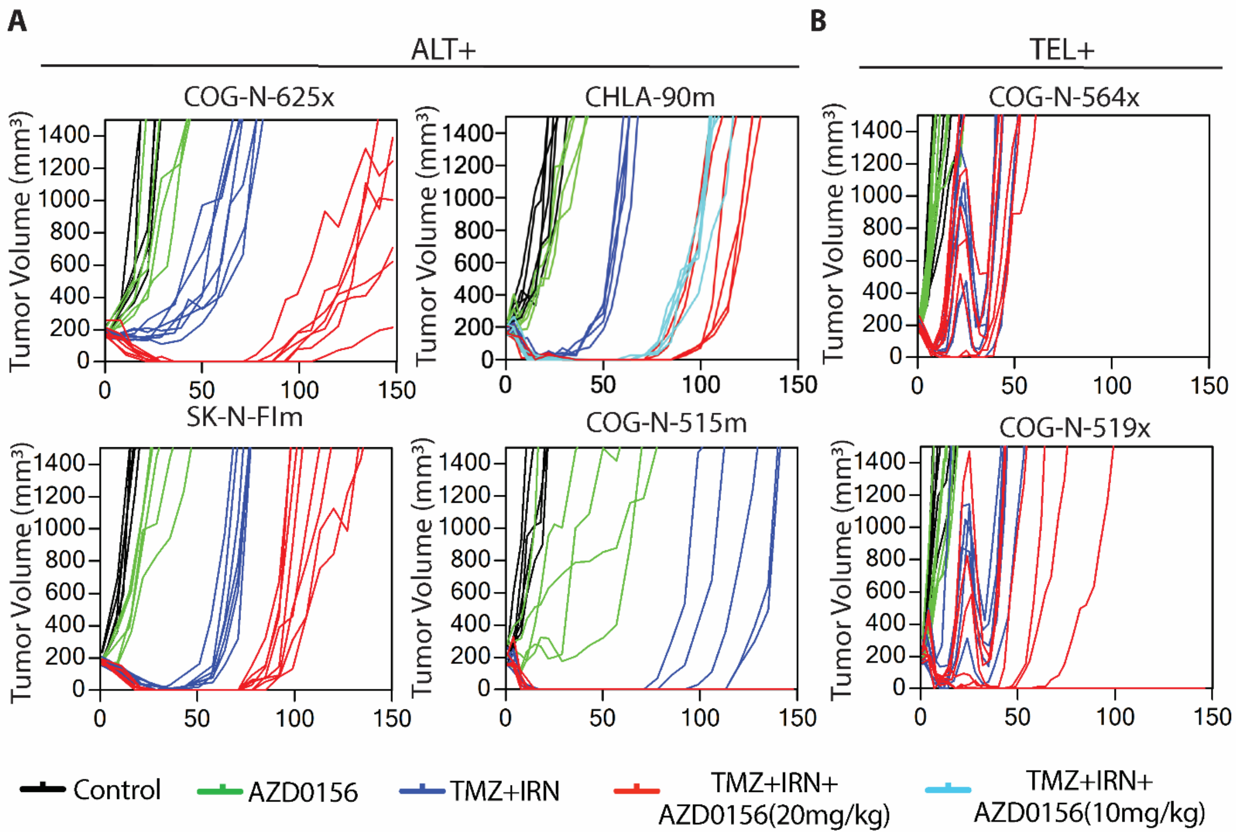

**Figure S16. Activity of TMZ+IRN +/- the ATM inhibitor AZD0156 combined with in ALT xenografts.** Tumor growth curves according to each treatment group for tumor-bearing (A) either ALT PDX (COG-N-625x) or ALT CDXs (CHLA-90m, SK-N-FIm or COG-N-515m) and (B) telomerase+ PDXs (COG-N-564x and COG-N-519x). Mice treated with temozolomide (TMZ) + irinotecan (IRN) were given 2 cycles of TMZ (25 mg/kg) + IRN (7.5 mg/kg) on Days 1-5 in a 21 day cycle; for the TMZ+IRN+AZD0156 group, AZD0156 (20 mg/kg and 10mg/kg (for CHLA-90 only)) was administered following TMZ+IRN treatment on Days 6-19;. Dosing was initiated when the initial tumor volume was between 150 to 250 $mm^3$.

**Figure S17**

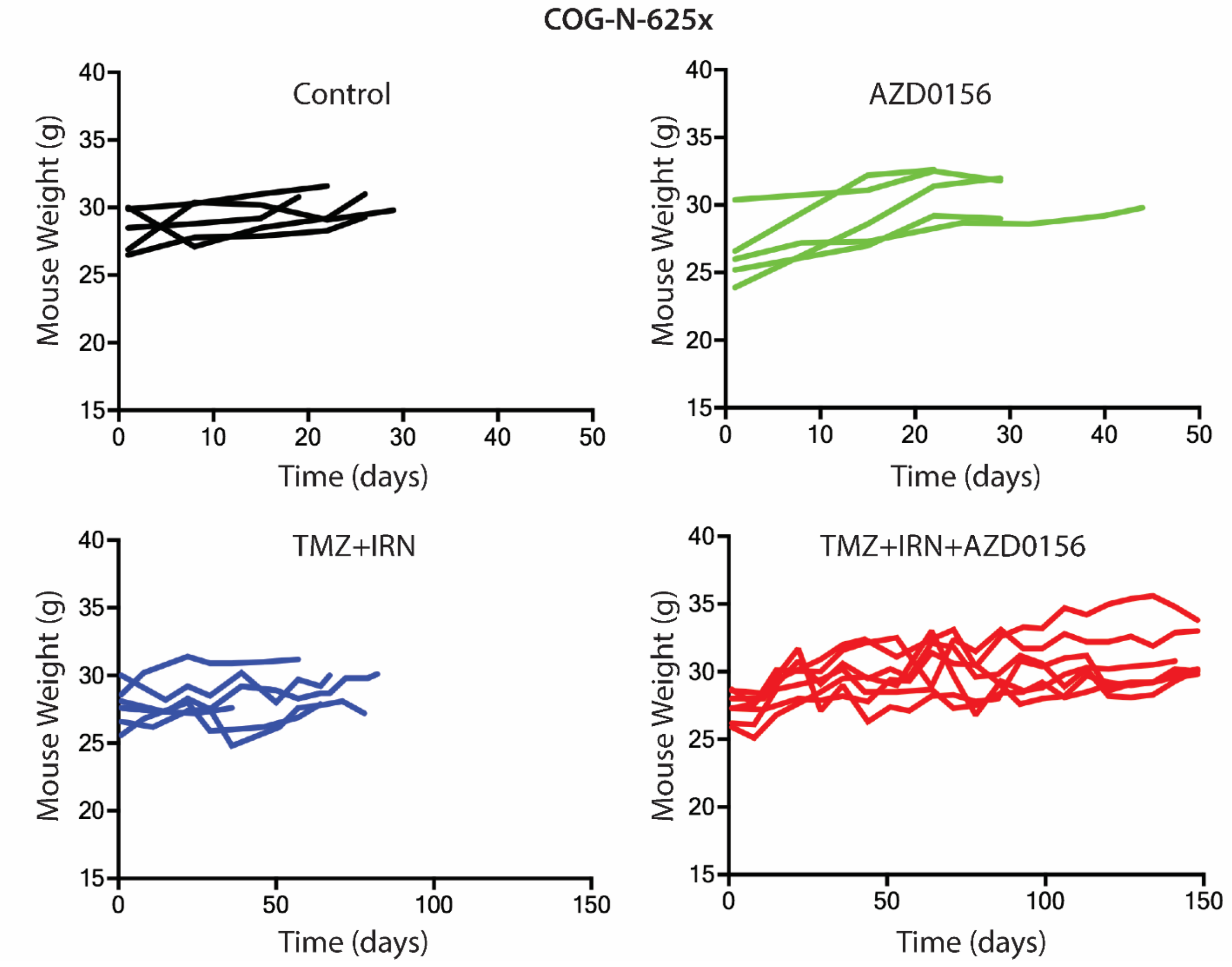

**Figure S17. AZD0156 sequenced following treatment with TMZ+IRN is well tolerated *in vivo*.** Body mass of COG-N-625x mice in each treatment group. None of the mice in any treatment group lost ≥ 20% of body mass at any point during treatment or follow-up.

#### Supplementary Methods

**Drugs and Chemicals.** Powdered temozolomide, melphalan (L-PAM), etoposide, topotecan hydrochloride hydrate, carboplatin, doxycycline hydrochloride, 7-Ethyl-10-hydroxycamptothecin (SN-38), and puromycin dihydrochloride were obtained from Sigma, St Louis, MO. 4-Hydroperoxycyclophosphamide was obtained from Chemos GmbH & Co, Altdorf, Germany. Irinotecan for intravenous injection *in vivo* was obtained from Sagent Pharmaceuticals, Schaumburg, IL. Captisol was obtained from CyDex Pharmaceuticals, Lenexa, KS. AZD0156 was provided by AstraZeneca, Cambridge, United Kingdom.

**Immunofluorescence and Telomere Fluorescence *in Situ* Hybridization (IF-FISH).** Neuroblastoma cells grown on sterile poly-L-lysine coated coverslips (Neuvitro, Vancouver, WA) were fixed in 4% paraformaldehyde at room temperature for 10 minutes and permeabilized for 10 minutes in 0.4% Triton X-100 buffer, then coverslips were incubated with blocking buffer (5% goat serum, in TBS) for 30 minutes, followed by incubation with primary antibody overnight. The antibodies used were 53BP1 (NB100-304; Novus; 1:500), MYC (9E10, Cell Signaling Technology, Danvers, MA; 1:100) and p-ATM and Anti-phospho-ATM (Ser1981) (05-740; Millipore, Burlington, MA; 1:400). The coverslips were washed with tris-buffered saline with 0.1% tween (TBST) for 3 times and then incubated with secondary antibody against rabbit or mouse that were conjugated with Alexa 555 or Alexa 488 (Molecular probes, Eugene, OR; Concentration: 1:1000). In case of IF staining alone, coverslips were washed with TBST for three times. The DNA was counterstained with DAPI in the second TBS-T wash. Coverslips were mounted in ProLong Gold antifade (Cell Signaling Technology, Danvers, MA) reagent and stored at -20°C until imaged. For IF-FISH, following secondary antibody incubation coverslips were washed thrice with TBS-T, fixed for 10 min in 4% paraformaldehyde at room temperature then dehydrated in 70%, 95% and 100% ethanol for 5 minutes each, and coverslips were allowed to dry. A FITC-OO-$[TTAGGG]_3$ PNA probe (PNA Bio, Newbury Park, CA) in buffer containing 70% deionized formamide, 10 mM Tris-HCl (pH 7.2) and 1 mg/ml blocking reagent (Roche, Basel, Switzerland) was added to coverslips and hybridized by denaturing on a heat block at 80°C for 5 minutes and then incubated in dark for 4 hours at room temperature. Coverslips were washed twice for 15 minutes with wash buffer containing 70% formamide, 10 mM Tris-HCl pH 7.2, and three times in TBS-T for 5 min each. The DNA was counterstained with DAPI in the second TBS-T wash. Coverslips were dehydrated in 70%, 95%, and 100% ethanol for 5 min each and allowed to air dry before mounting in ProLong gold antifade reagent. Images were obtained with a Nikon Ti-E microscope with A1 confocal system. Representative 2D images in the figures are maximum intensity projections of the Z stack. Images were analyzed using Fiji software. The average number of TIFs/cell across 3 ALT cell lines was 3.28 and 0.41 for telomerase positive cell lines, hence ≥ 3 TIFs was used a metric to compare ALT versus telomerase positive cell line. All image analysis was conducted blind.

**Immunoblotting.** Immunoblotting was performed as described previously (*53*). The following primary antibodies were used: TRF2 (#57130, Novus, Littleton, CO), anti-phospho-ATM (Ser1981) (#ab81292, Abcam, Cambridge, United Kingdom), ATM (#2873, Cell Signaling, Danvers, MA), Phospho-Chk2 (Thr68) (#2197, Cell Signaling), Chk2 (3440, Cell Signaling), Phospho-ATR (Ser428) (#2853, Cell Signaling), ATR (#2790, Cell Signaling), Phospho-Chk1 (Ser345) (#2344, Cell Signaling), Chk1 (2360, Cell Signaling), β-actin

(8457, Cell Signaling), Cleaved Caspase-3 (#9664, Cell Signaling), PARP (9542, Cell Signaling), and ATRX (sc-55584, Santa Cruz Biotechnology, Inc. Dallas, TX).

**C-circle Assay.** C-circle assay was performed as described previously (*29*).

**Comet assay.** Comet assay was performed on untreated cell lines using Comet Assay Kit (#ab238544, Abcam). Image analysis for comets was done blind using open comet plugin in Fiji software (*55*).

**Cell culture and Patient Derived Xenografts (PDXs).** All neuroblastoma patient-derived cell lines (PCLs) and xenografts (PDXs) were obtained from the ALSF/Children's Oncology Group (COG) Childhood Cancer Repository (www.CCcells.org). Cell lines and PDXs used in this study were established from blood, bone marrow, or tumor obtained at progressive disease following induction therapy (PD), progressive disease following myeloablative therapy (PD-BMT), or at post-mortem (PM) after death from progressive disease. COG- and CHLA- series cell lines were cultured in antibiotic-free Iscove's Modified Dulbecco's Medium supplemented with 20% fetal bovine serum (FBS) (GIBCO), 1 X ITS and 4 mM L-glutamine or with antibiotic and serum free neurobasal media supplemented with EGF, FGF, B-27 and N-2 supplements (cell lines where their name is followed by nb). SMS-, SK-N- and LA-N- series of cell lines were grown in RPMI-1640 supplemented with 10% FBS. Cell lines with h as suffix are established and grown in bone-marrow level oxygen tension (5% $O_2$). All human cell line identities were confirmed using the GenePrint 10 system (Promega, Madison, WI), compared to the database at www.CCcells.org and were routinely tested for lack of mycoplasma contamination using MycoAlert mycoplasma detection kit (Lonza, Basel, Switzerland). HEK293FT cells (ThemoFischer Scientific, Waltham, MA) were cultured in DMEM supplemented with 10% FBS, 4 mM L-glutamine, 1 mM MEM sodium pyruvate and 1% Pen-Strep (GIBCO, Waltham, MA).

Six- to eight-week-old athymic (nu/nu) mice (Jackson Laboratory, Bar Harbor, ME) were injected subcutaneously with 10 million neuroblastoma cells obtained directly from a subcutaneous tumor in nu/nu mice. The tumor volume and mouse weights were measured twice weekly. Tumor volumes were calculated as 0.5*height * width * length (*56*). The event-free survival (EFS) was defined as the time required for the tumor to grow from 150-250$mm^3$ to the endpoint (tumor volume ≥ 1500 $mm^3$), as described previously (*54*). Tissue obtained at the endpoint was verified for identity by STR profiling and human to mouse ratio was evaluated by PCR as described previously (*57*). Any PDX with less than 75% human cells were not used in this study.

**Stable Knockdown.** Lentiviral plasmid (pLKO.1) containing ATM (TRCN0000010299) shRNA, ATRX (TRCN0000013590) shRNA or EGFP shRNA (GE Healthcare Dharmacon, Inc, Lafayette, CO) were packaged using MISSION lentiviral packaging mix (Sigma, St Louis, MO). Desired cells were infected with viral media for 72 hrs. Successfully transduced cells were selected with 1.5 µg/ml of puromycin until stable clones were established.

***In vivo* Drug Testing.** The TTUHSC Institutional Animal Care and Use Committee (IACUC) approved all animal protocols. We used 4 cell line-derived xenografts (CDXs; CHLA-90m, SK-N-FIm, COG-N-515m,

FELIXm) and 5 patient-derived xenografts (PDXs; COG-N-625x, COG-N-519x, COG-N-564x, COG-N-452x and COG-N-623x) in this study. All the CDXs and PDXs were established from high-risk neuroblastoma patients at progressive disease after therapy or at postmortem at progressive disease after therapy. H&E staining of formalin-fixed paraffin-embedded sections and expression of tyrosine hydroxylase mRNA were assessed to verify neuroblastoma origin of new PDX models. PDXs and CDXs were verified to match the patient of origin by STR profiling and were passaged in mice (nu/nu) after establishment from the patient sample in NOD scid gamma mouse. Strained cells from a previous tumor were prepared for injection in RPMI-1640 and Matrigel (Corning). nu/nu mice were injected subcutaneously between the shoulder blades with 200 µl of cell preparation containing 10-20 million viable cells. Mice were randomized to treatment groups when progressively growing tumors reached 150-250 $mm^3$, determined as previously describe (*53*). CDX experiments were carried out similarly, as previously described (*58*).

*In Vivo* dosing of TMZ + IRN was designed to mimic clinical dosing (*59*). TMZ was prepared as a slurry in sterile water; 20 mg/ml of irinotecan hydrochloride was diluted 10x in 0.9% saline; AZD0156 was prepared in 10% DMSO and 30% captisol for oral gavage. In a 21-day cycle, on days 1 to 5 mice were treated with TMZ by oral gavage at 25 mg/kg, followed one hour later by IRN by tail vein injection at 7.5 mg/kg. Triple combination mice were additionally given AZD0156 by oral gavage at 20 mg/kg from days 6 to 19, sequencing of AZD0156 was designed to mimic clinical usage and previous *in vivo* testing (*48*).

*In vivo* responses were categorized based on National Cancer Institute Pediatric Preclinical Testing Program classification (*60*). For each mouse, complete response (CR) is defined as disappearance of measurable tumor mass (< 0.1$cm^3$) at any time following treatment during the study period. A CR is considered as maintained (MCR) if the tumor volume was undetectable (<0.1 $cm^3$) at the end of study period. Partial response (PR) is defined as tumor volume that reduces by > 50% of initial volume for at least one time point during the study but with a measurable tumor mass (≥ 0.1$cm^3$). Progressive disease (PD) was defined as < 50% reduction in initial tumor volume at any point during the study and > 25% increase in tumor volume by the end of the study. Stable disease was defined as < 50% reduction in initial tumor volume until the end of the study and ≤ 25% increase in tumor volume by the end of the study. An event-free survival of tumor/control (EFS T/C) was calculated by the ratio of the median time to an event in the treatment group divided by median time to an event in the control group. For EFS T/C measure, a treatment is considered as highly active if EFS T/C is > 2, statistically significant (Log-rank test: P ≤ 0.05) difference in EFS compared to control, and there is a net reduction in median tumor volume by the end of treatment. Treatment groups with EFS T/C > 2 with statistically significant (Log-rank test: P ≤ 0.05) difference in EFS compared to control, but no net reduction in median tumor volume is considered to have intermediate activity. For EFS T/C < 2, the agents are considered to have low activity. If the treatment group does not have a median EFS, then EFS T/C is defined as greater than the ratio of last day of follow up for the treatment group divided by median time to an event in the control group.

**Immunohistochemistry and TUNEL on FFPE sections.** Immunohistochemical staining was performed on 4 µm sections from formalin-fixed paraffin-embedded (FFPE) tissue blocks. Slides were first deparaffinized with serial xylene treatment for 5 minutes each, followed by hydration in a graded ethanol series. The hydrated slide sections were heated at 100-110°C for antigen retrieval for 30 minutes with EDTA buffer (pH 9.0) for phospho-ATM and citrate buffer (pH 6.0) for anti-Ki67 immunohistochemistry and TUNEL staining in a pressure cooker. For immunohistochemistry, after slides were cooled down to room temperature for 10 minutes, sections were blocked with Dual Endogenous Enzyme-Blocking Agent (Dako) for 10 minutes. Sections were blocked for 30 minutes with 10% serum in TBST, followed by incubation with either anti-phospho-ATM (Ser1981) (1:50, SAB4300100, Sigma) or anti-Ki-67 (1:500, PLA0228, Sigma) for 1 hour, followed by anti-rabbit secondary antibody (Leica Microsystems) for 30 minutes at room temperature, and detected with 3, 3'-diaminobenzidine (Sigma-Aldrich). All wash steps were performed with PBS containing 0.1% tween-20 three times for 5 minutes. Finally, sections were counterstained with hematoxylin, dehydrated with a graded ethanol and xylene series, and were mounted for microscopy. For TUNEL staining on FFPE sections, following blocking with 10% goat serum, tissue sections were incubated with tunnel staining mixture from the *In Situ* Cell Death Detection Kit (11684795910, Roche) for 1h at 37°C. Sections were washed 3 times with TBST and then mounted with ProLong Gold antifade reagent for microscopy. Images were obtained on Olympus BX51 fluorescent microscope using 20X objective.

**Pharmacokinetic evaluation of AZD0156.** The pharmacokinetics (PK) of AZD0156 was investigated by a multiple sampling PK study, consisting of three animals per time points. Swiss athymic nu/nu male mice were dosed at 5 mg/kg dose with PK samples taken at 0.5, 2, 6, 12 and 24h. The oral dosing formulation was AZD0156 dissolved in DMSO/water. The plasma concentrations of AZD0156 were determined using a protein precipitation procedure followed by liquid chromatography with a tandem mass spectrometric detection method. The lower limit of quantification was 0.0014 μM. PK for 20 mg/kg was scaled up by dose proportionality. The measured mouse plasma concentrations were fitted to obtain PK parameters using a non-compartmental method utilizing WinNonlin software.

**Table S1** **Patient-derived neuroblastoma cell lines and patient-derived xenografts**

| Cell Line | Phase of Therapy | Sample Type | Disease Stage | MYCN | TH mRNA | p53 Status | ALK Status | Doubling Time (hours) | TERT Expression | ALT Status | RNA-seq | Drug Response Data for Single Agents | Drug Response Data for Combination |
|---|---|---|---|---|---|---|---|---|---|---|---|---|---|
| CHLA-108 | PD | BM | 4 | A | + | NA | NA | NA | + | - | - | + | + |
| CHLA-11 | PD-BMT, PM | Blood | 4 | A | + | NF | WT | 53 | + | - | - | + | - |
| CHLA-119 | PD | BM | 4 | A | + | NF | WT | NA | + | - | + | + | + |
| CHLA-12 | PD-BMT, PM | BM | 4 | A | + | NF | WT | 53 | + | - | - | + | - |
| CHLA-125 | PD | BM | 4 | NA | + | NA | NA | NA | + | - | - | + | + |
| CHLA-136 | PD-BMT | Blood | 4 | A | + | F | WT | 44 | + | - | - | + | + |
| CHLA-140 | PD | BM | 4 | N | + | NA | WT | NA | + | - | - | + | - |
| CHLA-146 | PD | BM | 4 | NA | + | F | NA | NA | + | - | - | + | + |
| CHLA-150 | PD | BM | 4 | N | + | NA | R1275Q | NA | + | - | - | + | - |
| CHLA-171 | PD-PM | Blood | 4 | N | + | NF | WT | 89 | + | - | + | + | + |
| CHLA-172 | PD-BMT | BM | 4 | N | + | NF | WT | NA | + | - | - | + | + |
| CHLA-21 | PD | BM | 4 | A | + | NA | NA | NA | + | - | - | + | - |
| CHLA-22 | PD | BM | 4 | NA | + | NA | NA | NA | + | - | - | + | - |
| CHLA-225 | PD-PM | Fluid | 4 | A | + | NF | WT | 62 | + | - | - | + | + |
| CHLA-247 | PD | BM | 4 | A | + | NA | WT | NA | + | - | + | + | - |
| CHLA-257 | PD | BM | 4 | NA | + | F | NA | NA | + | - | - | + | + |
| CHLA-42 | Dx | BM | 4 | N | + | F | R1275Q | NA | + | - | - | + | - |
| CHLA-52 | PD-PM | BM | 4 | A | + | NF | F1174L | 88 | + | - | - | + | - |
| CHLA-53 | PD-PM | Blood | 4 | A | + | NF | F1174L | 92 | + | - | - | + | - |
| CHLA-60 | PD-PM | BM | 4 | N | + | F | WT | 96 | + | - | - | + | - |
| CHLA-61 | PD-PM | Blood | 4 | N | + | F | R1275Q | 102 | + | - | - | + | + |
| CHLA-70 | PD | BM | 4 | NA | + | F | NA | NA | + | - | - | + | + |
| CHLA-79 | PD-BMT | Tumor | 4 | N | + | F | WT | NA | + | - | - | + | + |
| CHLA-90 | PD-BMT | BM | 4 | N | + | NF | F1245V | NA | - | + | + | + | + |
| COG-N-261 | Dx | Tumor | 4 | A | + | NA | WT | 124 | + | - | - | + | - |
| COG-N-263 | Dx | Tumor | 1 | N | + | NA | WT | 119 | + | - | - | + | - |
| COG-N-269 | PD | Tumor | 2B | N | + | NA | WT | 91 | + | - | - | + | + |
| COG-N-278 | Dx | Tumor | 1 | A | + | NA | WT | 88 | + | - | - | + | - |
| COG-N-283 | Dx | Tumor | 4 | A | + | NA | NA | NA | + | - | - | + | - |
| COG-N-289 | PD | Tumor | 4 | A | + | NA | WT | 38 | + | - | - | + | - |
| COG-N-293 | Dx | Tumor | 4 | A | + | NA | WT | 88 | + | - | - | + | - |
| COG-N-295 | Dx | Tumor | 3 | N | + | NA | F1174L | 58 | + | - | - | + | + |
| COG-N-297 | Dx | Tumor | 4 | A | + | NA | WT | 28 | + | - | - | + | - |
| COG-N-299 | Dx | Tumor | 4 | A | + | NA | WT | 204 | + | - | - | + | - |
| COG-N-303 | PD | Tumor | 4s | A | + | NA | WT | 47 | + | - | - | + | + |
| COG-N-305 | Dx | Tumor | 4 | A | + | NA | WT | 187 | + | - | - | + | - |
| COG-N-312 | Dx | BM | 4 | A | + | F | NA | 35 | + | - | - | + | + |
| COG-N-316 | Dx | BM | 4 | A | + | NA | NA | NA | + | - | - | + | - |
| COG-N-318 | PD | Tumor | 4s | A | + | NA | WT | 129 | + | - | - | + | - |
| COG-N-321 | PD | BM | 3 | A | + | NF | NA | NA | + | - | - | + | - |
| COG-N-322 | Dx | Tumor | 4 | A | + | NA | WT | 106 | + | - | - | + | - |
| COG-N-323 | PD | Tumor | 4 | N | + | NA | WT | 25 | - | - | - | + | - |
| COG-N-325 | PD | Tumor | 4 | A | + | NA | WT | 55 | + | - | - | + | - |
| COG-N-327 | PD | BM | 4 | A | + | NA | WT | 204 | - | - | - | + | - |
| COG-N-328h | Dx | Tumor | 4 | A | + | NA | WT | 116 | + | - | - | + | - |
| COG-N-331 | Dx | BM | 4 | A | + | NA | WT | 55 | + | - | - | + | - |
| COG-N-333 | Dx | BM | 4 | A | + | NA | F1174L | 73 | + | - | - | + | - |
| COG-N-334 | PD | BM | 4 | A | + | NA | WT | 48 | + | - | - | + | - |
| COG-N-346h | PD | Tumor | 4 | N | + | NA | WT | 83 | - | - | - | + | - |
| COG-N-347 | Dx | BM | 4 | A | + | NA | WT | 126 | - | - | - | + | - |
| COG-N-349 | Dx | BM | 4 | A | + | NA | WT | 192 | + | - | - | + | - |
| COG-N-353 | Dx | BM | 4 | A | + | NA | WT | 144 | + | - | - | + | - |
| COG-N-354h | Dx | BM | 4 | A | + | NA | NA | 112 | + | - | - | + | - |
| COG-N-367 | Dx | BM | 4 | A | + | NA | R1275Q | 49 | + | - | - | + | - |
| COG-N-372 | PD | BM | 3 | N | + | NA | WT | 96 | - | - | - | + | + |
| COG-N-373 | Dx | BM | 4 | A | + | NA | WT | 126 | + | - | - | + | - |
| COG-N-387 | PD | BM | 4 | A | + | NA | WT | NA | - | - | - | + | - |
| COG-N-396h | PD | Tumor | 4 | NA | + | NA | NA | NA | + | - | - | + | - |
| COG-N-399 | PD-PM | Blood | 4 | A | + | F | WT | 40 | + | - | - | + | - |
| COG-N-415 | PD | Blood | 4 | A | + | NA | F1174L | 50 | + | - | + | + | + |
| COG-N-421h | PD-PM | Blood | 4 | A | + | NF | WT | 65 | + | - | + | + | - |
| COG-N-440h | PD-PM | Blood | 4 | A | + | F | WT | 65 | + | - | + | + | + |
| COG-N-442h | Dx | BM | 4 | N | + | NA | WT | 98 | - | - | - | + | + |
| COG-N-443h | PD | BM | 4 | N | + | NA | F1245C | 51 | + | - | - | + | + |
| COG-N-445 | Dx | BM | 4 | A | + | NA | WT | 217 | - | - | - | + | - |
| COG-N-452 | PD-PM | Blood | 4 | A | + | F | F1174L | 74 | + | - | + | + | + |
| COG-N-453 | PD-PM | Tumor | 4 | A | + | F | WT | 71 | + | - | + | + | + |
| COG-N-462h | Dx | BM | 4 | A | + | NA | WT | 120 | + | - | - | + | - |
| COG-N-463h2 | Dx | BM | 4 | A | + | NA | WT | NA | + | - | - | + | - |
| COG-N-469h | PD | BM | 4 | N | + | NA | F1174L | 53 | + | - | - | + | - |
| COG-N-470 | PD-PM | Blood | 4 | N | + | F | F1174L | 82 | + | - | - | + | + |
| COG-N-471 | PD-PM | Blood | 4 | A | + | NF | WT | 136 | + | - | - | + | + |
| COG-N-473 | PD | BM | 4 | A | + | NA | WT | NA | + | - | - | + | - |
| COG-N-474 | Dx | BM | 4 | A | + | NA | F1174L | 187 | + | - | - | + | - |
| COG-N-475h | PD | Tumor | 4 | A | + | NA | R1275Q | 96 | + | - | - | + | - |
| COG-N-476 | PD-BMT | Tumor | 4 | A | + | NA | WT | 50 | + | - | - | + | - |

| | | | | | | | | | | | | | |
|---|---|---|---|---|---|---|---|---|---|---|---|---|---|
| COG-N-478 | PD | BM | 4 | N | + | *NA* | WT | *NA* | + | - | - | + | - |
| COG-N-480hnb | Dx | BM | 4 | A | + | *NA* | *NA* | *NA* | + | - | - | + | - |
| COG-N-485h2 | Dx | BM | 4 | A | + | *NA* | *NA* | *NA* | + | - | - | + | - |
| COG-N-490h | Dx | BM | 4 | A | + | *NA* | WT | *NA* | + | - | - | + | - |
| COG-N-496 | Dx | BM | 4 | A | + | NF | WT | *NA* | + | - | - | + | - |
| COG-N-500 | Dx | BM | 4 | N | + | *NA* | *NA* | *NA* | + | - | - | + | - |
| COG-N-504 | Dx | BM | 4 | N | + | *NA* | *NA* | 315 | + | - | - | + | - |
| COG-N-505 | PD-PM | Blood | 4S | A | + | F | WT | 79 | + | - | + | + | - |
| COG-N-511hnb | PD | BM | 4 | N | + | *NA* | *NA* | *NA* | + | - | - | + | - |
| COG-N-512* | PD | BM | 4 | A | + | NF | *NA* | *NA* | - | + | + | + | + |
| COG-N-514 | PD | BM | 4 | A | + | *NA* | *NA* | *NA* | + | - | - | + | - |
| COG-N-515* | PD | BM | 4 | A | + | NF | *NA* | *NA* | - | + | + | + | + |
| COG-N-519 | PD-PM | Blood | 4 | A | + | NF | WT | 38 | + | - | + | + | + |
| COG-N-534 | PD-PM | Blood | 4 | N | + | NF | F1245C | 144 | + | - | - | + | - |
| COG-N-549h | PD | Tumor | 4 | N | + | *NA* | | 14 | + | - | - | + | - |
| COG-N-564 | PD-PM | Blood | 4 | A | + | NF | *NA* | *NA* | + | - | + | + | + |
| COG-N-579h | PD | BM | 4 | N | + | *NA* | *NA* | 120 | + | - | + | + | + |
| COG-N-615 | PD-PM | Blood | 4 | N | + | *NA* | *NA* | 117 | + | - | + | + | + |
| COG-N-618 | PD | BM | 4 | N | + | *NA* | *NA* | 57 | + | - | + | + | + |
| COG-N-619h | PD | BM | 4 | N | + | *NA* | *NA* | 94 | + | - | + | + | + |
| COG-N-708h | Dx | Tumor | 4 | N | + | *NA* | *NA* | 103 | + | - | + | + | + |
| COG-N-709 | Dx | Tumor | 4 | N | + | *NA* | *NA* | *NA* | + | - | + | + | + |
| Felix | PD-PM | Blood | 4 | N | + | F | F1245V | 62 | + | - | + | + | + |
| FU-NB-2006 | PD-PM | Blood | 4 | N | + | *NA* | WT | 48 | + | - | - | + | - |
| SK-N-BE(1)º | Dx | *NA* | 4 | A | + | F | WT | 96 | + | - | - | + | + |
| SK-N-BE(2)º | PD | *NA* | 4 | A | + | NF | WT | 27 | + | - | + | + | + |
| SK-N-FI | PD | BM | 4 | N | + | NF | WT | *NA* | - | + | + | + | + |
| SMS-SAN | Dx | BM | 4 | A | + | F | F1174L | *NA* | + | - | - | + | - |

| | |
|---|---|
| º | Established from same patient at different stage of Disease |
| * | Established from same patient at same time, but from different location of the body |

| | |
|---|---|
| BM: | Bone Marrow |
| PD: | Progressive Disease |
| DX: | Diagnosis |
| PD-PM: | Progressive Disease at Post Mortem |
| PD-BMT | Progressive Disease at Bone Marrow Transplantation |
| TH: | Tyrosine Hydroxylase |
| F | Functional |
| NF | Non-functional |
| N: | non-Amplified |
| A: | Amplified |
| *NA* : | Not Available |

| PDX | Phase of Therapy | Stage | MYCN Status | TH Expressio | ALK | Sample Type | ALT Status |
|---|---|---|---|---|---|---|---|
| COG-N-625x | PD | 4 | N | + | WT | Tumor | + |
| COG-N-452x | PD-PM | 4 | A | + | F1174L | Blood | - |
| COG-N-623x | PD | 4 | A | + | WT | BM | - |
| COG-N-421x | PD-PM | 4 | A | + | WT | Blood | - |
| COG-N-470x | PD-PM | 4 | N | + | F1174L | Blood | - |
| COG-N-561x | PD-PM | 4 | A | + | F1245I | Blood | - |
| COG-N-519x | PD-PM | 4 | A | + | WT | Blood | - |
| COG-N-564x | PD-PM | 4 | A | + | WT | Blood | - |

| CDX | Phase of Therapy | Stage | MYCN Status | TH Expressio | ALK | Sample Type | ALT Status |
|---|---|---|---|---|---|---|---|
| CHLA-90m | PD-BMT | 4 | N | + | F1245V | BM | + |
| SK-N-FIm | PD | 4 | N | + | WT | BM | + |
| COG-N-515m | PD | 4 | A | + | WT | BM | + |
| CHLA-79m | PD-BMT | 4 | N | + | WT | Tumor | - |
| CHLA-119m | PD | 4 | A | + | WT | BM | - |
| Felix-m | PD-PM | 4 | N | + | F1254C | Blood | - |

| | |
|---|---|
| PDX: | Patient Dervied Xenografts |
| CDX: | Cell-line Dervied Xenografts |
| BM: | Bone Marrow |
| PD: | Progressive Disease |
| PD-PM: | Progressive Disease at Post Mortem |
| PD-BMT: | Progressive Disease at Bone Marrow Transplantation |
| TH: | Tyrosine Hydroxylase |
| N: | non-Amplified |
| A: | Amplified |
| *NA* : | Not Available |

#### Table S2 Somatic mutations identified for COG-N-515 and COG-N-512

| Hugo Symbol | Entrez Gene | Chromosome | Start position | End position | Variant Classif | Variant | Reference | Tumor Seq Allele1 | Tumor Seq Allele2 | Tumor Sample Barcode1 | Genome Change | Annotation Transcript | Transcript Strand | Transcript Exon | Transcript Position | cDNA Change | Codon Change | Protein Change |
|---|---|---|---|---|---|---|---|---|---|---|---|---|---|---|---|---|---|---|
| CCDC19 | 0 | 1 | 159854305 | 159854305 | Missense Mut | SNP | G | G | A | COG-N-512-D | g.chr1:159854305G>A | ENST00000368099.4 | - | 7 | 882 | c.818C>T | c.(817-819)tCg>tTg | p.S273L |
| CCDC19 | 0 | 1 | 159854305 | 159854305 | Missense Mut | SNP | G | G | A | COG-N-515-D | g.chr1:159854305G>A | ENST00000368099.4 | - | 7 | 882 | c.818C>T | c.(817-819)tCg>tTg | p.S273L |
| FAM194A | 0 | 3 | 150398570 | 150398570 | Missense Mut | SNP | C | C | A | COG-N-512-D | g.chr3:150398570C>A | ENST00000295910.6 | - | 8 | 1082 | c.1030G>T | c.(1030-1032)Gcc>Tcc | p.A344S |
| FAM194A | 0 | 3 | 150398570 | 150398570 | Missense Mut | SNP | C | C | A | COG-N-515-D | g.chr3:150398570C>A | ENST00000295910.6 | - | 8 | 1082 | c.1030G>T | c.(1030-1032)Gcc>Tcc | p.A344S |
| ABCA3 | 21 | 16 | 2347515 | 2347515 | Missense Mut | SNP | G | G | A | COG-N-515-D | g.chr16:2347515G>A | ENST00000382381.3 | - | 16 | 2615 | c.1904C>T | c.(1903-1905)tCg>tTg | p.S635L |
| ADORA3 | 140 | 1 | 112031415 | 112031415 | Missense Mut | SNP | G | G | A | COG-N-512-D | g.chr1:112031415G>A | ENST00000369716.4 | - | 3 | 822 | c.689C>T | c.(688-690)aCg>aTg | p.T230M |
| ADORA3 | 140 | 1 | 112031415 | 112031415 | Missense Mut | SNP | G | G | A | COG-N-515-D | g.chr1:112031415G>A | ENST00000369716.4 | - | 3 | 822 | c.689C>T | c.(688-690)aCg>aTg | p.T230M |
| AQP3 | 360 | 9 | 33443355 | 33443355 | Missense Mut | SNP | A | A | G | COG-N-512-D | g.chr9:33443355A>G | ENST00000297991.4 | - | 3 | 417 | c.337T>C | c.(337-339)Ttc>Ctc | p.F113L |
| AQP3 | 360 | 9 | 33443355 | 33443355 | Missense Mut | SNP | A | A | G | COG-N-515-D | g.chr9:33443355A>G | ENST00000297991.4 | - | 3 | 417 | c.337T>C | c.(337-339)Ttc>Ctc | p.F113L |
| ASPH | 444 | 8 | 62479787 | 62479787 | Missense Mut | SNP | C | C | A | COG-N-515-D | g.chr8:62479787C>A | ENST00000379454.4 | - | 17 | 1427 | c.1240G>T | c.(1240-1242)Gtc>Ttc | p.V414F |
| BRAF | 673 | 7 | 140508707 | 140508707 | Missense Mut | SNP | T | T | A | COG-N-512-D | g.chr7:140508707T>A | ENST00000288602.6 | - | 4 | 653 | c.593A>T | c.(592-594)tAc>tTc | p.Y198F |
| BRAF | 673 | 7 | 140508707 | 140508707 | Missense Mut | SNP | T | T | A | COG-N-515-D | g.chr7:140508707T>A | ENST00000288602.6 | - | 4 | 653 | c.593A>T | c.(592-594)tAc>tTc | p.Y198F |
| LDLRAD4 | 753 | 18 | 13645129 | 13645129 | Missense Mut | SNP | A | A | G | COG-N-512-D | g.chr18:13645129A>G | ENST00000359446.5 | + | 6 | 862 | c.394A>G | c.(394-396)Atg>Gtg | p.M132V |
| CACNA1B | 774 | 9 | 140773504 | 140773504 | Splice Site | DEL | A | A | - | COG-N-515-D | g.chr9:140773504delA | ENST00000371372.1 | + | 2 | 429 | | c.e2-1 | |
| CCND3 | 896 | 6 | 41903784 | 41903784 | Missense Mut | SNP | G | G | A | COG-N-512-D | g.chr6:41903784G>A | ENST00000372991.4 | - | 5 | 971 | c.773C>T | c.(772-774)gCc>gTc | p.A258V |
| CCND3 | 896 | 6 | 41903784 | 41903784 | Missense Mut | SNP | G | G | A | COG-N-515-D | g.chr6:41903784G>A | ENST00000372991.4 | - | 5 | 971 | c.773C>T | c.(772-774)gCc>gTc | p.A258V |
| CEL | 1056 | 9 | 135945951 | 135945951 | Missense Mut | SNP | G | G | T | COG-N-512-D | g.chr9:135945951G>T | ENST00000372080.4 | + | 10 | 1415 | c.1399G>T | c.(1399-1401)Ggg>Tgg | p.G467W |
| CEL | 1056 | 9 | 135945951 | 135945951 | Missense Mut | SNP | G | G | T | COG-N-515-D | g.chr9:135945951G>T | ENST00000372080.4 | + | 10 | 1415 | c.1399G>T | c.(1399-1401)Ggg>Tgg | p.G467W |
| CENPC | 1060 | 4 | 68357993 | 68357993 | Splice Site | SNP | T | T | A | COG-N-515-D | g.chr4:68357993T>A | ENST00000273853.6 | - | 16 | 2670 | c.2420A>T | c.(2419-2421)aAg>aTg | p.K807M |
| CENPE | 1062 | 4 | 104115409 | 104115409 | Missense Mut | SNP | C | C | T | COG-N-512-D | g.chr4:104115409C>T | ENST00000380026.3 | - | 8 | 758 | c.670G>A | c.(670-672)Gga>Aga | p.G224R |
| CENPE | 1062 | 4 | 104115409 | 104115409 | Missense Mut | SNP | C | C | T | COG-N-515-D | g.chr4:104115409C>T | ENST00000380026.3 | - | 8 | 758 | c.670G>A | c.(670-672)Gga>Aga | p.G224R |
| CHD2 | 1106 | 15 | 93485170 | 93485170 | Missense Mut | SNP | C | C | G | COG-N-512-D | g.chr15:93485170C>G | ENST00000394196.4 | + | 8 | 1879 | c.811C>G | c.(811-813)Ctg>Gtg | p.L271V |
| CHD2 | 1106 | 15 | 93485170 | 93485170 | Missense Mut | SNP | C | C | G | COG-N-515-D | g.chr15:93485170C>G | ENST00000394196.4 | + | 8 | 1879 | c.811C>G | c.(811-813)Ctg>Gtg | p.L271V |
| ERCC8 | 1161 | 5 | 60183337 | 60183337 | Missense Mut | SNP | C | C | A | COG-N-512-D | g.chr5:60183337C>A | ENST00000265038.5 | - | 11 | 1094 | c.1052G>T | c.(1051-1053)aGt>aTt | p.S351I |
| ERCC8 | 1161 | 5 | 60183337 | 60183337 | Missense Mut | SNP | C | C | A | COG-N-515-D | g.chr5:60183337C>A | ENST00000265038.5 | - | 11 | 1094 | c.1052G>T | c.(1051-1053)aGt>aTt | p.S351I |
| COL11A1 | 1301 | 1 | 103544238 | 103544238 | Missense Mut | SNP | C | C | A | COG-N-512-D | g.chr1:103544238C>A | ENST00000370096.3 | - | 3 | 776 | c.464G>T | c.(463-465)aGa>aTa | p.R155I |
| COL11A1 | 1301 | 1 | 103544238 | 103544238 | Missense Mut | SNP | C | C | A | COG-N-515-D | g.chr1:103544238C>A | ENST00000370096.3 | - | 3 | 776 | c.464G>T | c.(463-465)aGa>aTa | p.R155I |
| CSF1R | 1436 | 5 | 149437096 | 149437096 | Missense Mut | SNP | G | G | T | COG-N-512-D | g.chr5:149437096G>T | ENST00000286301.3 | - | 16 | 2483 | c.2192C>A | c.(2191-2193)aCt>aAt | p.T731N |
| CSF1R | 1436 | 5 | 149437096 | 149437096 | Missense Mut | SNP | G | G | T | COG-N-515-D | g.chr5:149437096G>T | ENST00000286301.3 | - | 16 | 2483 | c.2192C>A | c.(2191-2193)aCt>aAt | p.T731N |
| CYP1B1 | 1545 | 2 | 38301951 | 38301951 | Missense Mut | SNP | C | C | G | COG-N-515-D | g.chr2:38301951C>G | ENST00000260630.3 | - | 2 | 982 | c.581G>C | c.(580-582)aGg>aCg | p.R194T |
| CYP3A4 | 1576 | 7 | 99358472 | 99358472 | Missense Mut | SNP | G | G | T | COG-N-512-D | g.chr7:99358472G>T | ENST00000336411.2 | - | 12 | 1569 | c.1386C>A | c.(1384-1386)aaC>aaA | p.N462K |
| CYP3A4 | 1576 | 7 | 99358472 | 99358472 | Missense Mut | SNP | G | G | T | COG-N-515-D | g.chr7:99358472G>T | ENST00000336411.2 | - | 12 | 1569 | c.1386C>A | c.(1384-1386)aaC>aaA | p.N462K |
| EGF | 1950 | 4 | 110932470 | 110932470 | Missense Mut | SNP | G | G | C | COG-N-512-D | g.chr4:110932470G>C | ENST00000509793.1 | + | 23 | 3809 | c.3357G>C | c.(3355-3357)atG>atC | p.M1119I |
| EGF | 1950 | 4 | 110932470 | 110932470 | Missense Mut | SNP | G | G | C | COG-N-515-D | g.chr4:110932470G>C | ENST00000509793.1 | + | 23 | 3809 | c.3357G>C | c.(3355-3357)atG>atC | p.M1119I |
| CELSR3 | 1951 | 3 | 48683048 | 48683048 | Missense Mut | SNP | A | A | T | COG-N-512-D | g.chr3:48683048A>T | ENST00000164024.4 | - | 24 | 7905 | c.7625T>A | c.(7624-7626)tTc>tAc | p.F2542Y |
| CELSR3 | 1951 | 3 | 48683048 | 48683048 | Missense Mut | SNP | A | A | T | COG-N-515-D | g.chr3:48683048A>T | ENST00000164024.4 | - | 24 | 7905 | c.7625T>A | c.(7624-7626)tTc>tAc | p.F2542Y |
| CELSR2 | 1952 | 1 | 109813656 | 109813656 | Missense Mut | SNP | G | G | C | COG-N-512-D | g.chr1:109813656G>C | ENST00000271332.3 | + | 25 | 7652 | c.7591G>C | c.(7591-7593)Gtc>Ctc | p.V2531L |
| CELSR2 | 1952 | 1 | 109813656 | 109813656 | Missense Mut | SNP | G | G | C | COG-N-515-D | g.chr1:109813656G>C | ENST00000271332.3 | + | 25 | 7652 | c.7591G>C | c.(7591-7593)Gtc>Ctc | p.V2531L |
| FBN1 | 2200 | 15 | 48756108 | 48756108 | Missense Mut | SNP | T | T | A | COG-N-512-D | g.chr15:48756108T>A | ENST00000316623.5 | - | 41 | 5508 | c.5053A>T | c.(5053-5055)Aat>Tat | p.N1685Y |
| FBN1 | 2200 | 15 | 48756108 | 48756108 | Missense Mut | SNP | T | T | A | COG-N-515-D | g.chr15:48756108T>A | ENST00000316623.5 | - | 41 | 5508 | c.5053A>T | c.(5053-5055)Aat>Tat | p.N1685Y |
| FUCA2 | 2519 | 6 | 143832707 | 143832708 | In Frame Ins | INS | - | - | GCAGCA | COG-N-512-D | g.chr6:143832707 1438327 | ENST00000002165.6 | - | 1 | 119 120 | c.64 65insTGCT( | c.(64-66)ccg>cTGCTGCc | p.21 22insLL |
| FUCA2 | 2519 | 6 | 143832707 | 143832708 | In Frame Ins | INS | - | - | GCAGCA | COG-N-515-D | g.chr6:143832707 1438327 | ENST00000002165.6 | - | 1 | 119 120 | c.64 65insTGCT( | c.(64-66)ccg>cTGCTGCc | p.21 22insLL |
| GP1BA | 2811 | 17 | 4837220 | 4837220 | Missense Mut | SNP | T | T | C | COG-N-515-D | g.chr17:4837220T>C | ENST00000329125.5 | + | 2 | 1396 | c.1321T>C | c.(1321-1323)Tca>Cca | p.S441P |
| GSTM4 | 2948 | 1 | 110200243 | 110200243 | Missense Mut | SNP | T | T | C | COG-N-512-D | g.chr1:110200243T>C | ENST00000369836.4 | + | 4 | 518 | c.209T>C | c.(208-210)aTc>aCc | p.I70T |
| GSTM4 | 2948 | 1 | 110200243 | 110200243 | Missense Mut | SNP | T | T | C | COG-N-515-D | g.chr1:110200243T>C | ENST00000369836.4 | + | 4 | 518 | c.209T>C | c.(208-210)aTc>aCc | p.I70T |
| HEXB | 3074 | 5 | 73981105 | 73981105 | Missense Mut | SNP | G | G | C | COG-N-512-D | g.chr5:73981105G>C | ENST00000261416.7 | + | 1 | 137 | c.20G>C | c.(19-21)gGg>gCg | p.G7A |
| HEXB | 3074 | 5 | 73981105 | 73981105 | Missense Mut | SNP | G | G | C | COG-N-515-D | g.chr5:73981105G>C | ENST00000261416.7 | + | 1 | 137 | c.20G>C | c.(19-21)gGg>gCg | p.G7A |
| IMPDH2 | 3615 | 3 | 49062349 | 49062349 | Missense Mut | SNP | G | G | T | COG-N-512-D | g.chr3:49062349G>T | ENST00000326739.4 | - | 11 | 1314 | c.1275C>A | c.(1273-1275)agC>agA | p.S425R |
| IMPDH2 | 3615 | 3 | 49062349 | 49062349 | Missense Mut | SNP | G | G | T | COG-N-515-D | g.chr3:49062349G>T | ENST00000326739.4 | - | 11 | 1314 | c.1275C>A | c.(1273-1275)agC>agA | p.S425R |
| INPP5B | 3633 | 1 | 38331495 | 38331495 | Missense Mut | SNP | T | T | C | COG-N-512-D | g.chr1:38331495T>C | ENST00000373024.3 | - | 22 | 2539 | c.2452A>G | c.(2452-2454)Atc>Gtc | p.I818V |
| INPP5B | 3633 | 1 | 38331495 | 38331495 | Missense Mut | SNP | T | T | C | COG-N-515-D | g.chr1:38331495T>C | ENST00000373024.3 | - | 22 | 2539 | c.2452A>G | c.(2452-2454)Atc>Gtc | p.I818V |
| KCNN3 | 3782 | 1 | 154685999 | 154685999 | Missense Mut | SNP | C | C | T | COG-N-512-D | g.chr1:154685999C>T | ENST00000271915.4 | - | 7 | 2155 | c.1840G>A | c.(1840-1842)Gtc>Atc | p.V614I |
| KCNN3 | 3782 | 1 | 154685999 | 154685999 | Missense Mut | SNP | C | C | T | COG-N-515-D | g.chr1:154685999C>T | ENST00000271915.4 | - | 7 | 2155 | c.1840G>A | c.(1840-1842)Gtc>Atc | p.V614I |
| KCNS1 | 3787 | 20 | 43723701 | 43723701 | Missense Mut | SNP | T | T | G | COG-N-512-D | g.chr20:43723701T>G | ENST00000306117.1 | - | 5 | 1787 | c.1391A>C | c.(1390-1392)tAc>tCc | p.Y464S |
| KCNS2 | 3788 | 8 | 99440901 | 99440901 | Missense Mut | SNP | C | C | A | COG-N-512-D | g.chr8:99440901C>A | ENST00000287042.4 | + | 2 | 1044 | c.694C>A | c.(694-696)Cac>Aac | p.H232N |
| KCNS2 | 3788 | 8 | 99440901 | 99440901 | Missense Mut | SNP | C | C | A | COG-N-515-D | g.chr8:99440901C>A | ENST00000287042.4 | + | 2 | 1044 | c.694C>A | c.(694-696)Cac>Aac | p.H232N |
| LAMA2 | 3908 | 6 | 129371187 | 129371187 | Missense Mut | SNP | G | G | T | COG-N-512-D | g.chr6:129371187G>T | ENST00000421865.2 | + | 2 | 286 | c.237G>T | c.(235-237)agG>agT | p.R79S |
| LAMA2 | 3908 | 6 | 129371187 | 129371187 | Missense Mut | SNP | G | G | T | COG-N-515-D | g.chr6:129371187G>T | ENST00000421865.2 | + | 2 | 286 | c.237G>T | c.(235-237)agG>agT | p.R79S |
| VWA5A | 4013 | 11 | 123989052 | 123989052 | Missense Mut | SNP | C | C | G | COG-N-515-D | g.chr11:123989052C>G | ENST00000392744.4 | + | 4 | 1779 | c.451C>G | c.(451-453)Ctg>Gtg | p.L151V |
| MUC1 | 4582 | 1 | 155160884 | 155160884 | Missense Mut | SNP | G | G | C | COG-N-512-D | g.chr1:155160884G>C | ENST00000368395.1 | - | 3 | 714 | c.643C>G | c.(643-645)Ccc>Gcc | p.P215A |
| MUC1 | 4582 | 1 | 155160884 | 155160884 | Missense Mut | SNP | G | G | C | COG-N-515-D | g.chr1:155160884G>C | ENST00000368395.1 | - | 3 | 714 | c.643C>G | c.(643-645)Ccc>Gcc | p.P215A |
| MYO9B | 4650 | 19 | 17318011 | 17318011 | Missense Mut | SNP | C | C | T | COG-N-512-D | g.chr19:17318011C>T | ENST00000594824.1 | + | 35 | 5729 | c.5582C>T | c.(5581-5583)tCg>tTg | p.S1861L |
| MYO9B | 4650 | 19 | 17318011 | 17318011 | Missense Mut | SNP | C | C | T | COG-N-515-D | g.chr19:17318011C>T | ENST00000594824.1 | + | 35 | 5729 | c.5582C>T | c.(5581-5583)tCg>tTg | p.S1861L |
| NEFH | 4744 | 22 | 29885728 | 29885728 | Missense Mut | SNP | A | A | C | COG-N-515-D | g.chr22:29885728A>C | ENST00000310624.6 | + | 4 | 2132 | c.2099A>C | c.(2098-2100)gAa>gCa | p.E700A |
| NOS2 | 4843 | 17 | 26094736 | 26094736 | Missense Mut | SNP | C | C | A | COG-N-512-D | g.chr17:26094736C>A | ENST00000313735.6 | - | 18 | 2395 | c.2162G>T | c.(2161-2163)aGc>aTc | p.S721I |
| NOS2 | 4843 | 17 | 26094736 | 26094736 | Missense Mut | SNP | C | C | A | COG-N-515-D | g.chr17:26094736C>A | ENST00000313735.6 | - | 18 | 2395 | c.2162G>T | c.(2161-2163)aGc>aTc | p.S721I |
| NPR3 | 4883 | 5 | 32710775 | 32710775 | Missense Mut | SNP | C | C | A | COG-N-515-D | g.chr5:32710775C>A | ENST00000434067.2 | + | 1 | 1263 | c.7C>A | c.(7-9)Cca>Aca | p.P3T |
| OAS3 | 4940 | 12 | 113407528 | 113407528 | Missense Mut | SNP | A | A | G | COG-N-512-D | g.chr12:113407528A>G | ENST00000228928.7 | + | 15 | 3399 | c.3220A>G | c.(3220-3222)Aat>Gat | p.N1074D |
| OAS3 | 4940 | 12 | 113407528 | 113407528 | Missense Mut | SNP | A | A | G | COG-N-515-D | g.chr12:113407528A>G | ENST00000228928.7 | + | 15 | 3399 | c.3220A>G | c.(3220-3222)Aat>Gat | p.N1074D |
| OSBP | 5007 | 11 | 59383064 | 59383065 | In Frame Ins | INS | - | - | CGC | COG-N-512-D | g.chr11:59383064 5938306 | ENST00000263847.1 | - | 1 | 552 553 | c.73 74insGCG | c.(73-75)gcc>gGCGcc | p.24 25insG |
| PAX3 | 5077 | 2 | 223163294 | 223163294 | Missense Mut | SNP | G | G | T | COG-N-515-D | g.chr2:223163294G>T | ENST00000409551.3 | - | 1 | 320 | c.41C>A | c.(40-42)cCg>cAg | p.P14Q |
| PFKM | 5213 | 12 | 48535710 | 48535710 | Missense Mut | SNP | G | G | T | COG-N-515-D | g.chr12:48535710G>T | ENST00000312352.7 | + | 17 | 1553 | c.1514G>T | c.(1513-1515)gGc>gTc | p.G505V |
| PPID | 5481 | 4 | 159634297 | 159634297 | Nonsense Mut | SNP | G | G | A | COG-N-512-D | g.chr4:159634297G>A | ENST00000307720.3 | - | 7 | 975 | c.868C>T | c.(868-870)Cag>Tag | p.Q290* |
| PPID | 5481 | 4 | 159634297 | 159634297 | Nonsense Mut | SNP | G | G | A | COG-N-515-D | g.chr4:159634297G>A | ENST00000307720.3 | - | 7 | 975 | c.868C>T | c.(868-870)Cag>Tag | p.Q290* |
| PPL | 5493 | 16 | 4934418 | 4934418 | Missense Mut | SNP | C | C | T | COG-N-515-D | g.chr16:4934418C>T | ENST00000345988.2 | - | 22 | 4327 | c.4238G>A | c.(4237-4239)cGc>cAc | p.R1413H |
| PSMC5 | 5705 | 17 | 61908190 | 61908190 | Missense Mut | SNP | C | C | G | COG-N-512-D | g.chr17:61908190C>G | ENST00000310144.6 | + | 7 | 882 | c.574C>G | c.(574-576)Cca>Gca | p.P192A |
| PSMC5 | 5705 | 17 | 61908190 | 61908190 | Missense Mut | SNP | C | C | G | COG-N-515-D | g.chr17:61908190C>G | ENST00000310144.6 | + | 7 | 882 | c.574C>G | c.(574-576)Cca>Gca | p.P192A |
| PTPRS | 5802 | 19 | 5221206 | 5221206 | Missense Mut | SNP | G | G | A | COG-N-512-D | g.chr19:5221206G>A | ENST00000587303.1 | - | 19 | 3359 | c.3260C>T | c.(3259-3261)aCg>aTg | p.T1087M |
| PTPRS | 5802 | 19 | 5221206 | 5221206 | Missense Mut | SNP | G | G | A | COG-N-515-D | g.chr19:5221206G>A | ENST00000587303.1 | - | 19 | 3359 | c.3260C>T | c.(3259-3261)aCg>aTg | p.T1087M |
| RASGRF2 | 5924 | 5 | 80388680 | 80388680 | Missense Mut | SNP | C | C | T | COG-N-512-D | g.chr5:80388680C>T | ENST00000265080.4 | + | 10 | 1518 | c.1451C>T | c.(1450-1452)tCg>tTg | p.S484L |
| RASGRF2 | 5924 | 5 | 80388680 | 80388680 | Missense Mut | SNP | C | C | T | COG-N-515-D | g.chr5:80388680C>T | ENST00000265080.4 | + | 10 | 1518 | c.1451C>T | c.(1450-1452)tCg>tTg | p.S484L |
| RET | 5979 | 10 | 43613822 | 43613822 | Splice Site | SNP | G | G | C | COG-N-512-D | g.chr10:43613822G>C | ENST00000355710.3 | + | 13 | 2518 | c.2286G>C | c.(2284-2286)gaG>gaC | p.E762D |
| RET | 5979 | 10 | 43613822 | 43613822 | Splice Site | SNP | G | G | C | COG-N-515-D | g.chr10:43613822G>C | ENST00000355710.3 | + | 13 | 2518 | c.2286G>C | c.(2284-2286)gaG>gaC | p.E762D |
| SFRP2 | 6423 | 4 | 154709612 | 154709612 | Missense Mut | SNP | C | C | A | COG-N-512-D | g.chr4:154709612C>A | ENST00000274063.4 | - | 1 | 660 | c.376G>T | c.(376-378)Gcc>Tcc | p.A126S |
| SFRP2 | 6423 | 4 | 154709612 | 154709612 | Missense Mut | SNP | C | C | A | COG-N-515-D | g.chr4:154709612C>A | ENST00000274063.4 | - | 1 | 660 | c.376G>T | c.(376-378)Gcc>Tcc | p.A126S |
| ITSN1 | 6453 | 21 | 35144485 | 35144485 | Frame Shift D | DEL | G | G | - | COG-N-512-D | g.chr21:35144485delG | ENST00000399367.3 | + | 12 | 1426 | c.1163delG | c.(1162-1164)cggfs | p.R388fs |
| ITSN1 | 6453 | 21 | 35144485 | 35144485 | Frame Shift D | DEL | G | G | - | COG-N-515-D | g.chr21:35144485delG | ENST00000399367.3 | + | 12 | 1426 | c.1163delG | c.(1162-1164)cggfs | p.R388fs |
| SLC13A1 | 6561 | 7 | 122787303 | 122787303 | Missense Mut | SNP | C | C | T | COG-N-512-D | g.chr7:122787303C>T | ENST00000194130.2 | - | 7 | 761 | c.722G>A | c.(721-723)tGt>tAt | p.C241Y |
| SLC13A1 | 6561 | 7 | 122787303 | 122787303 | Missense Mut | SNP | C | C | T | COG-N-515-D | g.chr7:122787303C>T | ENST00000194130.2 | - | 7 | 761 | c.722G>A | c.(721-723)tGt>tAt | p.C241Y |
| SLC18A2 | 6571 | 10 | 119017347 | 119017347 | Missense Mut | SNP | A | A | G | COG-N-512-D | g.chr10:119017347A>G | ENST00000298472.5 | + | 10 | 1078 | c.935A>G | c.(934-936)gAg>gGg | p.E312G |
| CAPN15 | 6650 | 16 | 597532 | 597532 | Missense Mut | SNP | T | T | C | COG-N-512-D | g.chr16:597532T>C | ENST00000219611.2 | + | 4 | 1057 | c.694T>C | c.(694-696)Ttc>Ctc | p.F232L |
| STK10 | 6793 | 5 | 171520934 | 171520935 | In Frame Ins | INS | - | - | AGA | COG-N-512-D | g.chr5:171520934 1715209 | ENST00000176763.5 | - | 9 | 1378 1379 | c.1035 1036ins1 | c.(1033-1038)tctgag>tc | p.345 346insS |
| STK10 | 6793 | 5 | 171520934 | 171520935 | In Frame Ins | INS | - | - | AGA | COG-N-515-D | g.chr5:171520934 1715209 | ENST00000176763.5 | - | 9 | 1378 1379 | c.1035 1036ins1 | c.(1033-1038)tctgag>tc | p.345 346insS |
| AURKC | 6795 | 19 | 57743577 | 57743577 | Missense Mut | SNP | T | T | C | COG-N-512-D | g.chr19:57743577T>C | ENST00000302804.7 | + | 3 | 467 | c.281T>C | c.(280-282)aTc>aCc | p.I94T |
| AURKC | 6795 | 19 | 57743577 | 57743577 | Missense Mut | SNP | T | T | C | COG-N-515-D | g.chr19:57743577T>C | ENST00000302804.7 | + | 3 | 467 | c.281T>C | c.(280-282)aTc>aCc | p.I94T |
| SUPT6H | 6830 | 17 | 27008981 | 27008981 | Missense Mut | SNP | G | G | C | COG-N-515-D | g.chr17:27008981G>C | ENST00000314616.6 | + | 13 | 1863 | c.1580G>C | c.(1579-1581)tGc>tCc | p.C527S |
| TEK | 7010 | 9 | 27169576 | 27169576 | Missense Mut | SNP | T | T | C | COG-N-512-D | g.chr9:27169576T>C | ENST00000519097.1 | + | 3 | 715 | c.265T>C | c.(265-267)Tat>Cat | p.Y89H |
| TEK | 7010 | 9 | 27169576 | 27169576 | Missense Mut | SNP | T | T | C | COG-N-515-D | g.chr9:27169576T>C | ENST00000519097.1 | + | 3 | 715 | c.265T>C | c.(265-267)Tat>Cat | p.Y89H |
| TJP1 | 7082 | 15 | 30010805 | 30010805 | Missense Mut | SNP | C | C | A | COG-N-512-D | g.chr15:30010805C>A | ENST00000400011.2 | - | 21 | 3554 | c.3313G>T | c.(3313-3315)Gca>Tca | p.A1105S |
| TJP1 | 7082 | 15 | 30010805 | 30010805 | Missense Mut | SNP | C | C | A | COG-N-515-D | g.chr15:30010805C>A | ENST00000400011.2 | - | 21 | 3554 | c.3313G>T | c.(3313-3315)Gca>Tca | p.A1105S |
| TNFSF4 | 7292 | 1 | 173155692 | 173155692 | Missense Mut | SNP | A | A | G | COG-N-512-D | g.chr1:173155692A>G | ENST00000281834.3 | - | 3 | 651 | c.515T>C | c.(514-516)cTt>cCt | p.L172P |
| TNFSF4 | 7292 | 1 | 173155692 | 173155692 | Missense Mut | SNP | A | A | G | COG-N-515-D | g.chr1:173155692A>G | ENST00000281834.3 | - | 3 | 651 | c.515T>C | c.(514-516)cTt>cCt | p.L172P |
| ZNF141 | 7700 | 4 | 366732 | 366732 | Missense Mut | SNP | G | G | T | COG-N-512-D | g.chr4:366732G>T | ENST00000240499.7 | + | 4 | 655 | c.506G>T | c.(505-507)gGa>gTa | p.G169V |
| ZNF141 | 7700 | 4 | 366732 | 366732 | Missense Mut | SNP | G | G | T | COG-N-515-D | g.chr4:366732G>T | ENST00000240499.7 | + | 4 | 655 | c.506G>T | c.(505-507)gGa>gTa | p.G169V |
| NCOA3 | 8202 | 20 | 46262355 | 46262355 | Missense Mut | SNP | G | G | T | COG-N-512-D | g.chr20:46262355G>T | ENST00000371998.3 | + | 9 | 1130 | c.939G>T | c.(937-939)tgG>tgT | p.W313C |
| HIRIP3 | 8479 | 16 | 30004637 | 30004637 | Missense Mut | SNP | C | C | G | COG-N-512-D | g.chr16:30004637C>G | ENST00000279392.3 | - | 7 | 2392 | c.1562G>C | c.(1561-1563)gGg>gCg | p.G521A |
| HIRIP3 | 8479 | 16 | 30004637 | 30004637 | Missense Mut | SNP | C | C | G | COG-N-515-D | g.chr16:30004637C>G | ENST00000279392.3 | - | 7 | 2392 | c.1562G>C | c.(1561-1563)gGg>gCg | p.G521A |
| KRT36 | 8689 | 17 | 39643934 | 39643934 | Missense Mut | SNP | G | G | T | COG-N-512-D | g.chr17:39643934G>T | ENST00000328119.6 | - | 4 | 754 | c.755C>A | c.(754-756)gCt>gAt | p.A252D |
| KRT36 | 8689 | 17 | 39643934 | 39643934 | Missense Mut | SNP | G | G | T | COG-N-515-D | g.chr17:39643934G>T | ENST00000328119.6 | - | 4 | 754 | c.755C>A | c.(754-756)gCt>gAt | p.A252D |
| CACNA1G | 8913 | 17 | 48703987 | 48703995 | In Frame Del | DEL | TCCAGCG/ | TCCAGCGAC | - | COG-N-515-D | g.chr17:48703987 4870399 | ENST00000515411.1 | + | 36 | 6820 6828 | c.6820 6828del | c.(6820-6828)tccagcgac | p.SSD2274del |
| HSPB3 | 8988 | 5 | 53751869 | 53751869 | Missense Mut | SNP | G | G | C | COG-N-515-D | g.chr5:53751869G>C | ENST00000302005.1 | + | 1 | 425 | c.250G>C | c.(250-252)Gac>Cac | p.D84H |
| PAPSS1 | 9061 | 4 | 108535506 | 108535506 | Missense Mut | SNP | G | G | A | COG-N-512-D | g.chr4:108535506G>A | ENST00000265174.4 | - | 12 | 2046 | c.1774C>T | c.(1774-1776)Cgc>Tgc | p.R592C |
| PAPSS1 | 9061 | 4 | 108535506 | 108535506 | Missense Mut | SNP | G | G | A | COG-N-515-D | g.chr4:108535506G>A | ENST00000265174.4 | - | 12 | 2046 | c.1774C>T | c.(1774-1776)Cgc>Tgc | p.R592C |
| KCNQ4 | 9132 | 1 | 41296926 | 41296926 | Missense Mut | SNP | G | G | A | COG-N-512-D | g.chr1:41296926G>A | ENST00000347132.5 | + | 10 | 1545 | c.1463G>A | c.(1462-1464)cGc>cAc | p.R488H |
| KCNQ4 | 9132 | 1 | 41296926 | 41296926 | Missense Mut | SNP | G | G | A | COG-N-515-D | g.chr1:41296926G>A | ENST00000347132.5 | + | 10 | 1545 | c.1463G>A | c.(1462-1464)cGc>cAc | p.R488H |
| DLGAP2 | 9228 | 8 | 1616572 | 1616572 | Missense Mut | SNP | A | A | T | COG-N-512-D | g.chr8:1616572A>T | ENST00000421627.2 | + | 6 | 1782 | c.1648A>T | c.(1648-1650)Acg>Tcg | p.T550S |
| DLGAP2 | 9228 | 8 | 1616572 | 1616572 | Missense Mut | SNP | A | A | T | COG-N-515-D | g.chr8:1616572A>T | ENST00000421627.2 | + | 6 | 1782 | c.1648A>T | c.(1648-1650)Acg>Tcg | p.T550S |
| GTF3C5 | 9328 | 9 | 135926225 | 135926225 | Missense Mut | SNP | G | G | A | COG-N-512-D | g.chr9:135926225G>A | ENST00000372097.5 | + | 4 | 951 | c.628G>A | c.(628-630)Gcc>Acc | p.A210T |
| GTF3C5 | 9328 | 9 | 135926225 | 135926225 | Missense Mut | SNP | G | G | A | COG-N-515-D | g.chr9:135926225G>A | ENST00000372097.5 | + | 4 | 951 | c.628G>A | c.(628-630)Gcc>Acc | p.A210T |
| THEMIS2 | 9473 | 1 | 28211836 | 28211836 | Missense Mut | SNP | G | G | A | COG-N-512-D | g.chr1:28211836G>A | ENST00000373921.3 | + | 5 | 1754 | c.1750G>A | c.(1750-1752)Gct>Act | p.A584T |
| THEMIS2 | 9473 | 1 | 28211836 | 28211836 | Missense Mut | SNP | G | G | A | COG-N-515-D | g.chr1:28211836G>A | ENST00000373921.3 | + | 5 | 1754 | c.1750G>A | c.(1750-1752)Gct>Act | p.A584T |
| H6PD | 9563 | 1 | 9305325 | 9305325 | Missense Mut | SNP | T | T | C | COG-N-512-D | g.chr1:9305325T>C | ENST00000377403.2 | + | 2 | 634 | c.332T>C | c.(331-333)cTg>cCg | p.L111P |
| H6PD | 9563 | 1 | 9305325 | 9305325 | Missense Mut | SNP | T | T | C | COG-N-515-D | g.chr1:9305325T>C | ENST00000377403.2 | + | 2 | 634 | c.332T>C | c.(331-333)cTg>cCg | p.L111P |
| EDEM1 | 9695 | 3 | 5229746 | 5229746 | Missense Mut | SNP | G | G | C | COG-N-515-D | g.chr3:5229746G>C | ENST00000256497.4 | + | 1 | 389 | c.256G>C | c.(256-258)Gct>Cct | p.A86P |
| SART3 | 9733 | 12 | 108926042 | 108926042 | Missense Mut | SNP | G | G | T | COG-N-512-D | g.chr12:108926042G>T | ENST00000228284.3 | - | 13 | 1859 | c.1625C>A | c.(1624-1626)cCa>cAa | p.P542Q |
| SART3 | 9733 | 12 | 108926042 | 108926042 | Missense Mut | SNP | G | G | T | COG-N-515-D | g.chr12:108926042G>T | ENST00000228284.3 | - | 13 | 1859 | c.1625C>A | c.(1624-1626)cCa>cAa | p.P542Q |
| FRMPD4 | 9758 | X | 12735693 | 12735693 | Missense Mut | SNP | G | G | T | COG-N-515-D | g.chrX:12735693G>T | ENST00000380682.1 | + | 16 | 3254 | c.2748G>T | c.(2746-2748)atG>atT | p.M916I |
| KIAA0753 | 9851 | 17 | 6524118 | 6524118 | Missense Mut | SNP | C | C | A | COG-N-512-D | g.chr17:6524118C>A | ENST00000361413.3 | - | 7 | 1663 | c.1305G>T | c.(1303-1305)caG>caT | p.Q435H |
| KIAA0753 | 9851 | 17 | 6524118 | 6524118 | Missense Mut | SNP | C | C | A | COG-N-515-D | g.chr17:6524118C>A | ENST00000361413.3 | - | 7 | 1663 | c.1305G>T | c.(1303-1305)caG>caT | p.Q435H |
| MRC2 | 9902 | 17 | 60759300 | 60759300 | Splice Site | SNP | T | T | C | COG-N-512-D | g.chr17:60759300T>C | ENST00000303375.5 | + | 19 | 3205 | | c.e19+2 | |
| ARNT2 | 9915 | 15 | 80885968 | 80885968 | Missense Mut | SNP | A | A | G | COG-N-512-D | g.chr15:80885968A>G | ENST00000303329.4 | + | 19 | 2248 | c.2083A>G | c.(2083-2085)Acc>Gcc | p.T695A |
| ARNT2 | 9915 | 15 | 80885968 | 80885968 | Missense Mut | SNP | A | A | G | COG-N-515-D | g.chr15:80885968A>G | ENST00000303329.4 | + | 19 | 2248 | c.2083A>G | c.(2083-2085)Acc>Gcc | p.T695A |
| USP15 | 9958 | 12 | 62790148 | 62790148 | Missense Mut | SNP | A | A | G | COG-N-512-D | g.chr12:62790148A>G | ENST00000280377.5 | + | 20 | 2702 | c.2644A>G | c.(2644-2646)Aac>Gac | p.N882D |
| USP15 | 9958 | 12 | 62790148 | 62790148 | Missense Mut | SNP | A | A | G | COG-N-515-D | g.chr12:62790148A>G | ENST00000280377.5 | + | 20 | 2702 | c.2644A>G | c.(2644-2646)Aac>Gac | p.N882D |
| NR1H3 | 10062 | 11 | 47281995 | 47281995 | Missense Mut | SNP | C | C | T | COG-N-515-D | g.chr11:47281995C>T | ENST00000467728.1 | + | 3 | 1506 | c.268C>T | c.(268-270)Ccc>Tcc | p.P90S |
| COQ7 | 10229 | 16 | 19087122 | 19087122 | Nonsense Mut | SNP | C | C | A | COG-N-512-D | g.chr16:19087122C>A | ENST00000569127.1 | + | 4 | 1264 | c.378C>A | c.(376-378)taC>taA | p.Y126* |
| COQ7 | 10229 | 16 | 19087122 | 19087122 | Nonsense Mut | SNP | C | C | A | COG-N-515-D | g.chr16:19087122C>A | ENST00000569127.1 | + | 4 | 1264 | c.378C>A | c.(376-378)taC>taA | p.Y126* |
| MYL9 | 10398 | 20 | 35173325 | 35173325 | Missense Mut | SNP | A | A | G | COG-N-512-D | g.chr20:35173325A>G | ENST00000279022.2 | + | 2 | 142 | c.38A>G | c.(37-39)aAg>aGg | p.K13R |
| MYL9 | 10398 | 20 | 35173325 | 35173325 | Missense Mut | SNP | A | A | G | COG-N-515-D | g.chr20:35173325A>G | ENST00000279022.2 | + | 2 | 142 | c.38A>G | c.(37-39)aAg>aGg | p.K13R |
| NCOA2 | 10499 | 8 | 71044183 | 71044183 | Missense Mut | SNP | C | C | A | COG-N-512-D | g.chr8:71044183C>A | ENST00000452400.2 | - | 16 | 3394 | c.3213G>T | c.(3211-3213)gaG>gaT | p.E1071D |
| NCOA2 | 10499 | 8 | 71044183 | 71044183 | Missense Mut | SNP | C | C | A | COG-N-515-D | g.chr8:71044183C>A | ENST00000452400.2 | - | 16 | 3394 | c.3213G>T | c.(3211-3213)gaG>gaT | p.E1071D |
| FBXW10 | 10517 | 17 | 18653104 | 18653104 | Missense Mut | SNP | T | T | A | COG-N-512-D | g.chr17:18653104T>A | ENST00000395665.4 | + | 3 | 961 | c.740T>A | c.(739-741)aTa>aAa | p.I247K |
| IPO7 | 10527 | 11 | 9444613 | 9444613 | Nonsense Mut | SNP | C | C | T | COG-N-515-D | g.chr11:9444613C>T | ENST00000379719.3 | + | 9 | 1109 | c.967C>T | c.(967-969)Caa>Taa | p.Q323* |
| SLC19A2 | 10560 | 1 | 169435094 | 169435094 | Missense Mut | SNP | G | G | A | COG-N-512-D | g.chr1:169435094G>A | ENST00000236137.5 | - | 6 | 1723 | c.1487C>T | c.(1486-1488)aCt>aTt | p.T496I |
| SLC19A2 | 10560 | 1 | 169435094 | 169435094 | Missense Mut | SNP | G | G | A | COG-N-515-D | g.chr1:169435094G>A | ENST00000236137.5 | - | 6 | 1723 | c.1487C>T | c.(1486-1488)aCt>aTt | p.T496I |
| COLEC10 | 10584 | 8 | 120118124 | 120118124 | Nonsense Mut | SNP | C | C | A | COG-N-512-D | g.chr8:120118124C>A | ENST00000332843.2 | + | 6 | 569 | c.528C>A | c.(526-528)tgC>tgA | p.C176* |
| COLEC10 | 10584 | 8 | 120118124 | 120118124 | Nonsense Mut | SNP | C | C | A | COG-N-515-D | g.chr8:120118124C>A | ENST00000332843.2 | + | 6 | 569 | c.528C>A | c.(526-528)tgC>tgA | p.C176* |
| RBCK1 | 10616 | 20 | 409676 | 409676 | Missense Mut | SNP | C | C | T | COG-N-512-D | g.chr20:409676C>T | ENST00000356286.5 | + | 11 | 2095 | c.1390C>T | c.(1390-1392)Cgc>Tgc | p.R464C |
| RBCK1 | 10616 | 20 | 409676 | 409676 | Missense Mut | SNP | C | C | T | COG-N-515-D | g.chr20:409676C>T | ENST00000356286.5 | + | 11 | 2095 | c.1390C>T | c.(1390-1392)Cgc>Tgc | p.R464C |
| LIAS | 11019 | 4 | 39462537 | 39462537 | Missense Mut | SNP | C | C | T | COG-N-512-D | g.chr4:39462537C>T | ENST00000513731.1 | + | 2 | 225 | c.173C>T | c.(172-174)aCc>aTc | p.T58I |
| LIAS | 11019 | 4 | 39462537 | 39462537 | Missense Mut | SNP | C | C | T | COG-N-515-D | g.chr4:39462537C>T | ENST00000513731.1 | + | 2 | 225 | c.173C>T | c.(172-174)aCc>aTc | p.T58I |
| LILRA1 | 11024 | 19 | 55105908 | 55105908 | Missense Mut | SNP | C | C | A | COG-N-512-D | g.chr19:55105908C>A | ENST00000251372.3 | + | 3 | 219 | c.37C>A | c.(37-39)Ctg>Atg | p.L13M |
| LILRA1 | 11024 | 19 | 55105908 | 55105908 | Missense Mut | SNP | C | C | A | COG-N-515-D | g.chr19:55105908C>A | ENST00000251372.3 | + | 3 | 219 | c.37C>A | c.(37-39)Ctg>Atg | p.L13M |
| B4GALT7 | 11285 | 5 | 177034442 | 177034442 | Missense Mut | SNP | G | G | A | COG-N-512-D | g.chr5:177034442G>A | ENST00000029410.5 | + | 3 | 664 | c.553G>A | c.(553-555)Gtg>Atg | p.V185M |
| B4GALT7 | 11285 | 5 | 177034442 | 177034442 | Missense Mut | SNP | G | G | A | COG-N-515-D | g.chr5:177034442G>A | ENST00000029410.5 | + | 3 | 664 | c.553G>A | c.(553-555)Gtg>Atg | p.V185M |
| U2AF2 | 11338 | 19 | 56180474 | 56180474 | Missense Mut | SNP | A | A | T | COG-N-515-D | g.chr19:56180474A>T | ENST00000308924.4 | + | 10 | 1011 | c.971A>T | c.(970-972)cAg>cTg | p.Q324L |
| SCAF8 | 22828 | 6 | 155055051 | 155055051 | Missense Mut | SNP | C | C | T | COG-N-512-D | g.chr6:155055051C>T | ENST00000367178.3 | + | 1 | 593 | c.17C>T | c.(16-18)aCc>aTc | p.T6I |
| SCAF8 | 22828 | 6 | 155055051 | 155055051 | Missense Mut | SNP | C | C | T | COG-N-515-D | g.chr6:155055051C>T | ENST00000367178.3 | + | 1 | 593 | c.17C>T | c.(16-18)aCc>aTc | p.T6I |
| SLC4A1AP | 22950 | 2 | 27905218 | 27905218 | Missense Mut | SNP | A | A | T | COG-N-512-D | g.chr2:27905218A>T | ENST00000326019.6 | + | 9 | 2149 | c.1867A>T | c.(1867-1869)Aca>Tca | p.T623S |

| | | | | | | | | | | | | | | | | | | | | |
|---|---|---|---|---|---|---|---|---|---|---|---|---|---|---|---|---|---|---|---|---|
| PCNX | 22990 | 14 | 71455295 | 71455295 | Missense Mut | SNP | A | A | T | COG-N-512-D | g.chr14:71455295A>T | ENST00000304743.2 | + | 7 | | | 2769 | c.2323A>T | c.(2323-2325)Agc>Tgc | p.S775C |
| PCNX | 22990 | 14 | 71455295 | 71455295 | Missense Mut | SNP | A | A | T | COG-N-515-D | g.chr14:71455295A>T | ENST00000304743.2 | + | 7 | | | 2769 | c.2323A>T | c.(2323-2325)Agc>Tgc | p.S775C |
| TNRC6B | 23112 | 22 | 40681746 | 40681746 | Missense Mut | SNP | A | A | T | COG-N-515-D | g.chr22:40681746A>T | ENST00000301923.9 | + | 13 | | | 1570 | c.1268A>T | c.(1267-1269)cAa>cTa | p.Q423L |
| RAD54L2 | 23132 | 3 | 51691635 | 51691635 | Missense Mut | SNP | A | A | G | COG-N-512-D | g.chr3:51691635A>G | ENST00000409535.2 | + | 20 | | | 3391 | c.3266A>G | c.(3265-3267)cAg>cGg | p.Q1089R |
| RAD54L2 | 23132 | 3 | 51691635 | 51691635 | Missense Mut | SNP | A | A | G | COG-N-515-D | g.chr3:51691635A>G | ENST00000409535.2 | + | 20 | | | 3391 | c.3266A>G | c.(3265-3267)cAg>cGg | p.Q1089R |
| EFR3A | 23167 | 8 | 132958758 | 132958758 | Missense Mut | SNP | G | G | A | COG-N-512-D | g.chr8:132958758G>A | ENST00000254624.5 | + | 4 | | | 469 | c.244G>A | c.(244-246)Gac>Aac | p.D82N |
| EFR3A | 23167 | 8 | 132958758 | 132958758 | Missense Mut | SNP | G | G | A | COG-N-515-D | g.chr8:132958758G>A | ENST00000254624.5 | + | 4 | | | 469 | c.244G>A | c.(244-246)Gac>Aac | p.D82N |
| SYNE1 | 23345 | 6 | 152771851 | 152771851 | Missense Mut | SNP | G | G | T | COG-N-512-D | g.chr6:152771851G>T | ENST00000367255.5 | - | 27 | | | 3905 | c.3304C>A | c.(3304-3306)Cac>Aac | p.H1102N |
| SYNE1 | 23345 | 6 | 152771851 | 152771851 | Missense Mut | SNP | G | G | T | COG-N-515-D | g.chr6:152771851G>T | ENST00000367255.5 | - | 27 | | | 3905 | c.3304C>A | c.(3304-3306)Cac>Aac | p.H1102N |
| GRIP1 | 23426 | 12 | 66765616 | 66765616 | Missense Mut | SNP | C | C | T | COG-N-512-D | g.chr12:66765616C>T | ENST00000398016.3 | - | 22 | | | 2782 | c.2714G>A | c.(2713-2715)cGc>cAc | p.R905H |
| GRIP1 | 23426 | 12 | 66765616 | 66765616 | Missense Mut | SNP | C | C | T | COG-N-515-D | g.chr12:66765616C>T | ENST00000398016.3 | - | 22 | | | 2782 | c.2714G>A | c.(2713-2715)cGc>cAc | p.R905H |
| EDC4 | 23644 | 16 | 67915980 | 67915980 | Missense Mut | SNP | T | T | C | COG-N-512-D | g.chr16:67915980T>C | ENST00000358933.5 | + | 23 | | | 3367 | c.3128T>C | c.(3127-3129)aTa>aCa | p.I1043T |
| EDC4 | 23644 | 16 | 67915980 | 67915980 | Missense Mut | SNP | T | T | C | COG-N-515-D | g.chr16:67915980T>C | ENST00000358933.5 | + | 23 | | | 3367 | c.3128T>C | c.(3127-3129)aTa>aCa | p.I1043T |
| POLA2 | 23649 | 11 | 65063037 | 65063037 | Missense Mut | SNP | G | G | C | COG-N-515-D | g.chr11:65063037G>C | ENST00000265465.3 | + | 16 | | | 2018 | c.1487G>C | c.(1486-1488)aGc>aCc | p.S496T |
| RAD54B | 25788 | 8 | 95423425 | 95423425 | Nonsense Mut | SNP | C | C | T | COG-N-512-D | g.chr8:95423425C>T | ENST00000336148.5 | - | 4 | | | 547 | c.423G>A | c.(421-423)tgG>tgA | p.W141* |
| RAD54B | 25788 | 8 | 95423425 | 95423425 | Nonsense Mut | SNP | C | C | T | COG-N-515-D | g.chr8:95423425C>T | ENST00000336148.5 | - | 4 | | | 547 | c.423G>A | c.(421-423)tgG>tgA | p.W141* |
| SPDEF | 25803 | 6 | 34508791 | 34508791 | Missense Mut | SNP | G | G | T | COG-N-512-D | g.chr6:34508791G>T | ENST00000374037.3 | - | 3 | | | 1018 | c.604C>A | c.(604-606)Ctg>Atg | p.L202M |
| SPDEF | 25803 | 6 | 34508911 | 34508911 | Missense Mut | SNP | T | T | G | COG-N-512-D | g.chr6:34508911T>G | ENST00000374037.3 | - | 3 | | | 898 | c.484A>C | c.(484-486)Aca>Cca | p.T162P |
| SPDEF | 25803 | 6 | 34508911 | 34508911 | Missense Mut | SNP | T | T | G | COG-N-515-D | g.chr6:34508911T>G | ENST00000374037.3 | - | 3 | | | 898 | c.484A>C | c.(484-486)Aca>Cca | p.T162P |
| SPDEF | 25803 | 6 | 34508791 | 34508791 | Missense Mut | SNP | G | G | T | COG-N-515-D | g.chr6:34508791G>T | ENST00000374037.3 | - | 3 | | | 1018 | c.604C>A | c.(604-606)Ctg>Atg | p.L202M |
| DHRS7B | 25979 | 17 | 21075462 | 21075462 | Missense Mut | SNP | G | G | A | COG-N-512-D | g.chr17:21075462G>A | ENST00000395511.3 | + | 2 | | | 472 | c.152G>A | c.(151-153)cGg>cAg | p.R51Q |
| DHRS7B | 25979 | 17 | 21075462 | 21075462 | Missense Mut | SNP | G | G | A | COG-N-515-D | g.chr17:21075462G>A | ENST00000395511.3 | + | 2 | | | 472 | c.152G>A | c.(151-153)cGg>cAg | p.R51Q |
| FBXO25 | 26260 | 8 | 385667 | 385667 | Missense Mut | SNP | G | G | A | COG-N-512-D | g.chr8:385667G>A | ENST00000350302.3 | + | 5 | | | 607 | c.341G>A | c.(340-342)aGt>aAt | p.S114N |
| FBXO25 | 26260 | 8 | 385667 | 385667 | Missense Mut | SNP | G | G | A | COG-N-515-D | g.chr8:385667G>A | ENST00000350302.3 | + | 5 | | | 607 | c.341G>A | c.(340-342)aGt>aAt | p.S114N |
| ERAL1 | 26284 | 17 | 27182179 | 27182179 | Missense Mut | SNP | T | T | C | COG-N-512-D | g.chr17:27182179T>C | ENST00000254928.5 | + | 1 | | | 224 | c.127T>C | c.(127-129)Tgc>Cgc | p.C43R |
| ERAL1 | 26284 | 17 | 27182179 | 27182179 | Missense Mut | SNP | T | T | C | COG-N-515-D | g.chr17:27182179T>C | ENST00000254928.5 | + | 1 | | | 224 | c.127T>C | c.(127-129)Tgc>Cgc | p.C43R |
| HSPB8 | 26353 | 12 | 119617367 | 119617367 | Missense Mut | SNP | G | G | A | COG-N-512-D | g.chr12:119617367G>A | ENST00000281938.2 | + | 1 | | | 921 | c.250G>A | c.(250-252)Gag>Aag | p.E84K |
| HSPB8 | 26353 | 12 | 119617367 | 119617367 | Missense Mut | SNP | G | G | A | COG-N-515-D | g.chr12:119617367G>A | ENST00000281938.2 | + | 1 | | | 921 | c.250G>A | c.(250-252)Gag>Aag | p.E84K |
| TMPRSS11E | 28983 | 4 | 69343348 | 69343348 | Splice Site | SNP | T | T | G | COG-N-512-D | g.chr4:69343348T>G | ENST00000305363.4 | + | 8 | | | 1031 | | c.e8+2 | |
| TMPRSS11E | 28983 | 4 | 69343348 | 69343348 | Splice Site | SNP | T | T | G | COG-N-515-D | g.chr4:69343348T>G | ENST00000305363.4 | + | 8 | | | 1031 | | c.e8+2 | |
| TRPM5 | 29850 | 11 | 2434115 | 2434115 | Missense Mut | SNP | A | A | G | COG-N-512-D | g.chr11:2434115A>G | ENST00000155858.6 | - | 15 | | | 2232 | c.2224T>C | c.(2224-2226)Tac>Cac | p.Y742H |
| TRPM5 | 29850 | 11 | 2434115 | 2434115 | Missense Mut | SNP | A | A | G | COG-N-515-D | g.chr11:2434115A>G | ENST00000155858.6 | - | 15 | | | 2232 | c.2224T>C | c.(2224-2226)Tac>Cac | p.Y742H |
| EPN1 | 29924 | 19 | 56203166 | 56203171 | In Frame Del | DEL | CCCCGA | CCCCGA | - | COG-N-512-D | g.chr19:56203166 5620317 | ENST00000270460.6 | + | 7 | 1120 | 1125 | | c.809 814delCC( | c.(808-816)gccccgacc>g | p.PT271del |
| EPN1 | 29924 | 19 | 56203166 | 56203171 | In Frame Del | DEL | CCCCGA | CCCCGA | - | COG-N-515-D | g.chr19:56203166 5620317 | ENST00000270460.6 | + | 7 | 1120 | 1125 | | c.809 814delCC( | c.(808-816)gccccgacc>g | p.PT271del |
| SCAPER | 49855 | 15 | 76668643 | 76668643 | Missense Mut | SNP | C | C | T | COG-N-512-D | g.chr15:76668643C>T | ENST00000563290.1 | - | 29 | | | 3810 | c.3715G>A | c.(3715-3717)Ggg>Agg | p.G1239R |
| RDH8 | 50700 | 19 | 10132327 | 10132327 | Missense Mut | SNP | A | A | C | COG-N-515-D | g.chr19:10132327A>C | ENST00000591589.1 | + | 6 | | | 1087 | c.898A>C | c.(898-900)Acg>Ccg | p.T300P |
| MLXIPL | 51085 | 7 | 73011238 | 73011238 | Missense Mut | SNP | G | G | C | COG-N-515-D | g.chr7:73011238G>C | ENST00000313375.3 | - | 10 | | | 1675 | c.1628C>G | c.(1627-1629)cCa>cGa | p.P543R |
| ADIPOR1 | 51094 | 1 | 202913030 | 202913030 | Missense Mut | SNP | C | C | A | COG-N-512-D | g.chr1:202913030C>A | ENST00000340990.5 | - | 6 | | | 959 | c.661G>T | c.(661-663)Gtc>Ttc | p.V221F |
| PIPOX | 51268 | 17 | 27382139 | 27382139 | Missense Mut | SNP | G | G | A | COG-N-512-D | g.chr17:27382139G>A | ENST00000323372.4 | + | 6 | | | 1192 | c.866G>A | c.(865-867)cGc>cAc | p.R289H |
| PIPOX | 51268 | 17 | 27382139 | 27382139 | Missense Mut | SNP | G | G | A | COG-N-515-D | g.chr17:27382139G>A | ENST00000323372.4 | + | 6 | | | 1192 | c.866G>A | c.(865-867)cGc>cAc | p.R289H |
| PCF11 | 51585 | 11 | 82880861 | 82880861 | Missense Mut | SNP | C | C | G | COG-N-512-D | g.chr11:82880861C>G | ENST00000298281.4 | + | 8 | | | 3936 | c.3484C>G | c.(3484-3486)Cct>Gct | p.P1162A |
| PCF11 | 51585 | 11 | 82880861 | 82880861 | Missense Mut | SNP | C | C | G | COG-N-515-D | g.chr11:82880861C>G | ENST00000298281.4 | + | 8 | | | 3936 | c.3484C>G | c.(3484-3486)Cct>Gct | p.P1162A |
| NBAS | 51594 | 2 | 15493744 | 15493744 | Missense Mut | SNP | G | G | A | COG-N-512-D | g.chr2:15493744G>A | ENST00000281513.5 | - | 34 | | | 4047 | c.4022C>T | c.(4021-4023)gCt>gTt | p.A1341V |
| NBAS | 51594 | 2 | 15493744 | 15493744 | Missense Mut | SNP | G | G | A | COG-N-515-D | g.chr2:15493744G>A | ENST00000281513.5 | - | 34 | | | 4047 | c.4022C>T | c.(4021-4023)gCt>gTt | p.A1341V |
| PIGT | 51604 | 20 | 44054459 | 44054459 | Missense Mut | SNP | C | C | A | COG-N-512-D | g.chr20:44054459C>A | ENST00000279036.6 | + | 12 | | | 1810 | c.1730C>A | c.(1729-1731)cCa>cAa | p.P577Q |
| PIGT | 51604 | 20 | 44054459 | 44054459 | Missense Mut | SNP | C | C | A | COG-N-515-D | g.chr20:44054459C>A | ENST00000279036.6 | + | 12 | | | 1810 | c.1730C>A | c.(1729-1731)cCa>cAa | p.P577Q |
| RSF1 | 51773 | 11 | 77436649 | 77436649 | Missense Mut | SNP | C | C | G | COG-N-512-D | g.chr11:77436649C>G | ENST00000308488.6 | - | 5 | | | 976 | c.674G>C | c.(673-675)cGg>cCg | p.R225P |
| GALNT7 | 51809 | 4 | 174219438 | 174219438 | Missense Mut | SNP | G | G | C | COG-N-512-D | g.chr4:174219438G>C | ENST00000265000.4 | + | 6 | | | 1221 | c.1138G>C | c.(1138-1140)Gaa>Caa | p.E380Q |
| GALNT7 | 51809 | 4 | 174219438 | 174219438 | Missense Mut | SNP | G | G | C | COG-N-515-D | g.chr4:174219438G>C | ENST00000265000.4 | + | 6 | | | 1221 | c.1138G>C | c.(1138-1140)Gaa>Caa | p.E380Q |
| ANLN | 54443 | 7 | 36435896 | 36435896 | Missense Mut | SNP | G | G | T | COG-N-512-D | g.chr7:36435896G>T | ENST00000265748.2 | + | 2 | | | 261 | c.40G>T | c.(40-42)Gcc>Tcc | p.A14S |
| ANLN | 54443 | 7 | 36435896 | 36435896 | Missense Mut | SNP | G | G | T | COG-N-515-D | g.chr7:36435896G>T | ENST00000265748.2 | + | 2 | | | 261 | c.40G>T | c.(40-42)Gcc>Tcc | p.A14S |
| FAM134B | 54463 | 5 | 16617014 | 16617014 | Frame Shift D | DEL | C | C | - | COG-N-512-D | g.chr5:16617014delC | ENST00000306320.9 | - | 1 | | | 153 | c.67delG | c.(67-69)gcgfs | p.A23fs |
| USP53 | 54532 | 4 | 120192740 | 120192740 | Missense Mut | SNP | C | C | A | COG-N-515-D | g.chr4:120192740C>A | ENST00000274030.6 | + | 16 | | | 2904 | c.1725C>A | c.(1723-1725)agC>agA | p.S575R |
| SDK2 | 54549 | 17 | 71419527 | 71419527 | Missense Mut | SNP | T | T | C | COG-N-512-D | g.chr17:71419527T>C | ENST00000392650.3 | - | 14 | | | 1895 | c.1895A>G | c.(1894-1896)aAc>aGc | p.N632S |
| SDK2 | 54549 | 17 | 71419527 | 71419527 | Missense Mut | SNP | T | T | C | COG-N-515-D | g.chr17:71419527T>C | ENST00000392650.3 | - | 14 | | | 1895 | c.1895A>G | c.(1894-1896)aAc>aGc | p.N632S |
| CNGB3 | 54714 | 8 | 87616339 | 87616339 | Missense Mut | SNP | G | G | A | COG-N-512-D | g.chr8:87616339G>A | ENST00000320005.5 | - | 15 | | | 1810 | c.1763C>T | c.(1762-1764)tCg>tTg | p.S588L |
| CNGB3 | 54714 | 8 | 87616339 | 87616339 | Missense Mut | SNP | G | G | A | COG-N-515-D | g.chr8:87616339G>A | ENST00000320005.5 | - | 15 | | | 1810 | c.1763C>T | c.(1762-1764)tCg>tTg | p.S588L |
| NUTM2F | 54754 | 9 | 97080947 | 97080947 | Missense Mut | SNP | A | A | G | COG-N-515-D | g.chr9:97080947A>G | ENST00000253262.4 | - | 7 | | | 2091 | c.2071T>C | c.(2071-2073)Tct>Cct | p.S691P |
| CDHR2 | 54825 | 5 | 176004663 | 176004663 | Missense Mut | SNP | A | A | C | COG-N-512-D | g.chr5:176004663A>C | ENST00000510636.1 | + | 14 | | | 1650 | c.1376A>C | c.(1375-1377)cAg>cCg | p.Q459P |
| CDHR2 | 54825 | 5 | 176004663 | 176004663 | Missense Mut | SNP | A | A | C | COG-N-515-D | g.chr5:176004663A>C | ENST00000510636.1 | + | 14 | | | 1650 | c.1376A>C | c.(1375-1377)cAg>cCg | p.Q459P |
| ELP6 | 54859 | 3 | 47537644 | 47537644 | Missense Mut | SNP | C | C | T | COG-N-512-D | g.chr3:47537644C>T | ENST00000446787.1 | - | 6 | | | 685 | c.497G>A | c.(496-498)cGg>cAg | p.R166Q |
| ELP6 | 54859 | 3 | 47537644 | 47537644 | Missense Mut | SNP | C | C | T | COG-N-515-D | g.chr3:47537644C>T | ENST00000446787.1 | - | 6 | | | 685 | c.497G>A | c.(496-498)cGg>cAg | p.R166Q |
| RETSAT | 54884 | 2 | 85571717 | 85571717 | Splice Site | SNP | G | G | A | COG-N-512-D | g.chr2:85571717G>A | ENST00000295802.4 | - | 7 | | | 1368 | c.1256C>T | c.(1255-1257)gCg>gTg | p.A419V |
| RETSAT | 54884 | 2 | 85571717 | 85571717 | Splice Site | SNP | G | G | A | COG-N-515-D | g.chr2:85571717G>A | ENST00000295802.4 | - | 7 | | | 1368 | c.1256C>T | c.(1255-1257)gCg>gTg | p.A419V |
| UBE2R2 | 54926 | 9 | 33917176 | 33917178 | In Frame Del | DEL | GAG | GAG | - | COG-N-515-D | g.chr9:33917176 33917178 | ENST00000263228.3 | + | 5 | 849 | 851 | | c.658 660delGA | c.(658-660)gagdel | p.E223del |
| BANP | 54971 | 16 | 88039749 | 88039749 | Missense Mut | SNP | A | A | C | COG-N-512-D | g.chr16:88039749A>C | ENST00000393207.1 | + | 6 | | | 730 | c.509A>C | c.(508-510)gAa>gCa | p.E170A |
| BANP | 54971 | 16 | 88039749 | 88039749 | Missense Mut | SNP | A | A | C | COG-N-515-D | g.chr16:88039749A>C | ENST00000393207.1 | + | 6 | | | 730 | c.509A>C | c.(508-510)gAa>gCa | p.E170A |
| WDYHV1 | 55093 | 8 | 124442251 | 124442251 | Missense Mut | SNP | C | C | T | COG-N-512-D | g.chr8:124442251C>T | ENST00000523356.1 | + | 3 | | | 249 | c.212C>T | c.(211-213)cCt>cTt | p.P71L |
| WDYHV1 | 55093 | 8 | 124442251 | 124442251 | Missense Mut | SNP | C | C | T | COG-N-515-D | g.chr8:124442251C>T | ENST00000523356.1 | + | 3 | | | 249 | c.212C>T | c.(211-213)cCt>cTt | p.P71L |
| TMEM39B | 55116 | 1 | 32557521 | 32557521 | Missense Mut | SNP | A | A | G | COG-N-512-D | g.chr1:32557521A>G | ENST00000336294.5 | + | 6 | | | 982 | c.836A>G | c.(835-837)aAg>aGg | p.K279R |
| TMEM39B | 55116 | 1 | 32557521 | 32557521 | Missense Mut | SNP | A | A | G | COG-N-515-D | g.chr1:32557521A>G | ENST00000336294.5 | + | 6 | | | 982 | c.836A>G | c.(835-837)aAg>aGg | p.K279R |
| CCDC87 | 55231 | 11 | 66358100 | 66358100 | Missense Mut | SNP | T | T | A | COG-N-515-D | g.chr11:66358100T>A | ENST00000333861.3 | - | 1 | | | 2454 | c.2387A>T | c.(2386-2388)gAg>gTg | p.E796V |
| NDUFAF7 | 55471 | 2 | 37464921 | 37464921 | Missense Mut | SNP | A | A | C | COG-N-512-D | g.chr2:37464921A>C | ENST00000002125.4 | + | 4 | | | 359 | c.319A>C | c.(319-321)Agt>Cgt | p.S107R |
| NDUFAF7 | 55471 | 2 | 37464921 | 37464921 | Missense Mut | SNP | A | A | C | COG-N-515-D | g.chr2:37464921A>C | ENST00000002125.4 | + | 4 | | | 359 | c.319A>C | c.(319-321)Agt>Cgt | p.S107R |
| ASAP3 | 55616 | 1 | 23763148 | 23763148 | Splice Site | SNP | C | C | T | COG-N-512-D | g.chr1:23763148C>T | ENST00000336689.3 | - | 16 | | | 1521 | | c.e16-1 | |
| CARKD | 55739 | 13 | 111290730 | 111290730 | Missense Mut | SNP | G | G | C | COG-N-512-D | g.chr13:111290730G>C | ENST00000309957.2 | + | 10 | | | 1049 | c.1035G>C | c.(1033-1035)caG>caC | p.Q345H |
| CARKD | 55739 | 13 | 111290730 | 111290730 | Missense Mut | SNP | G | G | C | COG-N-515-D | g.chr13:111290730G>C | ENST00000309957.2 | + | 10 | | | 1049 | c.1035G>C | c.(1033-1035)caG>caC | p.Q345H |
| EXOC1 | 55763 | 4 | 56737334 | 56737334 | Missense Mut | SNP | G | G | T | COG-N-512-D | g.chr4:56737334G>T | ENST00000381295.2 | + | 7 | | | 1247 | c.899G>T | c.(898-900)gGc>gTc | p.G300V |
| EXOC1 | 55763 | 4 | 56737334 | 56737334 | Missense Mut | SNP | G | G | T | COG-N-515-D | g.chr4:56737334G>T | ENST00000381295.2 | + | 7 | | | 1247 | c.899G>T | c.(898-900)gGc>gTc | p.G300V |
| GPRC5C | 55890 | 17 | 72436078 | 72436078 | Missense Mut | SNP | G | G | A | COG-N-512-D | g.chr17:72436078G>A | ENST00000481232.1 | + | 2 | | | 809 | c.298G>A | c.(298-300)Gcc>Acc | p.A100T |
| GPRC5C | 55890 | 17 | 72436078 | 72436078 | Missense Mut | SNP | G | G | A | COG-N-515-D | g.chr17:72436078G>A | ENST00000481232.1 | + | 2 | | | 809 | c.298G>A | c.(298-300)Gcc>Acc | p.A100T |
| PCDHGA1 | 56114 | 5 | 140710411 | 140710411 | Missense Mut | SNP | C | C | A | COG-N-512-D | g.chr5:140710411C>A | ENST00000517417.1 | + | 1 | | | 160 | c.160C>A | c.(160-162)Ctg>Atg | p.L54M |
| PCDHGA1 | 56114 | 5 | 140710411 | 140710411 | Missense Mut | SNP | C | C | A | COG-N-515-D | g.chr5:140710411C>A | ENST00000517417.1 | + | 1 | | | 160 | c.160C>A | c.(160-162)Ctg>Atg | p.L54M |
| PCDHA1 | 56147 | 5 | 140166287 | 140166287 | Missense Mut | SNP | C | C | A | COG-N-512-D | g.chr5:140166287C>A | ENST00000504120.2 | + | 1 | | | 412 | c.412C>A | c.(412-414)Caa>Aaa | p.Q138K |
| PARD3 | 56288 | 10 | 34420464 | 34420464 | Missense Mut | SNP | T | T | G | COG-N-512-D | g.chr10:34420464T>G | ENST00000374789.3 | - | 23 | | | 3801 | c.3476A>C | c.(3475-3477)cAa>cCa | p.Q1159P |
| PARD3 | 56288 | 10 | 34420464 | 34420464 | Missense Mut | SNP | T | T | G | COG-N-515-D | g.chr10:34420464T>G | ENST00000374789.3 | - | 23 | | | 3801 | c.3476A>C | c.(3475-3477)cAa>cCa | p.Q1159P |
| NMUR2 | 56923 | 5 | 151784035 | 151784035 | Missense Mut | SNP | T | T | C | COG-N-515-D | g.chr5:151784035T>C | ENST00000255262.3 | - | 1 | | | 805 | c.640A>G | c.(640-642)Aat>Gat | p.N214D |
| CASC5 | 57082 | 15 | 40912899 | 40912899 | Nonsense Mut | SNP | C | C | A | COG-N-515-D | g.chr15:40912899C>A | ENST00000346991.5 | + | 11 | | | 905 | c.515C>A | c.(514-516)tCa>tAa | p.S172* |
| RGL3 | 57139 | 19 | 11505163 | 11505163 | Missense Mut | SNP | T | T | C | COG-N-512-D | g.chr19:11505163T>C | ENST00000380456.3 | - | 19 | | | 2111 | c.2048A>G | c.(2047-2049)tAt>tGt | p.Y683C |
| RGL3 | 57139 | 19 | 11505163 | 11505163 | Missense Mut | SNP | T | T | C | COG-N-515-D | g.chr19:11505163T>C | ENST00000380456.3 | - | 19 | | | 2111 | c.2048A>G | c.(2047-2049)tAt>tGt | p.Y683C |
| KIF17 | 57576 | 1 | 21044046 | 21044046 | Missense Mut | SNP | A | A | G | COG-N-512-D | g.chr1:21044046A>G | ENST00000247986.2 | - | 1 | | | 464 | c.154T>C | c.(154-156)Ttc>Ctc | p.F52L |
| KIF17 | 57576 | 1 | 21044046 | 21044046 | Missense Mut | SNP | A | A | G | COG-N-515-D | g.chr1:21044046A>G | ENST00000247986.2 | - | 1 | | | 464 | c.154T>C | c.(154-156)Ttc>Ctc | p.F52L |
| EBF4 | 57593 | 20 | 2736431 | 2736431 | Missense Mut | SNP | C | C | A | COG-N-512-D | g.chr20:2736431C>A | ENST00000609451.1 | + | 15 | | | 1771 | c.1699C>A | c.(1699-1701)Ccc>Acc | p.P567T |
| EBF4 | 57593 | 20 | 2736431 | 2736431 | Missense Mut | SNP | C | C | A | COG-N-515-D | g.chr20:2736431C>A | ENST00000609451.1 | + | 15 | | | 1771 | c.1699C>A | c.(1699-1701)Ccc>Acc | p.P567T |
| PITPNM2 | 57605 | 12 | 123470680 | 123470680 | Missense Mut | SNP | G | G | A | COG-N-515-D | g.chr12:123470680G>A | ENST00000542749.1 | - | 24 | | | 4007 | c.3944C>T | c.(3943-3945)gCg>gTg | p.A1315V |
| EP400 | 57634 | 12 | 132514316 | 132514321 | In Frame Del | DEL | TACAGC | TACAGC | - | COG-N-512-D | g.chr12:132514316 132514 | ENST00000389561.2 | + | 28 | 5561 | 5566 | | c.5452 5457del | c.(5452-5457)tacagcdel | p.YS1818del |
| CAMSAP3 | 57662 | 19 | 7676467 | 7676467 | Missense Mut | SNP | G | G | T | COG-N-512-D | g.chr19:7676467G>T | ENST00000446248.2 | + | 12 | | | 1370 | c.1269G>T | c.(1267-1269)caG>caT | p.Q423H |
| CAMSAP3 | 57662 | 19 | 7676467 | 7676467 | Missense Mut | SNP | G | G | T | COG-N-515-D | g.chr19:7676467G>T | ENST00000446248.2 | + | 12 | | | 1370 | c.1269G>T | c.(1267-1269)caG>caT | p.Q423H |
| ZBTB26 | 57684 | 9 | 125681228 | 125681228 | Missense Mut | SNP | T | T | C | COG-N-512-D | g.chr9:125681228T>C | ENST00000373656.3 | - | 2 | | | 1059 | c.986A>G | c.(985-987)aAa>aGa | p.K329R |
| ZBTB26 | 57684 | 9 | 125681228 | 125681228 | Missense Mut | SNP | T | T | C | COG-N-515-D | g.chr9:125681228T>C | ENST00000373656.3 | - | 2 | | | 1059 | c.986A>G | c.(985-987)aAa>aGa | p.K329R |
| RXFP1 | 59350 | 4 | 159493919 | 159493919 | Missense Mut | SNP | G | G | A | COG-N-512-D | g.chr4:159493919G>A | ENST00000470033.1 | + | 2 | | | 119 | c.119G>A | c.(118-120)tGc>tAc | p.C40Y |
| RXFP1 | 59350 | 4 | 159493919 | 159493919 | Missense Mut | SNP | G | G | A | COG-N-515-D | g.chr4:159493919G>A | ENST00000470033.1 | + | 2 | | | 119 | c.119G>A | c.(118-120)tGc>tAc | p.C40Y |
| FAM111A | 63901 | 11 | 58920151 | 58920151 | Missense Mut | SNP | T | T | C | COG-N-512-D | g.chr11:58920151T>C | ENST00000528737.1 | + | 5 | | | 3828 | c.1010T>C | c.(1009-1011)gTa>gCa | p.V337A |
| FAM111A | 63901 | 11 | 58920151 | 58920151 | Missense Mut | SNP | T | T | C | COG-N-515-D | g.chr11:58920151T>C | ENST00000528737.1 | + | 5 | | | 3828 | c.1010T>C | c.(1009-1011)gTa>gCa | p.V337A |
| NAPB | 63908 | 20 | 23358077 | 23358077 | Missense Mut | SNP | G | G | T | COG-N-512-D | g.chr20:23358077G>T | ENST00000377026.4 | - | 11 | | | 918 | c.833C>A | c.(832-834)aCc>aAc | p.T278N |
| NAPB | 63908 | 20 | 23358077 | 23358077 | Missense Mut | SNP | G | G | T | COG-N-515-D | g.chr20:23358077G>T | ENST00000377026.4 | - | 11 | | | 918 | c.833C>A | c.(832-834)aCc>aAc | p.T278N |
| KIF13A | 63971 | 6 | 17764972 | 17764972 | Missense Mut | SNP | G | G | A | COG-N-515-D | g.chr6:17764972G>A | ENST00000259711.6 | - | 39 | | | 4892 | c.4787C>T | c.(4786-4788)aCc>aTc | p.T1596I |
| PRDM15 | 63977 | 21 | 43299383 | 43299383 | Missense Mut | SNP | C | C | T | COG-N-512-D | g.chr21:43299383C>T | ENST00000398548.1 | - | 1 | | | 199 | c.98G>A | c.(97-99)aGg>aAg | p.R33K |
| PRDM15 | 63977 | 21 | 43299383 | 43299383 | Missense Mut | SNP | C | C | T | COG-N-515-D | g.chr21:43299383C>T | ENST00000398548.1 | - | 1 | | | 199 | c.98G>A | c.(97-99)aGg>aAg | p.R33K |
| MMP27 | 64066 | 11 | 102567521 | 102567521 | Missense Mut | SNP | A | A | T | COG-N-512-D | g.chr11:102567521A>T | ENST00000260229.4 | - | 5 | | | 756 | c.665T>A | c.(664-666)cTg>cAg | p.L222Q |
| MMP27 | 64066 | 11 | 102567521 | 102567521 | Missense Mut | SNP | A | A | T | COG-N-515-D | g.chr11:102567521A>T | ENST00000260229.4 | - | 5 | | | 756 | c.665T>A | c.(664-666)cTg>cAg | p.L222Q |
| S100PBP | 64766 | 1 | 33293660 | 33293660 | Missense Mut | SNP | C | C | T | COG-N-512-D | g.chr1:33293660C>T | ENST00000373475.5 | + | 4 | | | 1152 | c.898C>T | c.(898-900)Cct>Tct | p.P300S |
| S100PBP | 64766 | 1 | 33293660 | 33293660 | Missense Mut | SNP | C | C | T | COG-N-515-D | g.chr1:33293660C>T | ENST00000373475.5 | + | 4 | | | 1152 | c.898C>T | c.(898-900)Cct>Tct | p.P300S |
| LPIN3 | 64900 | 20 | 39977300 | 39977300 | Missense Mut | SNP | G | G | C | COG-N-512-D | g.chr20:39977300G>C | ENST00000373257.3 | + | 4 | | | 421 | c.330G>C | c.(328-330)tgG>tgC | p.W110C |
| LPIN3 | 64900 | 20 | 39977300 | 39977300 | Missense Mut | SNP | G | G | C | COG-N-515-D | g.chr20:39977300G>C | ENST00000373257.3 | + | 4 | | | 421 | c.330G>C | c.(328-330)tgG>tgC | p.W110C |
| NBEAL1 | 65065 | 2 | 204073908 | 204073908 | Missense Mut | SNP | A | A | G | COG-N-512-D | g.chr2:204073908A>G | ENST00000449802.1 | + | 52 | | | 7894 | c.7561A>G | c.(7561-7563)Att>Gtt | p.I2521V |
| NBEAL1 | 65065 | 2 | 204073908 | 204073908 | Missense Mut | SNP | A | A | G | COG-N-515-D | g.chr2:204073908A>G | ENST00000449802.1 | + | 52 | | | 7894 | c.7561A>G | c.(7561-7563)Att>Gtt | p.I2521V |
| PHACTR4 | 65979 | 1 | 28792238 | 28792238 | Missense Mut | SNP | A | A | G | COG-N-512-D | g.chr1:28792238A>G | ENST00000373836.3 | + | 4 | | | 558 | c.344A>G | c.(343-345)aAt>aGt | p.N115S |
| PHACTR4 | 65979 | 1 | 28792238 | 28792238 | Missense Mut | SNP | A | A | G | COG-N-515-D | g.chr1:28792238A>G | ENST00000373836.3 | + | 4 | | | 558 | c.344A>G | c.(343-345)aAt>aGt | p.N115S |
| DLK2 | 65989 | 6 | 43420863 | 43420863 | Missense Mut | SNP | C | C | G | COG-N-512-D | g.chr6:43420863C>G | ENST00000357338.3 | - | 4 | | | 851 | c.151G>C | c.(151-153)Ggc>Cgc | p.G51R |
| PALB2 | 79728 | 16 | 23614814 | 23614814 | Missense Mut | SNP | T | T | A | COG-N-515-D | g.chr16:23614814T>A | ENST00000261584.4 | - | 13 | | | 3679 | c.3527A>T | c.(3526-3528)aAa>aTa | p.K1176I |
| SLC25A22 | 79751 | 11 | 794852 | 794852 | Missense Mut | SNP | T | T | A | COG-N-515-D | g.chr11:794852T>A | ENST00000320230.5 | - | 3 | | | 551 | c.70A>T | c.(70-72)Acc>Tcc | p.T24S |
| ZFHX4 | 79776 | 8 | 77763172 | 77763172 | Missense Mut | SNP | G | G | T | COG-N-515-D | g.chr8:77763172G>T | ENST00000521891.2 | + | 10 | | | 4463 | c.4015G>T | c.(4015-4017)Gct>Tct | p.A1339S |
| CPED1 | 79974 | 7 | 120776144 | 120776144 | Splice Site | SNP | G | G | T | COG-N-512-D | g.chr7:120776144G>T | ENST00000310396.5 | + | 14 | | | 2166 | c.1699G>T | c.(1699-1701)Gat>Tat | p.D567Y |
| CPED1 | 79974 | 7 | 120776144 | 120776144 | Splice Site | SNP | G | G | T | COG-N-515-D | g.chr7:120776144G>T | ENST00000310396.5 | + | 14 | | | 2166 | c.1699G>T | c.(1699-1701)Gat>Tat | p.D567Y |
| GREB1L | 80000 | 18 | 19079882 | 19079882 | Missense Mut | SNP | G | G | A | COG-N-512-D | g.chr18:19079882G>A | ENST00000580732.2 | + | 22 | | | 3965 | c.3584G>A | c.(3583-3585)aGc>aAc | p.S1195N |
| GREB1L | 80000 | 18 | 19079882 | 19079882 | Missense Mut | SNP | G | G | A | COG-N-515-D | g.chr18:19079882G>A | ENST00000580732.2 | + | 22 | | | 3965 | c.3584G>A | c.(3583-3585)aGc>aAc | p.S1195N |
| ADAMTS20 | 80070 | 12 | 43770437 | 43770437 | Missense Mut | SNP | A | A | G | COG-N-512-D | g.chr12:43770437A>G | ENST00000389420.3 | - | 33 | | | 5014 | c.5015T>C | c.(5014-5016)aTt>aCt | p.I1672T |
| ADAMTS20 | 80070 | 12 | 43770437 | 43770437 | Missense Mut | SNP | A | A | G | COG-N-515-D | g.chr12:43770437A>G | ENST00000389420.3 | - | 33 | | | 5014 | c.5015T>C | c.(5014-5016)aTt>aCt | p.I1672T |
| BPIFB2 | 80341 | 20 | 31608200 | 31608200 | Splice Site | SNP | G | G | C | COG-N-512-D | g.chr20:31608200G>C | ENST00000170150.3 | + | 12 | | | 1342 | c.1147G>C | c.(1147-1149)Ggg>Cgg | p.G383R |
| ZFP91 | 80829 | 11 | 58377494 | 58377494 | Missense Mut | SNP | A | A | T | COG-N-515-D | g.chr11:58377494A>T | ENST00000316059.6 | + | 3 | | | 733 | c.562A>T | c.(562-564)Aca>Tca | p.T188S |
| SYNC | 81493 | 1 | 33160492 | 33160492 | Nonsense Mut | SNP | G | G | A | COG-N-512-D | g.chr1:33160492G>A | ENST00000409190.3 | - | 2 | | | 1665 | c.1207C>T | c.(1207-1209)Cga>Tga | p.R403* |
| SYNC | 81493 | 1 | 33160492 | 33160492 | Nonsense Mut | SNP | G | G | A | COG-N-515-D | g.chr1:33160492G>A | ENST00000409190.3 | - | 2 | | | 1665 | c.1207C>T | c.(1207-1209)Cga>Tga | p.R403* |
| PVRL4 | 81607 | 1 | 161043547 | 161043547 | Missense Mut | SNP | C | C | A | COG-N-512-D | g.chr1:161043547C>A | ENST00000368012.3 | - | 7 | | | 1498 | c.1196G>T | c.(1195-1197)cGg>cTg | p.R399L |
| PVRL4 | 81607 | 1 | 161043547 | 161043547 | Missense Mut | SNP | C | C | A | COG-N-515-D | g.chr1:161043547C>A | ENST00000368012.3 | - | 7 | | | 1498 | c.1196G>T | c.(1195-1197)cGg>cTg | p.R399L |
| PMFBP1 | 83449 | 16 | 72170475 | 72170475 | Missense Mut | SNP | C | C | T | COG-N-512-D | g.chr16:72170475C>T | ENST00000237353.10 | - | 9 | | | 1336 | c.1075G>A | c.(1075-1077)Gga>Aga | p.G359R |
| PMFBP1 | 83449 | 16 | 72170475 | 72170475 | Missense Mut | SNP | C | C | T | COG-N-515-D | g.chr16:72170475C>T | ENST00000237353.10 | - | 9 | | | 1336 | c.1075G>A | c.(1075-1077)Gga>Aga | p.G359R |
| PARP9 | 83666 | 3 | 122259381 | 122259381 | Missense Mut | SNP | A | A | G | COG-N-512-D | g.chr3:122259381A>G | ENST00000360356.2 | - | 8 | | | 2035 | c.1808T>C | c.(1807-1809)aTg>aCg | p.M603T |
| PARP9 | 83666 | 3 | 122259381 | 122259381 | Missense Mut | SNP | A | A | G | COG-N-515-D | g.chr3:122259381A>G | ENST00000360356.2 | - | 8 | | | 2035 | c.1808T>C | c.(1807-1809)aTg>aCg | p.M603T |
| TM2D1 | 83941 | 1 | 62190659 | 62190659 | Missense Mut | SNP | G | G | A | COG-N-512-D | g.chr1:62190659G>A | ENST00000606498.1 | - | 1 | | | 154 | c.134C>T | c.(133-135)tCg>tTg | p.S45L |
| TM2D1 | 83941 | 1 | 62190659 | 62190659 | Missense Mut | SNP | G | G | A | COG-N-515-D | g.chr1:62190659G>A | ENST00000606498.1 | - | 1 | | | 154 | c.134C>T | c.(133-135)tCg>tTg | p.S45L |
| PLEKHN1 | 84069 | 1 | 906082 | 906082 | Missense Mut | SNP | C | C | T | COG-N-512-D | g.chr1:906082C>T | ENST00000379410.3 | + | 5 | | | 463 | c.428C>T | c.(427-429)aCg>aTg | p.T143M |
| PLEKHN1 | 84069 | 1 | 906082 | 906082 | Missense Mut | SNP | C | C | T | COG-N-515-D | g.chr1:906082C>T | ENST00000379410.3 | + | 5 | | | 463 | c.428C>T | c.(427-429)aCg>aTg | p.T143M |
| FAM186B | 84070 | 12 | 49997138 | 49997138 | Missense Mut | SNP | G | G | T | COG-N-512-D | g.chr12:49997138G>T | ENST00000257894.2 | - | 3 | | | 496 | c.335C>A | c.(334-336)aCc>aAc | p.T112N |
| LRRIQ1 | 84125 | 12 | 85441180 | 85441180 | Nonsense Mut | SNP | G | G | T | COG-N-512-D | g.chr12:85441180G>T | ENST00000393217.2 | + | 6 | | | 671 | c.610G>T | c.(610-612)Gaa>Taa | p.E204* |
| KIAA1109 | 84162 | 4 | 123167859 | 123167859 | Missense Mut | SNP | G | G | A | COG-N-512-D | g.chr4:123167859G>A | ENST00000264501.4 | + | 34 | | | 5579 | c.5206G>A | c.(5206-5208)Gcc>Acc | p.A1736T |
| KIAA1109 | 84162 | 4 | 123167859 | 123167859 | Missense Mut | SNP | G | G | A | COG-N-515-D | g.chr4:123167859G>A | ENST00000264501.4 | + | 34 | | | 5579 | c.5206G>A | c.(5206-5208)Gcc>Acc | p.A1736T |
| NOA1 | 84273 | 4 | 57829741 | 57829741 | Missense Mut | SNP | G | G | A | COG-N-512-D | g.chr4:57829741G>A | ENST00000264230.4 | - | 7 | | | 3209 | c.1972C>T | c.(1972-1974)Cct>Tct | p.P658S |
| NOA1 | 84273 | 4 | 57829741 | 57829741 | Missense Mut | SNP | G | G | A | COG-N-515-D | g.chr4:57829741G>A | ENST00000264230.4 | - | 7 | | | 3209 | c.1972C>T | c.(1972-1974)Cct>Tct | p.P658S |
| ALG10 | 84920 | 12 | 34179436 | 34179436 | Missense Mut | SNP | A | A | C | COG-N-512-D | g.chr12:34179436A>C | ENST00000266483.2 | + | 3 | | | 1327 | c.1008A>C | c.(1006-1008)ttA>ttC | p.L336F |
| ALG10 | 84920 | 12 | 34179436 | 34179436 | Missense Mut | SNP | A | A | C | COG-N-515-D | g.chr12:34179436A>C | ENST00000266483.2 | + | 3 | | | 1327 | c.1008A>C | c.(1006-1008)ttA>ttC | p.L336F |
| KNDC1 | 85442 | 10 | 134980883 | 134980883 | Splice Site | SNP | A | A | T | COG-N-512-D | g.chr10:134980883A>T | ENST00000304613.3 | + | 2 | | | 123 | | c.e2-1 | |
| KNDC1 | 85442 | 10 | 134980883 | 134980883 | Splice Site | SNP | A | A | T | COG-N-515-D | g.chr10:134980883A>T | ENST00000304613.3 | + | 2 | | | 123 | | c.e2-1 | |
| GAL3ST3 | 89792 | 11 | 65811122 | 65811122 | Missense Mut | SNP | C | C | A | COG-N-512-D | g.chr11:65811122C>A | ENST00000312006.4 | - | 3 | | | 433 | c.152G>T | c.(151-153)tGc>tTc | p.C51F |
| GAL3ST3 | 89792 | 11 | 65811122 | 65811122 | Missense Mut | SNP | C | C | A | COG-N-515-D | g.chr11:65811122C>A | ENST00000312006.4 | - | 3 | | | 433 | c.152G>T | c.(151-153)tGc>tTc | p.C51F |
| ZNF697 | 90874 | 1 | 120168566 | 120168566 | Missense Mut | SNP | G | G | A | COG-N-512-D | g.chr1:120168566G>A | ENST00000421812.2 | - | 2 | | | 277 | c.158C>T | c.(157-159)cCg>cTg | p.P53L |
| ZNF697 | 90874 | 1 | 120168566 | 120168566 | Missense Mut | SNP | G | G | A | COG-N-515-D | g.chr1:120168566G>A | ENST00000421812.2 | - | 2 | | | 277 | c.158C>T | c.(157-159)cCg>cTg | p.P53L |
| POM121L2 | 94026 | 6 | 27278395 | 27278395 | Missense Mut | SNP | G | G | T | COG-N-512-D | g.chr6:27278395G>T | ENST00000444565.1 | - | 1 | | | 1554 | c.1555C>A | c.(1555-1557)Ctt>Att | p.L519I |
| POM121L2 | 94026 | 6 | 27278395 | 27278395 | Missense Mut | SNP | G | G | T | COG-N-515-D | g.chr6:27278395G>T | ENST00000444565.1 | - | 1 | | | 1554 | c.1555C>A | c.(1555-1557)Ctt>Att | p.L519I |
| PGLYRP3 | 114771 | 1 | 153276484 | 153276484 | Splice Site | SNP | G | G | T | COG-N-512-D | g.chr1:153276484G>T | ENST00000290722.1 | - | 4 | | | 430 | c.378C>A | c.(376-378)ggC>ggA | p.G126G |
| PGLYRP3 | 114771 | 1 | 153276484 | 153276484 | Splice Site | SNP | G | G | T | COG-N-515-D | g.chr1:153276484G>T | ENST00000290722.1 | - | 4 | | | 430 | c.378C>A | c.(376-378)ggC>ggA | p.G126G |
| C1QTNF2 | 114898 | 5 | 159776486 | 159776486 | Missense Mut | SNP | A | A | T | COG-N-512-D | g.chr5:159776486A>T | ENST00000393975.3 | - | 3 | | | 685 | c.682T>A | c.(682-684)Tcc>Acc | p.S228T |

| | | | | | | | | | | | | | | | | | | | | | |
|---|---|---|---|---|---|---|---|---|---|---|---|---|---|---|---|---|---|---|---|---|---|
| C1QTNF2 | 114898 | | 5 | 159776489 | 159776489 | Missense Mut | SNP | C | C | T | COG-N-512-D | g.chr5:159776489C>T | ENST00000393975.3 | - | 3 | | | 682 | c.679G>A | c.(679-681)Gct>Act | p.A227T |
| C1QTNF2 | 114898 | | 5 | 159776489 | 159776489 | Missense Mut | SNP | C | C | T | COG-N-515-D | g.chr5:159776489C>T | ENST00000393975.3 | - | 3 | | | 682 | c.679G>A | c.(679-681)Gct>Act | p.A227T |
| C1QTNF2 | 114898 | | 5 | 159776486 | 159776486 | Missense Mut | SNP | A | A | T | COG-N-515-D | g.chr5:159776486A>T | ENST00000393975.3 | - | 3 | | | 685 | c.682T>A | c.(682-684)Tcc>Acc | p.S228T |
| NOSTRIN | 115677 | | 2 | 169707622 | 169707622 | Missense Mut | SNP | T | T | A | COG-N-512-D | g.chr2:169707622T>A | ENST00000458381.2 | + | 14 | | | 1588 | c.830T>A | c.(829-831)aTt>aAt | p.I277N |
| NOSTRIN | 115677 | | 2 | 169707622 | 169707622 | Missense Mut | SNP | T | T | A | COG-N-515-D | g.chr2:169707622T>A | ENST00000458381.2 | + | 14 | | | 1588 | c.830T>A | c.(829-831)aTt>aAt | p.I277N |
| CATSPER1 | 117144 | | 11 | 65790530 | 65790530 | Missense Mut | SNP | C | C | G | COG-N-512-D | g.chr11:65790530C>G | ENST00000312106.5 | - | 2 | | | 1356 | c.1219G>C | c.(1219-1221)Ggc>Cgc | p.G407R |
| CATSPER1 | 117144 | | 11 | 65790530 | 65790530 | Missense Mut | SNP | C | C | G | COG-N-515-D | g.chr11:65790530C>G | ENST00000312106.5 | - | 2 | | | 1356 | c.1219G>C | c.(1219-1221)Ggc>Cgc | p.G407R |
| CPXM2 | 119587 | | 10 | 125528089 | 125528089 | Missense Mut | SNP | C | C | T | COG-N-512-D | g.chr10:125528089C>T | ENST00000241305.3 | - | 9 | | | 1406 | c.1252G>A | c.(1252-1254)Gtc>Atc | p.V418I |
| CPXM2 | 119587 | | 10 | 125528089 | 125528089 | Missense Mut | SNP | C | C | T | COG-N-515-D | g.chr10:125528089C>T | ENST00000241305.3 | - | 9 | | | 1406 | c.1252G>A | c.(1252-1254)Gtc>Atc | p.V418I |
| PLD4 | 122618 | | 14 | 105398403 | 105398404 | Frame Shift De | DEL | GC | GC | - | COG-N-512-D | g.chr14:105398403_105398 | ENST00000392593.4 | + | 9 | 1281 | 1282 | | c.1113_1114del( | c.(1111-1116)gtgcgcfs | p.R372fs |
| PLD4 | 122618 | | 14 | 105398403 | 105398404 | Frame Shift De | DEL | GC | GC | - | COG-N-515-D | g.chr14:105398403_105398 | ENST00000392593.4 | + | 9 | 1281 | 1282 | | c.1113_1114del( | c.(1111-1116)gtgcgcfs | p.R372fs |
| SOCS4 | 122809 | | 14 | 55510127 | 55510127 | Missense Mut | SNP | A | A | G | COG-N-512-D | g.chr14:55510127A>G | ENST00000395472.2 | + | 2 | | | 700 | c.368A>G | c.(367-369)cAt>cGt | p.H123R |
| SOCS4 | 122809 | | 14 | 55510127 | 55510127 | Missense Mut | SNP | A | A | G | COG-N-515-D | g.chr14:55510127A>G | ENST00000395472.2 | + | 2 | | | 700 | c.368A>G | c.(367-369)cAt>cGt | p.H123R |
| USP43 | 124739 | | 17 | 9604481 | 9604481 | Missense Mut | SNP | T | T | G | COG-N-512-D | g.chr17:9604481T>G | ENST00000285199.7 | + | 11 | | | 1677 | c.1581T>G | c.(1579-1581)agT>agG | p.S527R |
| USP43 | 124739 | | 17 | 9604481 | 9604481 | Missense Mut | SNP | T | T | G | COG-N-515-D | g.chr17:9604481T>G | ENST00000285199.7 | + | 11 | | | 1677 | c.1581T>G | c.(1579-1581)agT>agG | p.S527R |
| TMEM132E | 124842 | | 17 | 32956132 | 32956132 | Missense Mut | SNP | G | G | A | COG-N-512-D | g.chr17:32956132G>A | ENST00000321639.5 | + | 5 | | | 1305 | c.977G>A | c.(976-978)cGt>cAt | p.R326H |
| TMEM132E | 124842 | | 17 | 32956132 | 32956132 | Missense Mut | SNP | G | G | A | COG-N-515-D | g.chr17:32956132G>A | ENST00000321639.5 | + | 5 | | | 1305 | c.977G>A | c.(976-978)cGt>cAt | p.R326H |
| TTC39C | 125488 | | 18 | 21649130 | 21649130 | Nonsense Mut | SNP | C | C | T | COG-N-512-D | g.chr18:21649130C>T | ENST00000317571.3 | + | 4 | | | 591 | c.355C>T | c.(355-357)Cga>Tga | p.R119* |
| TTC39C | 125488 | | 18 | 21649130 | 21649130 | Nonsense Mut | SNP | C | C | T | COG-N-515-D | g.chr18:21649130C>T | ENST00000317571.3 | + | 4 | | | 591 | c.355C>T | c.(355-357)Cga>Tga | p.R119* |
| ARID3C | 138715 | | 9 | 34625813 | 34625813 | Splice Site | SNP | T | T | G | COG-N-512-D | g.chr9:34625813T>G | ENST00000378909.2 | - | 2 | | | 411 | | c.e2-2 | |
| ARID3C | 138715 | | 9 | 34625813 | 34625813 | Splice Site | SNP | T | T | G | COG-N-515-D | g.chr9:34625813T>G | ENST00000378909.2 | - | 2 | | | 411 | | c.e2-2 | |
| DGKK | 139189 | X | | 50213388 | 50213423 | RNA | DEL | GGTTCTG( | GGTTCTGGGGCCGGTT | - | COG-N-512-D | g.chrX:50213388_50213423 | ENST00000376025.2 | - | 0 | 314 | 349 | | | | |
| ASB11 | 140456 | X | | 15306174 | 15306174 | Nonsense Mut | SNP | G | G | A | COG-N-512-D | g.chrX:15306174G>A | ENST00000480796.1 | - | 6 | | | 726 | c.676C>T | c.(676-678)Cag>Tag | p.Q226* |
| ASB11 | 140456 | X | | 15306174 | 15306174 | Nonsense Mut | SNP | G | G | A | COG-N-515-D | g.chrX:15306174G>A | ENST00000480796.1 | - | 6 | | | 726 | c.676C>T | c.(676-678)Cag>Tag | p.Q226* |
| SOGA1 | 140710 | | 20 | 35443779 | 35443779 | Missense Mut | SNP | C | C | T | COG-N-512-D | g.chr20:35443779C>T | ENST00000237536.4 | - | 5 | | | 2407 | c.2066G>A | c.(2065-2067)tGt>tAt | p.C689Y |
| SOGA1 | 140710 | | 20 | 35443779 | 35443779 | Missense Mut | SNP | C | C | T | COG-N-515-D | g.chr20:35443779C>T | ENST00000237536.4 | - | 5 | | | 2407 | c.2066G>A | c.(2065-2067)tGt>tAt | p.C689Y |
| FAM65C | 140876 | | 20 | 49225253 | 49225253 | Missense Mut | SNP | T | T | A | COG-N-512-D | g.chr20:49225253T>A | ENST00000327979.2 | - | 10 | | | 1106 | c.695A>T | c.(694-696)cAg>cTg | p.Q232L |
| FAM65C | 140876 | | 20 | 49225253 | 49225253 | Missense Mut | SNP | T | T | A | COG-N-515-D | g.chr20:49225253T>A | ENST00000327979.2 | - | 10 | | | 1106 | c.695A>T | c.(694-696)cAg>cTg | p.Q232L |
| CPNE8 | 144402 | | 12 | 39155977 | 39155977 | Missense Mut | SNP | A | A | T | COG-N-512-D | g.chr12:39155977A>T | ENST00000331366.5 | - | 9 | | | 713 | c.617T>A | c.(616-618)cTa>cAa | p.L206Q |
| CPNE8 | 144402 | | 12 | 39155977 | 39155977 | Missense Mut | SNP | A | A | T | COG-N-515-D | g.chr12:39155977A>T | ENST00000331366.5 | - | 9 | | | 713 | c.617T>A | c.(616-618)cTa>cAa | p.L206Q |
| KRT80 | 144501 | | 12 | 52585647 | 52585647 | Missense Mut | SNP | T | T | G | COG-N-512-D | g.chr12:52585647T>G | ENST00000394815.2 | - | 1 | | | 137 | c.40A>C | c.(40-42)Agc>Cgc | p.S14R |
| B3GALTL | 145173 | | 13 | 31860866 | 31860866 | Missense Mut | SNP | G | G | A | COG-N-512-D | g.chr13:31860866G>A | ENST00000343307.4 | + | 12 | | | 1123 | c.974G>A | c.(973-975)gGa>gAa | p.G325E |
| B3GALTL | 145173 | | 13 | 31860866 | 31860866 | Missense Mut | SNP | G | G | A | COG-N-515-D | g.chr13:31860866G>A | ENST00000343307.4 | + | 12 | | | 1123 | c.974G>A | c.(973-975)gGa>gAa | p.G325E |
| ZNF688 | 146542 | | 16 | 30581440 | 30581440 | Missense Mut | SNP | G | G | A | COG-N-515-D | g.chr16:30581440G>A | ENST00000223459.6 | - | 3 | | | 1732 | c.628C>T | c.(628-630)Cct>Tct | p.P210S |
| TMC8 | 147138 | | 17 | 76136988 | 76136988 | Missense Mut | SNP | C | C | T | COG-N-512-D | g.chr17:76136988C>T | ENST00000318430.5 | + | 16 | | | 2350 | c.1976C>T | c.(1975-1977)cCg>cTg | p.P659L |
| TMC8 | 147138 | | 17 | 76136988 | 76136988 | Missense Mut | SNP | C | C | T | COG-N-515-D | g.chr17:76136988C>T | ENST00000318430.5 | + | 16 | | | 2350 | c.1976C>T | c.(1975-1977)cCg>cTg | p.P659L |
| BTBD19 | 149478 | | 1 | 45279425 | 45279425 | Missense Mut | SNP | T | T | C | COG-N-512-D | g.chr1:45279425T>C | ENST00000409335.2 | + | 6 | | | 1041 | c.743T>C | c.(742-744)cTg>cCg | p.L248P |
| BTBD19 | 149478 | | 1 | 45279425 | 45279425 | Missense Mut | SNP | T | T | C | COG-N-515-D | g.chr1:45279425T>C | ENST00000409335.2 | + | 6 | | | 1041 | c.743T>C | c.(742-744)cTg>cCg | p.L248P |
| SLC9B1 | 150159 | | 4 | 103822372 | 103822372 | Nonsense Mut | SNP | C | C | A | COG-N-512-D | g.chr4:103822372C>A | ENST00000296422.7 | - | 12 | | | 1591 | c.1450G>T | c.(1450-1452)Gga>Tga | p.G484* |
| SLC9B1 | 150159 | | 4 | 103822372 | 103822372 | Nonsense Mut | SNP | C | C | A | COG-N-515-D | g.chr4:103822372C>A | ENST00000296422.7 | - | 12 | | | 1591 | c.1450G>T | c.(1450-1452)Gga>Tga | p.G484* |
| CCDC80 | 151887 | | 3 | 112337889 | 112337889 | Missense Mut | SNP | C | C | T | COG-N-512-D | g.chr3:112337889C>T | ENST00000206423.3 | - | 4 | | | 3051 | c.2098G>A | c.(2098-2100)Gag>Aag | p.E700K |
| CCDC80 | 151887 | | 3 | 112337889 | 112337889 | Missense Mut | SNP | C | C | T | COG-N-515-D | g.chr3:112337889C>T | ENST00000206423.3 | - | 4 | | | 3051 | c.2098G>A | c.(2098-2100)Gag>Aag | p.E700K |
| FAM53A | 152877 | | 4 | 1656812 | 1656812 | Missense Mut | SNP | G | G | T | COG-N-512-D | g.chr4:1656812G>T | ENST00000489363.1 | - | 4 | | | 1372 | c.775C>A | c.(775-777)Cgc>Agc | p.R259S |
| FAM53A | 152877 | | 4 | 1656812 | 1656812 | Missense Mut | SNP | G | G | T | COG-N-515-D | g.chr4:1656812G>T | ENST00000489363.1 | - | 4 | | | 1372 | c.775C>A | c.(775-777)Cgc>Agc | p.R259S |
| DAB2IP | 153090 | | 9 | 124526082 | 124526082 | Missense Mut | SNP | G | G | A | COG-N-512-D | g.chr9:124526082G>A | ENST00000309989.1 | + | 6 | | | 1159 | c.1012G>A | c.(1012-1014)Gct>Act | p.A338T |
| DAB2IP | 153090 | | 9 | 124526082 | 124526082 | Missense Mut | SNP | G | G | A | COG-N-515-D | g.chr9:124526082G>A | ENST00000309989.1 | + | 6 | | | 1159 | c.1012G>A | c.(1012-1014)Gct>Act | p.A338T |
| KCNU1 | 157855 | | 8 | 36790519 | 36790519 | Missense Mut | SNP | G | G | T | COG-N-512-D | g.chr8:36790519G>T | ENST00000399881.3 | + | 26 | | | 3050 | c.3013G>T | c.(3013-3015)Gat>Tat | p.D1005Y |
| KCNU1 | 157855 | | 8 | 36790519 | 36790519 | Missense Mut | SNP | G | G | T | COG-N-515-D | g.chr8:36790519G>T | ENST00000399881.3 | + | 26 | | | 3050 | c.3013G>T | c.(3013-3015)Gat>Tat | p.D1005Y |
| MDGA2 | 161357 | | 14 | 47687383 | 47687383 | Frame Shift De | DEL | T | T | - | COG-N-512-D | g.chr14:47687383delT | ENST00000399232.2 | - | 3 | | | 593 | c.229delA | c.(229-231)acafs | p.T77fs |
| MDGA2 | 161357 | | 14 | 47687383 | 47687383 | Frame Shift De | DEL | T | T | - | COG-N-515-D | g.chr14:47687383delT | ENST00000399232.2 | - | 3 | | | 593 | c.229delA | c.(229-231)acafs | p.T77fs |
| WDR90 | 197335 | | 16 | 711635 | 711635 | Missense Mut | SNP | G | G | A | COG-N-512-D | g.chr16:711635G>A | ENST00000293879.4 | + | 31 | | | 3712 | c.3712G>A | c.(3712-3714)Gcc>Acc | p.A1238T |
| UNC5B | 219699 | | 10 | 73056469 | 73056469 | Missense Mut | SNP | G | G | C | COG-N-515-D | g.chr10:73056469G>C | ENST00000335350.6 | + | 15 | | | 2876 | c.2460G>C | c.(2458-2460)caG>caC | p.Q820H |
| ANKK1 | 255239 | | 11 | 113270892 | 113270892 | Missense Mut | SNP | G | G | A | COG-N-512-D | g.chr11:113270892G>A | ENST00000303941.3 | + | 8 | | | 2295 | c.2201G>A | c.(2200-2202)cGc>cAc | p.R734H |
| ANKK1 | 255239 | | 11 | 113270892 | 113270892 | Missense Mut | SNP | G | G | A | COG-N-515-D | g.chr11:113270892G>A | ENST00000303941.3 | + | 8 | | | 2295 | c.2201G>A | c.(2200-2202)cGc>cAc | p.R734H |
| C8orf31 | 286122 | | 8 | 144124496 | 144124496 | Missense Mut | SNP | G | G | T | COG-N-515-D | g.chr8:144124496G>T | ENST00000395172.1 | + | 2 | | | 430 | c.78G>T | c.(76-78)agG>agT | p.R26S |
| LIN9 | 286826 | | 1 | 226421221 | 226421221 | Missense Mut | SNP | C | C | A | COG-N-512-D | g.chr1:226421221C>A | ENST00000328205.5 | - | 13 | | | 1842 | c.1297G>T | c.(1297-1299)Gct>Tct | p.A433S |
| LIN9 | 286826 | | 1 | 226421221 | 226421221 | Missense Mut | SNP | C | C | A | COG-N-515-D | g.chr1:226421221C>A | ENST00000328205.5 | - | 13 | | | 1842 | c.1297G>T | c.(1297-1299)Gct>Tct | p.A433S |
| RAB37 | 326624 | | 17 | 72733482 | 72733482 | Missense Mut | SNP | C | C | A | COG-N-515-D | g.chr17:72733482C>A | ENST00000392613.5 | + | 1 | | | 127 | c.71C>A | c.(70-72)cCg>cAg | p.P24Q |
| RNASE10 | 338879 | | 14 | 20979126 | 20979126 | Missense Mut | SNP | T | T | A | COG-N-512-D | g.chr14:20979126T>A | ENST00000328444.5 | + | 1 | | | 515 | c.496T>A | c.(496-498)Tgt>Agt | p.C166S |
| RNASE10 | 338879 | | 14 | 20979126 | 20979126 | Missense Mut | SNP | T | T | A | COG-N-515-D | g.chr14:20979126T>A | ENST00000328444.5 | + | 1 | | | 515 | c.496T>A | c.(496-498)Tgt>Agt | p.C166S |
| CCDC178 | 374864 | | 18 | 30926253 | 30926253 | Nonsense Mut | SNP | T | T | A | COG-N-515-D | g.chr18:30926253T>A | ENST00000383096.3 | - | 9 | | | 762 | c.580A>T | c.(580-582)Aga>Tga | p.R194* |
| CEACAM16 | 388551 | | 19 | 45206671 | 45206671 | Missense Mut | SNP | G | G | T | COG-N-512-D | g.chr19:45206671G>T | ENST00000405314.2 | + | 2 | | | 187 | c.90G>T | c.(88-90)caG>caT | p.Q30H |
| CEACAM16 | 388551 | | 19 | 45206671 | 45206671 | Missense Mut | SNP | G | G | T | COG-N-515-D | g.chr19:45206671G>T | ENST00000405314.2 | + | 2 | | | 187 | c.90G>T | c.(88-90)caG>caT | p.Q30H |
| CD164L2 | 388611 | | 1 | 27709666 | 27709666 | Missense Mut | SNP | G | G | C | COG-N-512-D | g.chr1:27709666G>C | ENST00000374027.3 | - | 1 | | | 127 | c.64C>G | c.(64-66)Cta>Gta | p.L22V |
| CD164L2 | 388611 | | 1 | 27709666 | 27709666 | Missense Mut | SNP | G | G | C | COG-N-515-D | g.chr1:27709666G>C | ENST00000374027.3 | - | 1 | | | 127 | c.64C>G | c.(64-66)Cta>Gta | p.L22V |
| OR51I1 | 390063 | | 11 | 5462205 | 5462205 | Nonsense Mut | SNP | G | G | T | COG-N-512-D | g.chr11:5462205G>T | ENST00000380211.1 | - | 1 | | | 539 | c.540C>A | c.(538-540)taC>taA | p.Y180* |
| OR51I1 | 390063 | | 11 | 5462213 | 5462213 | Missense Mut | SNP | G | G | A | COG-N-512-D | g.chr11:5462213G>A | ENST00000380211.1 | - | 1 | | | 531 | c.532C>T | c.(532-534)Cac>Tac | p.H178Y |
| OR51I1 | 390063 | | 11 | 5462205 | 5462205 | Nonsense Mut | SNP | G | G | T | COG-N-515-D | g.chr11:5462205G>T | ENST00000380211.1 | - | 1 | | | 539 | c.540C>A | c.(538-540)taC>taA | p.Y180* |
| OR51I1 | 390063 | | 11 | 5462213 | 5462213 | Missense Mut | SNP | G | G | A | COG-N-515-D | g.chr11:5462213G>A | ENST00000380211.1 | - | 1 | | | 531 | c.532C>T | c.(532-534)Cac>Tac | p.H178Y |
| C4orf3 | 401152 | | 4 | 120221566 | 120221566 | Missense Mut | SNP | T | T | G | COG-N-512-D | g.chr4:120221566T>G | ENST00000504110.1 | - | 1 | | | 510 | c.125A>C | c.(124-126)cAc>cCc | p.H42P |
| C4orf3 | 401152 | | 4 | 120221566 | 120221566 | Missense Mut | SNP | T | T | G | COG-N-515-D | g.chr4:120221566T>G | ENST00000504110.1 | - | 1 | | | 510 | c.125A>C | c.(124-126)cAc>cCc | p.H42P |
| IFIT1B | 439996 | | 10 | 91143855 | 91143855 | Missense Mut | SNP | G | G | T | COG-N-512-D | g.chr10:91143855G>T | ENST00000371809.3 | + | 2 | | | 865 | c.785G>T | c.(784-786)cGa>cTa | p.R262L |
| IFIT1B | 439996 | | 10 | 91143855 | 91143855 | Missense Mut | SNP | G | G | T | COG-N-515-D | g.chr10:91143855G>T | ENST00000371809.3 | + | 2 | | | 865 | c.785G>T | c.(784-786)cGa>cTa | p.R262L |
| CBLN3 | 643866 | | 14 | 24897000 | 24897000 | Missense Mut | SNP | G | G | C | COG-N-512-D | g.chr14:24897000G>C | ENST00000555436.1 | - | 3 | | | 528 | c.460C>G | c.(460-462)Ctc>Gtc | p.L154V |
| CBLN3 | 643866 | | 14 | 24897000 | 24897000 | Missense Mut | SNP | G | G | C | COG-N-515-D | g.chr14:24897000G>C | ENST00000555436.1 | - | 3 | | | 528 | c.460C>G | c.(460-462)Ctc>Gtc | p.L154V |
| FAM83G | 644815 | | 17 | 18880988 | 18880988 | Missense Mut | SNP | C | C | T | COG-N-512-D | g.chr17:18880988C>T | ENST00000388995.6 | - | 5 | | | 2214 | c.1991G>A | c.(1990-1992)gGt>gAt | p.G664D |
| FAM83G | 644815 | | 17 | 18880988 | 18880988 | Missense Mut | SNP | C | C | T | COG-N-515-D | g.chr17:18880988C>T | ENST00000388995.6 | - | 5 | | | 2214 | c.1991G>A | c.(1990-1992)gGt>gAt | p.G664D |

#### Table S3

**GSEA results for BIOCARTA gene sets comparing ALT vs telomerase positive non-MYCN amplified cell lines**

| NAME | NES | NOM p-val | FDR q-val | logP(+-) |
|---|---|---|---|---|
| BIOCARTA_ATRBRCA_PATHWAY | 2.1289535 | 3.93E-04 | 0.004126982 | 3.41 |
| BIOCARTA_ATM_PATHWAY | 1.8838812 | 0.00109172 | 0.03761379 | 2.96 |
| BIOCARTA_MCM_PATHWAY | 1.8753248 | 0.00127032 | 0.027966915 | 2.90 |
| BIOCARTA_EFP_PATHWAY | 1.8419476 | 0.00183936 | 0.03204158 | 2.74 |
| BIOCARTA_TID_PATHWAY | 1.7171627 | 0.00723666 | 0.10407296 | 2.14 |
| BIOCARTA_CASPASE_PATHWAY | 1.7124015 | 0.0067613 | 0.09083461 | 2.17 |
| BIOCARTA_TNFR2_PATHWAY | 1.6792283 | 0.01093231 | 0.10671545 | 1.96 |
| BIOCARTA_G2_PATHWAY | 1.6025441 | 0.01744426 | 0.18314111 | 1.76 |
| BIOCARTA_FAS_PATHWAY | 1.6010855 | 0.01307202 | 0.16493185 | 1.88 |
| BIOCARTA_STRESS_PATHWAY | 1.5994318 | 0.01782958 | 0.1504423 | 1.75 |
| BIOCARTA_NTHI_PATHWAY | 1.5789211 | 0.01978796 | 0.1613891 | 1.70 |
| BIOCARTA_TEL_PATHWAY | 1.544748 | 0.03210611 | 0.1931536 | 1.49 |
| BIOCARTA_CELLCYCLE_PATHWAY | 1.5426427 | 0.02840808 | 0.18106544 | 1.55 |
| BIOCARTA_VDR_PATHWAY | 1.5285511 | 0.02820927 | 0.18637303 | 1.55 |
| BIOCARTA_G1_PATHWAY | 1.5183979 | 0.02735938 | 0.18724297 | 1.56 |
| BIOCARTA_HIVNEF_PATHWAY | 1.5058615 | 0.01457663 | 0.19186962 | 1.84 |
| BIOCARTA_RACCYCD_PATHWAY | 1.4652721 | 0.04466555 | 0.23800822 | 1.35 |
| BIOCARTA_MITOCHONDRIA_PATHWAY | 1.421541 | 0.06839762 | 0.29755273 | 1.16 |
| BIOCARTA_41BB_PATHWAY | 1.4149821 | 0.0731664 | 0.29365885 | 1.14 |
| BIOCARTA_TOLL_PATHWAY | 1.3481433 | 0.09532993 | 0.41185823 | 1.02 |
| BIOCARTA_ECM_PATHWAY | 1.347879 | 0.10451555 | 0.3928318 | 0.98 |
| BIOCARTA_PML_PATHWAY | 1.3377569 | 0.11524693 | 0.39642933 | 0.94 |
| BIOCARTA_AMI_PATHWAY | 1.2983904 | 0.13252491 | 0.46804678 | 0.88 |
| BIOCARTA_CTCF_PATHWAY | 1.2880445 | 0.13023813 | 0.4731839 | 0.89 |
| BIOCARTA_ALK_PATHWAY | 1.283034 | 0.11520051 | 0.46607974 | 0.94 |
| BIOCARTA_DEATH_PATHWAY | 1.2741102 | 0.1327306 | 0.46858498 | 0.88 |
| BIOCARTA_VIP_PATHWAY | 1.2674688 | 0.14111607 | 0.4664202 | 0.85 |
| BIOCARTA_AKT_PATHWAY | 1.2580549 | 0.15756527 | 0.47109658 | 0.80 |
| BIOCARTA_EIF_PATHWAY | 1.2576509 | 0.17181334 | 0.45575204 | 0.76 |
| BIOCARTA_ERAD_PATHWAY | 1.2553302 | 0.16540907 | 0.44555822 | 0.78 |
| BIOCARTA_TNFR1_PATHWAY | 1.2484311 | 0.15225486 | 0.44573745 | 0.82 |
| BIOCARTA_P53_PATHWAY | 1.2253678 | 0.1963206 | 0.48195443 | 0.71 |
| BIOCARTA_NKCELLS_PATHWAY | 1.2020913 | 0.2062149 | 0.52011365 | 0.69 |
| BIOCARTA_KERATINOCYTE_PATHWAY | 1.1879789 | 0.18301101 | 0.5375874 | 0.74 |
| BIOCARTA_FMLP_PATHWAY | 1.1862072 | 0.19754252 | 0.5262874 | 0.70 |
| BIOCARTA_STATHMIN_PATHWAY | 1.164876 | 0.24508008 | 0.5614999 | 0.61 |
| BIOCARTA_IGF1MTOR_PATHWAY | 1.1634442 | 0.24514458 | 0.5496671 | 0.61 |
| BIOCARTA_CARDIACEGF_PATHWAY | 1.1595668 | 0.2547105 | 0.544126 | 0.59 |
| BIOCARTA_CDMAC_PATHWAY | 1.1580538 | 0.26100716 | 0.5335806 | 0.58 |
| BIOCARTA_SPRY_PATHWAY | 1.1108587 | 0.30970767 | 0.6309019 | 0.51 |
| BIOCARTA_NFKB_PATHWAY | 1.1096506 | 0.30381197 | 0.61843 | 0.52 |
| BIOCARTA_CHEMICAL_PATHWAY | 1.1074022 | 0.311513 | 0.6090333 | 0.51 |

| | | | | |
|---|---|---|---|---|
| BIOCARTA_TOB1_PATHWAY | 1.0882934 | 0.33364326 | 0.6406274 | 0.48 |
| BIOCARTA_RHO_PATHWAY | 1.0790172 | 0.3410946 | 0.6485116 | 0.47 |
| BIOCARTA_MCALPAIN_PATHWAY | 1.0677994 | 0.35990152 | 0.66100603 | 0.44 |
| BIOCARTA_INTRINSIC_PATHWAY | 1.0670543 | 0.3536968 | 0.64840794 | 0.45 |
| BIOCARTA_CYTOKINE_PATHWAY | 1.0626427 | 0.36824724 | 0.64497465 | 0.43 |
| BIOCARTA_MAL_PATHWAY | 1.0447583 | 0.3880034 | 0.67371356 | 0.41 |
| BIOCARTA_CALCINEURIN_PATHWAY | 1.0395671 | 0.3974903 | 0.67226565 | 0.40 |
| BIOCARTA_IL12_PATHWAY | 1.0345026 | 0.39895976 | 0.6707603 | 0.40 |
| BIOCARTA_IL1R_PATHWAY | 1.0253613 | 0.4075279 | 0.6789541 | 0.39 |
| BIOCARTA_TCR_PATHWAY | 1.0225375 | 0.4086661 | 0.67244893 | 0.39 |
| BIOCARTA_ARF_PATHWAY | 1.0184048 | 0.42733568 | 0.6692041 | 0.37 |
| BIOCARTA_NFAT_PATHWAY | 1.0114245 | 0.42573303 | 0.672665 | 0.37 |
| BIOCARTA_RAS_PATHWAY | 0.9913913 | 0.46243596 | 0.70591766 | 0.33 |
| BIOCARTA_RAC1_PATHWAY | 0.97415924 | 0.48812994 | 0.73297566 | 0.31 |
| BIOCARTA_ETS_PATHWAY | 0.959066 | 0.508913 | 0.7551856 | 0.29 |
| BIOCARTA_SPPA_PATHWAY | 0.9519709 | 0.51895386 | 0.7586634 | 0.28 |
| BIOCARTA_IGF1_PATHWAY | 0.9384135 | 0.5433746 | 0.7769473 | 0.26 |
| BIOCARTA_MEF2D_PATHWAY | 0.9355983 | 0.5468334 | 0.77040946 | 0.26 |
| BIOCARTA_LIS1_PATHWAY | 0.91710526 | 0.57502216 | 0.79978126 | 0.24 |
| BIOCARTA_EGF_PATHWAY | 0.91509145 | 0.58734375 | 0.7914895 | 0.23 |
| BIOCARTA_P53HYPOXIA_PATHWAY | 0.9058744 | 0.5990761 | 0.7994072 | 0.22 |
| BIOCARTA_LONGEVITY_PATHWAY | 0.89838165 | 0.60268766 | 0.80340034 | 0.22 |
| BIOCARTA_MTOR_PATHWAY | 0.89706117 | 0.6131761 | 0.79391605 | 0.21 |
| BIOCARTA_INTEGRIN_PATHWAY | 0.8948167 | 0.63394225 | 0.7866518 | 0.20 |
| BIOCARTA_GCR_PATHWAY | 0.85959905 | 0.6648847 | 0.84936625 | 0.18 |
| BIOCARTA_TFF_PATHWAY | 0.8476242 | 0.7020563 | 0.8617997 | 0.15 |
| BIOCARTA_BCR_PATHWAY | 0.84715205 | 0.723042 | 0.8502531 | 0.14 |
| BIOCARTA_PAR1_PATHWAY | 0.8460295 | 0.6891448 | 0.84036493 | 0.16 |
| BIOCARTA_PPARA_PATHWAY | 0.84454286 | 0.76018596 | 0.83146054 | 0.12 |
| BIOCARTA_P38MAPK_PATHWAY | 0.8395507 | 0.7427122 | 0.8296007 | 0.13 |
| BIOCARTA_GPCR_PATHWAY | 0.8289004 | 0.7488292 | 0.838532 | 0.13 |
| BIOCARTA_CCR3_PATHWAY | 0.8148836 | 0.7393919 | 0.85308176 | 0.13 |
| BIOCARTA_HCMV_PATHWAY | 0.7631095 | 0.81239027 | 0.92986834 | 0.09 |
| BIOCARTA_TALL1_PATHWAY | 0.75543344 | 0.8116551 | 0.9292582 | 0.09 |
| BIOCARTA_TH1TH2_PATHWAY | 0.7362943 | 0.87137955 | 0.9445508 | 0.06 |
| BIOCARTA_VEGF_PATHWAY | 0.7277883 | 0.89335394 | 0.943691 | 0.05 |
| BIOCARTA_CARM_ER_PATHWAY | 0.7218952 | 0.896598 | 0.93916774 | 0.05 |
| BIOCARTA_HIF_PATHWAY | 0.692204 | 0.89008534 | 0.96051615 | 0.05 |
| BIOCARTA_DC_PATHWAY | 0.68051153 | 0.9110301 | 0.959867 | 0.04 |
| BIOCARTA_TGFB_PATHWAY | 0.6781106 | 0.91836494 | 0.9503005 | 0.04 |
| BIOCARTA_ARENRF2_PATHWAY | 0.6558775 | 0.93987936 | 0.9566516 | 0.03 |
| BIOCARTA_BCELLSURVIVAL_PATHWAY | 0.6399668 | 0.93809664 | 0.95585734 | 0.03 |
| BIOCARTA_PDZS_PATHWAY | -1.6436162 | 0.01594262 | 0.74460626 | -1.80 |
| BIOCARTA_BAD_PATHWAY | -1.4724541 | 0.0425139 | 1 | -1.37 |
| BIOCARTA_CREB_PATHWAY | -1.3831431 | 0.08020075 | 1 | -1.10 |
| BIOCARTA_IGF1R_PATHWAY | -1.3691378 | 0.08513583 | 1 | -1.07 |

| | | | | |
|---|---|---|---|---|
| BIOCARTA_EICOSANOID_PATHWAY | -1.268871 | 0.1496557 | 1 | -0.82 |
| BIOCARTA_BARRESTIN_SRC_PATHWAY | -1.1952072 | 0.22363271 | 1 | -0.65 |
| BIOCARTA_NO1_PATHWAY | -1.1557053 | 0.23832116 | 1 | -0.62 |
| BIOCARTA_NOS1_PATHWAY | -1.1424361 | 0.26863128 | 1 | -0.57 |
| BIOCARTA_AGR_PATHWAY | -1.1418123 | 0.24896383 | 1 | -0.60 |
| BIOCARTA_PYK2_PATHWAY | -1.1405711 | 0.2573413 | 1 | -0.59 |
| BIOCARTA_GH_PATHWAY | -1.1346002 | 0.26595184 | 1 | -0.58 |
| BIOCARTA_IL2_PATHWAY | -1.118907 | 0.2906249 | 1 | -0.54 |
| BIOCARTA_CSK_PATHWAY | -1.1043952 | 0.3062085 | 1 | -0.51 |
| BIOCARTA_PTDINS_PATHWAY | -1.0823604 | 0.33151498 | 1 | -0.48 |
| BIOCARTA_EPO_PATHWAY | -1.0810517 | 0.3416722 | 1 | -0.47 |
| BIOCARTA_IL7_PATHWAY | -1.0604036 | 0.36941144 | 1 | -0.43 |
| BIOCARTA_CTLA4_PATHWAY | -1.0563803 | 0.36639023 | 1 | -0.44 |
| BIOCARTA_CERAMIDE_PATHWAY | -1.0557604 | 0.3697992 | 1 | -0.43 |
| BIOCARTA_MPR_PATHWAY | -1.0409753 | 0.38825387 | 1 | -0.41 |
| BIOCARTA_IL2RB_PATHWAY | -1.0380114 | 0.3834126 | 1 | -0.42 |
| BIOCARTA_INFLAM_PATHWAY | -1.0362148 | 0.39471075 | 1 | -0.40 |
| BIOCARTA_IL3_PATHWAY | -1.0133523 | 0.42958453 | 1 | -0.37 |
| BIOCARTA_CK1_PATHWAY | -0.9977731 | 0.4543368 | 1 | -0.34 |
| BIOCARTA_HER2_PATHWAY | -0.9756703 | 0.48728025 | 1 | -0.31 |
| BIOCARTA_ARAP_PATHWAY | -0.9549202 | 0.51437217 | 1 | -0.29 |
| BIOCARTA_GLEEVEC_PATHWAY | -0.9394226 | 0.5441683 | 1 | -0.26 |
| BIOCARTA_CCR5_PATHWAY | -0.9378719 | 0.53882855 | 1 | -0.27 |
| BIOCARTA_CHREBP_PATHWAY | -0.9326578 | 0.5474827 | 1 | -0.26 |
| BIOCARTA_AT1R_PATHWAY | -0.9308376 | 0.5602626 | 1 | -0.25 |
| BIOCARTA_EDG1_PATHWAY | -0.9184533 | 0.57482415 | 1 | -0.24 |
| BIOCARTA_TPO_PATHWAY | -0.8959867 | 0.61872894 | 1 | -0.21 |
| BIOCARTA_MET_PATHWAY | -0.8952578 | 0.6347447 | 1 | -0.20 |
| BIOCARTA_MAPK_PATHWAY | -0.8911594 | 0.7018361 | 1 | -0.15 |
| BIOCARTA_HDAC_PATHWAY | -0.8781146 | 0.6493985 | 1 | -0.19 |
| BIOCARTA_COMP_PATHWAY | -0.8759257 | 0.6439494 | 1 | -0.19 |
| BIOCARTA_PTEN_PATHWAY | -0.8523405 | 0.67915314 | 1 | -0.17 |
| BIOCARTA_NGF_PATHWAY | -0.847661 | 0.69049686 | 1 | -0.16 |
| BIOCARTA_MTA3_PATHWAY | -0.8460824 | 0.6849484 | 1 | -0.16 |
| BIOCARTA_CXCR4_PATHWAY | -0.8433474 | 0.6971057 | 1 | -0.16 |
| BIOCARTA_BIOPEPTIDES_PATHWAY | -0.8382102 | 0.73102087 | 1 | -0.14 |
| BIOCARTA_PDGF_PATHWAY | -0.8350279 | 0.73501784 | 1 | -0.13 |
| BIOCARTA_CLASSIC_PATHWAY | -0.8298029 | 0.7026356 | 1 | -0.15 |
| BIOCARTA_NDKDYNAMIN_PATHWAY | -0.814623 | 0.7343068 | 1 | -0.13 |
| BIOCARTA_SHH_PATHWAY | -0.8132258 | 0.73469883 | 1 | -0.13 |
| BIOCARTA_ERK_PATHWAY | -0.8110458 | 0.7721108 | 0.9941731 | -0.11 |
| BIOCARTA_PITX2_PATHWAY | -0.7987005 | 0.7576211 | 0.99767715 | -0.12 |
| BIOCARTA_NKT_PATHWAY | -0.7984607 | 0.78173447 | 0.97691625 | -0.11 |
| BIOCARTA_EIF4_PATHWAY | -0.7735193 | 0.8255202 | 1 | -0.08 |
| BIOCARTA_GSK3_PATHWAY | -0.7711451 | 0.81183255 | 0.9867842 | -0.09 |
| BIOCARTA_WNT_PATHWAY | -0.7403485 | 0.8688135 | 1 | -0.06 |

| | | | | |
|---|---|---|---|---|
| BIOCARTA_IL6_PATHWAY | -0.7192193 | 0.8854084 | 1 | -0.05 |
| BIOCARTA_FCER1_PATHWAY | -0.7125605 | 0.9376057 | 1 | -0.03 |
| BIOCARTA_INSULIN_PATHWAY | -0.6874885 | 0.91713446 | 1 | -0.04 |
| BIOCARTA_NUCLEARRS_PATHWAY | -0.6851767 | 0.950931 | 1 | -0.02 |
| BIOCARTA_NO2IL12_PATHWAY | -0.6598453 | 0.92338455 | 1 | -0.03 |
| BIOCARTA_ASBCELL_PATHWAY | -0.6509321 | 0.9290697 | 1 | -0.03 |
| BIOCARTA_LAIR_PATHWAY | -0.6115861 | 0.96031505 | 1 | -0.02 |
| BIOCARTA_AMAN_PATHWAY | -0.6016997 | 0.9637118 | 0.9924618 | -0.02 |
| BIOCARTA_PGC1A_PATHWAY | -0.5551261 | 0.9826503 | 0.9894193 | -0.01 |

#### Table S4 Combination index values for ALT neuroblastoma cell lines treated with TMZ+SN-38+AZD0156

##### ALT + NB Cell Lines

| CHLA-90 | Combination Index value |
|---|---|
| AZD0156 (100nM)+TMZ (50uM)+SN38(5nM) | 0.429 |
| AZD0156 (50nM)+TMZ (25)+SN38(2.5nM) | 0.369 |
| AZD0156 (25nM)+TMZ (12.5uM)+SN38(1.25nM) | 0.275 |
| AZD0156 (12.5nM)+TMZ (6.25uM)+SN38(0.625nM) | 0.359 |

| SK-N-FI | Combination Index value |
|---|---|
| AZD0156 (100nM)+TMZ (50uM)+SN38(5nM) | 0.125 |
| AZD0156 (50nM)+TMZ (25)+SN38(2.5nM) | 0.265 |
| AZD0156 (25nM)+TMZ (12.5uM)+SN38(1.25nM) | 0.379 |
| AZD0156 (12.5nM)+TMZ (6.25uM)+SN38(0.625nM) | 1.104 |

| COG-N-515 | Combination Index value |
|---|---|
| AZD0156 (100nM)+TMZ (50uM)+SN38(5nM) | 0.321 |
| AZD0156 (50nM)+TMZ (25)+SN38(2.5nM) | 0.295 |
| AZD0156 (25nM)+TMZ (12.5uM)+SN38(1.25nM) | 0.476 |
| AZD0156 (12.5nM)+TMZ (6.25uM)+SN38(0.625nM) | 0.765 |

| COG-N-512 | Combination Index value |
|---|---|
| AZD0156 (100nM)+TMZ (50uM)+SN38(5nM) | 0.602 |
| AZD0156 (50nM)+TMZ (25)+SN38(2.5nM) | 0.452 |
| AZD0156 (25nM)+TMZ (12.5uM)+SN38(1.25nM) | 0.493 |
| AZD0156 (12.5nM)+TMZ (6.25uM)+SN38(0.625nM) | 0.533 |

### Table S5 Pharmacokinetics of AZD0156 in swiss athymic nu/nu male mice

AZD0156 5 mg/kg unbound (free) concentration (nM)

| (h) | AZD0156 @ 5 mg/kg | ± Error | Animal 1 | Animal 2 | Animal 3 |
|---|---|---|---|---|---|
| 0.5 | 40.2840 | 10.5801 | 41.472 | 50.22 | 29.16 |
| 2.0 | 51.0840 | 1.6861 | 50.544 | 49.734 | 52.974 |
| 6.0 | 24.7860 | 6.5264 | 22.032 | 32.238 | 20.088 |
| 12.0 | 14.1480 | 1.6227 | 15.714 | 12.474 | 14.256 |
| 24.0 | 5.8320 | 1.0117 | 6.966 | 5.508 | 5.022 |

| | | |
|---|---|---|
| AUCinf | 540.422676 | h.nM |
| AUC0-t | 467.019 | h.nM |
| Cmax/C0 | 51.084 | nM |
| Tmax | 2 | h |

Assuming dose proportionality

AZD0156 20 mg/kg unbound (free) concentration (nM)

| (h) | AZD0156 @20mg/kg | ± Error | Animal 1 | Animal 2 | Animal 3 |
|---|---|---|---|---|---|
| 0.5 | 161.1360 | 42.3206 | 165.8880 | 200.8800 | 116.6400 |
| 2.0 | 204.3360 | 6.7446 | 202.1760 | 198.9360 | 211.8960 |
| 6.0 | 99.1440 | 26.1056 | 88.1280 | 128.9520 | 80.3520 |
| 12.0 | 56.5920 | 6.4908 | 62.8560 | 49.8960 | 57.0240 |
| 24.0 | 23.3280 | 4.0468 | 27.8640 | 22.0320 | 20.0880 |

| | | |
|---|---|---|
| AUCinf | 2161.690704 | h.nM |
| AUC0-t | 1868.076 | h.nM |
| Cmax/C0 | 204.336 | nM |
| Tmax | 2 | h |
